# Integrating Microbial Ecology and Sustainability: Linking Agricultural Waste Gas Streams to Microbial Community Shifts in a Photobioreactor

**DOI:** 10.64898/2026.11.06.764810

**Authors:** Marta Velaz Martín, Till Glockow, Laura Meisch, John Vollmers, Anne-Kristin Kaster, Kersten S. Rabe, Christof M. Niemeyer

## Abstract

Photobioreactors (PBRs) coupled to animal housing offer potential for mitigating gaseous emissions such as CO2 and NH3 while simultaneously producing biomass. Here, we present a time-resolved characterization of microbial community dynamics, with a focus on the genetic potential for carbon and nitrogen metabolism. Using a cone-shaped, helical PBR cultivating *Arthrospira platensis* (Spirulina) and supplied with exhaust air from a chicken house, we characterized microbial communities over continuous exhaust-air exposure and periodic biomass harvesting. Microbiome and gene-centric profiling revealed temporal variation in community composition and in the representation of genes associated with carbon and nitrogen metabolism. Cycle-based dissolved inorganic nitrogen balances showed that the decrease in dissolved inorganic nitrogen was only partially recovered in harvested biomass, while ammonia-oxidizing microorganisms were associated with structured, low-light niches. These observations indicate ammonia-oxidizing potential but do not establish a quantitatively relevant nitrification flux. Structured carrier matrices provided attached-growth niches harboring microbial populations underrepresented in the circulating suspension, while pre-colonized carriers facilitated biomass development during re-inoculation. Overall, our results show that coupling PBRs to agricultural exhaust streams can support biomass production while sustaining spatially structured microbial communities with taxonomic and functional potential. These findings may inform their further development for emission mitigation and circular biomass production.

**Short Synopsis Statement:** Photobioreactors coupled to agricultural exhaust streams offer potential for linking emission mitigation with biomass production, while the underlying nitrogen transformations remain poorly understood. We show that the decrease in measured dissolved inorganic nitrogen is only partially recovered in biomass and that ammonia-oxidizing microorganisms preferentially associate with structured, low-light niches, while their quantitative contribution to nitrogen transformation remains unresolved.

**Table of Contents Graphic:** 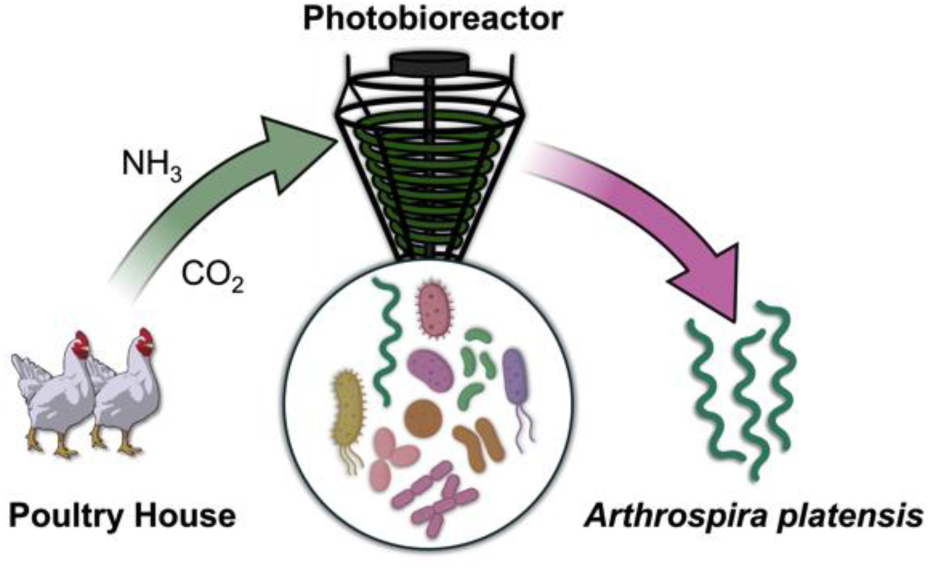

## 1. Introduction

In the face of the climate crisis, increasing water scarcity, and growing demands for efficient agricultural production, closed-loop approaches are becoming increasingly important in agriculture.^1^ This is particularly evident in controlled environment agriculture (CEA)^2–5^, where resource use can be optimized through recycling of nutrients, substrates, and water. Microbial communities play a central role in such systems by driving nutrient turnover and stabilizing biological processes.^6^

Building on this concept, we recently introduced a circular approach linking animal housing with biomass production using photosynthetic microorganisms (PMOs).^7^ These organisms can convert animal-exhaled CO_2_ into valuable compounds and are also capable of removing nitrogen, sulfur, and other gaseous or aerosol-bound substances from the air.^8^ Previous studies demonstrated efficient air purification and biomass production, highlighting the potential of such systems to convert livestock emissions into value-added biomass within circular agricultural concepts, and provided first insights into biomass composition and microbial community dynamics.^7, 9, 10^

Despite these advances, the underlying microbial processes governing system performance remain insufficiently resolved. In particular, microbial analyses in such systems are often descriptive and focus on community composition, while process-level understanding of nutrient transformation pathways is limited. Furthermore, photobioreactors represent spatially heterogeneous environments, where gradients in light and physicochemical conditions may give rise to distinct microbial niches that are not captured by bulk analyses.

In mixed microalgal-bacterial systems, NH_3_/NH_4_^+^ can be assimilated into phototrophic biomass or serve as substrate for ammonia-oxidizing microorganisms.^11, 12^ Nitrifiers can be inhibited by high irradiance in algal-bacterial photobioreactors,^13, 14^ whereas attached-growth environments can provide protected niches that support nitrifying and other surface-associated microbial populations.^12, 15–17^ These observations motivated the inclusion of carrier materials in this study to capture microbial populations potentially underrepresented by bulk suspension sampling.

Here, we investigate microbial community development in an operational photobioreactor cultivating *Arthrospira platensis* (*Spirulina*) under continuous exposure to poultry-house exhaust air. Reactor operation and biomass development were quantitatively characterized to provide the process context for interpreting these community patterns. Structured carriers were used to examine surface-associated communities, while a separate complementary experiment using air-scrubber process water examined changes in dissolved nitrogen species and cycle-based nitrogen balances. Ammonia-oxidizing microorganisms were preferentially associated with structured, low-light niches; however, their quantitative contribution to nitrogen conversion was not assessed.

Building on previous studies that established the photobioreactor concept and its application to *Arthrospira* cultivation using poultry-house exhaust air,^7, 9, 10^ the present study focuses on the temporal development of the microbial community and its spatial partitioning between the bulk suspension and structured, low-light niches.

## 2. Materials and Methods

### 2.1. Microalgae cultivation and sample preparation

A mixed culture of *Arthrospira platensis* (*A. platensis*) originating from an open pond (ROVAL GmbH, Germany) was cultivated in a single closed photobioreactor (∼150 L), as reported previously.^7^ The experiment comprised one reactor run without independent biological reactor replicates; samples collected over time therefore represent repeated temporal samples from the same experimental system. The culture was intentionally maintained under non-axenic conditions and represents an *Arthrospira*-dominated microbial community rather than a monoculture. Detailed cultivation conditions and medium composition are provided in SI Section S1.

The culture was first propagated in 10-L algal bags before transfer to the photobioreactor. Following transfer, the reactor was operated for 7 days with continuous ambient-air aeration (15 L min^-1^; ∼400 ppm CO_2_; NH_3_ not detectable) to allow acclimation to the airlift cultivation conditions. On day 0, the gas source was switched from ambient air to poultry-house exhaust air, which was supplied continuously until the end of the experiment. Approximately 1 m^3^ h^-1^ of poultry-house exhaust air was directed through the reactor, corresponding to ∼0.6% of the total poultry-house ventilation flow. The reactor therefore represented an experimental side-stream configuration rather than a system dimensioned for treatment of the complete poultry-house ventilation stream.

The photobioreactor included a connected reservoir compartment (∼30-40 L) providing additional low-light and surface-associated microenvironments. The system was operated semi-continuously, with biomass harvested on days 5, 10, 13, 18, and 21 after the switch to poultry-house exhaust air. Harvesting was performed by rotational filtration (30 µm), with the clarified medium re-fertilized and returned to the reactor (SI Section S1). Because harvesting did not involve proportional replacement of the reactor liquid volume, a conventional hydraulic retention time (HRT) based on continuous influent and effluent flow cannot be assigned. Similarly, a single solids retention time (SRT) cannot be assigned to the complete microbial community because filtration selectively concentrated filamentous *Arthrospira*, as reflected by the >96% *Cyanobacteriota* abundance in the harvested fraction (Figure 4). Biomass concentration was estimated from OD_565_ using the experimentally validated relationship with gravimetric dry biomass concentration (R² = 0.9973; Supplementary Figure S1); individual harvesting events reduced suspended biomass concentration by approximately 36-76% (Supplementary Figure S2).

Samples of the reactor suspension were collected over 33 days and assigned to three temporal sampling windows relative to the switch from ambient air to poultry-house exhaust air: before exhaust-air exposure (B; n = 5), initial exhaust-air exposure (I; days 1-10, n = 5), and later exhaust-air exposure (L; days 14-25, n = 8) (Figure 1B). These categories represent temporal sampling windows rather than distinct aeration regimes; the reactor remained continuously aerated throughout the experiment. The initial and later windows additionally differ in elapsed time and cumulative reactor operation, including harvesting and re-fertilization events; consequently, differences between them cannot be attributed specifically to increasing duration of exhaust-air exposure.

**Figure 1:**
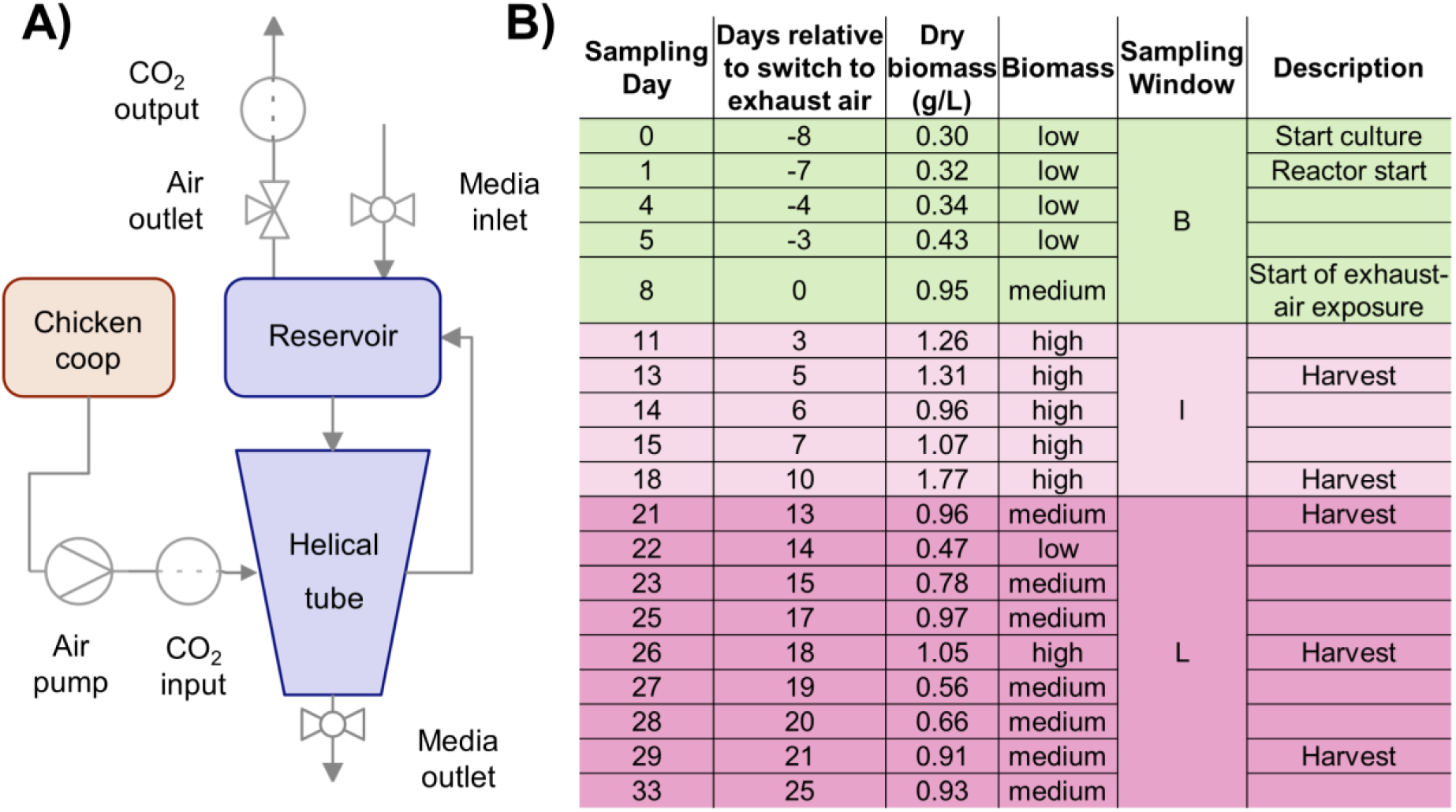
Experimental setup. (A) The photobioreactor was inoculated with *A. platensis* pre-cultivated in 10-L algal bags and operated for 7 days in airlift mode using ambient air (∼400 ppm CO_2_; NH_3_ not detectable) to allow acclimation to the reactor conditions. On day 0, the gas supply was switched to poultry-house exhaust air and maintained continuously until the end of the experiment. (B) Suspension samples were collected at defined time points and grouped into three temporal sampling windows: before exhaust-air exposure (B, green), initial sampling window under continuous exhaust-air exposure (I, light pink), and later sampling window under continuous exhaust-air exposure (L, dark pink). Biomass was periodically harvested on days 5, 10, 13, 18, and 21 relative to the switch to poultry-house exhaust air.

Suspension samples were obtained directly from the circulating culture, whereas harvested biomass samples correspond to the concentrated biomass fraction obtained during routine rotational filtration. Harvesting was restricted to the suspended culture; carrier-associated biomass was not removed or included in the harvested fraction. Further details on cultivation, poultry housing, gas composition and monitoring, sampling, and sample processing are provided in SI Section S1.

### 2.2. Nitrogen dynamics and biomass-associated nitrogen

To complement the poultry-exhaust-air experiment, nitrogen dynamics were investigated in a separate experiment using process water from an industrial-scale air scrubber treating livestock exhaust air. Air scrubbing concentrates nitrogen from a large volume of dilute exhaust into a smaller aqueous stream, while absorbed NH_3_ may additionally be biologically oxidized, resulting in nitrate accumulation in the scrubbing liquid.^18^ The process water was therefore considered an agricultural emission-derived but already processed nitrogen stream rather than a proxy for untreated poultry-house exhaust. Its use enabled controlled dosing and monitoring of dissolved inorganic nitrogen independently of short-term variability in the gas-phase input.

Nitrogen dynamics were evaluated based on dissolved inorganic nitrogen (DIN; NH ^+^ + NO ^-^ + NO ^-^) and the individual inorganic nitrogen species. The system was operated under semi-continuous conditions, and changes in DIN were evaluated from time-resolved concentration profiles and cycle-based balances. Biomass-associated nitrogen (N_biomass) was estimated from OD_565_-derived biomass production (Section 2.1) and the nitrogen content of *Arthrospira* estimated from the measured protein content. Nitrogen closure was defined as the fraction of the decrease in measured DIN recovered in biomass. The fate of the remaining nitrogen cannot be resolved from the present measurements and may include unmeasured nitrogen pools and gaseous losses, including NH_3_ volatilization under the alkaline and continuously aerated cultivation conditions. Further analytical details, calculations, and assumptions are provided in SI Section S2.

### 2.3. Cultivation in matrix materials

To investigate microbial community development in localized attached-growth niches, inert carrier matrices were introduced into the photobioreactor. Macroporous elastomeric silicone foams (MESIF) and commercial biofilm carriers (MUTAG™ BioChip) were used as standardized colonization substrates. The carriers provided defined, retrievable surfaces for examining microbial populations that preferentially established under attached conditions relative to the circulating suspension. They were confined to the reservoir compartment without direct artificial illumination; thus, surface attachment and reduced light availability co-varied in this experimental design. The carriers were not intended or evaluated as a process-scale biofilm module, and the quantitative contribution of carrier-associated communities to whole-reactor performance was not assessed.

MESIF materials were prepared as described previously,^19^ and detailed preparation protocols are provided in SI Section S3. MUTAG™ carriers were used according to manufacturer specifications. For each experiment, approximately 100 MESIF pieces and 100 MUTAG™ carriers (each approximately 2 × 2 cm) were placed in a coarse-mesh net within the central reservoir compartment of the reactor, where they remained spatially confined while being continuously exposed to the circulating culture medium. Unlike the suspension circulating through the artificially illuminated helical tube, the carriers were confined to the reservoir compartment without direct artificial illumination. Although some incidental ambient light may have reached the reservoir, light intensity within this compartment was not measured. At defined sampling time points (Figure 1), individual carriers (n = 3 per material) were retrieved from the net and immediately stored at −20 °C for downstream analysis. The total carrier volume was approximately 0.2 L, corresponding to ∼0.13% of the ∼150 L reactor volume.

To assess whether matrix-associated microbial communities contribute to culture establishment and biomass development, re-inoculation experiments were performed using MESIF carrier matrices. Pre-colonized MESIFs obtained from previous photobioreactor operation were introduced into newly established cultures, while non-colonized MESIFs served as controls.

To evaluate the generalizability of the observed effects, a different laboratory strain (*Arthrospira maxima*) was used. Cultures were established under identical conditions and aerated with exhaust air. Biomass development was monitored over time using absorbance measurements as a proxy for growth. Further details are provided in SI Section S3.

### 2.4. DNA extraction and sequencing

Genomic DNA was extracted from both suspension samples and matrix-associated biomass using a commercial soil DNA extraction kit following the manufacturer’s protocol. Microbial community composition was analyzed using 16S rRNA gene amplicon sequencing targeting the V3 region. In addition, shotgun metagenomic sequencing was performed for selected samples to obtain taxonomic and functional insights. Metagenomic data were further used to identify both prokaryotic and eukaryotic taxa based on small- and large-subunit rRNA gene fragments predicted from assembled metagenomes and classified against the SILVA 138.2 database. The combination of both approaches enabled high-resolution taxonomic profiling alongside functional characterization of the microbial community. Detailed descriptions of DNA extraction, sequencing protocols, and bioinformatic analyses are provided in SI Section S4.

### 2.5. Data Availability

Raw sequence data are available in the NCBI Sequence Read Archive (SRA) under accession PRJNA1330278. Additional datasets can be obtained from the corresponding author upon reasonable request.

## 3. Results and Discussion

### 3.1. Cultivation, sampling and physico-chemical analysis

To investigate microbiome dynamics in a photobioreactor supplied with poultry exhaust air, we used a previously established cultivation system in which a mixed culture of *Arthrospira platensis* is grown in a closed photobioreactor.^7^ The inoculum was first propagated in 10-L algal bags and subsequently transferred to the ∼150-L photobioreactor. During a 7-day acclimation period, the reactor was operated in airlift mode with ambient air (∼400 ppm CO_2_; NH_3_ not detectable). On day 0, the gas supply was switched to poultry-house exhaust air and maintained continuously until the end of the experiment. To characterize microbial communities beyond the circulating suspension, MESIF and MUTAG™ carriers were introduced at low loading as standardized colonization substrates providing localized attached-growth niches within the reservoir compartment. Suspension samples (n = 18) were analyzed by amplicon and shotgun metagenomic sequencing; MESIF and MUTAG™ samples (n = 12 each) were analyzed by amplicon sequencing, with three samples per carrier type additionally subjected to shotgun metagenomics.

Under continuous poultry-exhaust-air exposure, the culture maintained repeated biomass accumulation between harvesting events throughout the experimental period (Figures 2 and S2). Based on the increase in OD_565_-derived dry biomass concentration between successive harvesting events (Supplementary Figure S2), apparent volumetric biomass productivity ranged from 0.072 to 0.186 g L^-1^ d^-1^, with an average of 0.134 g L^-1^ d^-1^ over days 0-25, corresponding to an apparent biomass accumulation of approximately 20 g d^-1^ for the 150 L reactor.

**Figure 2:**
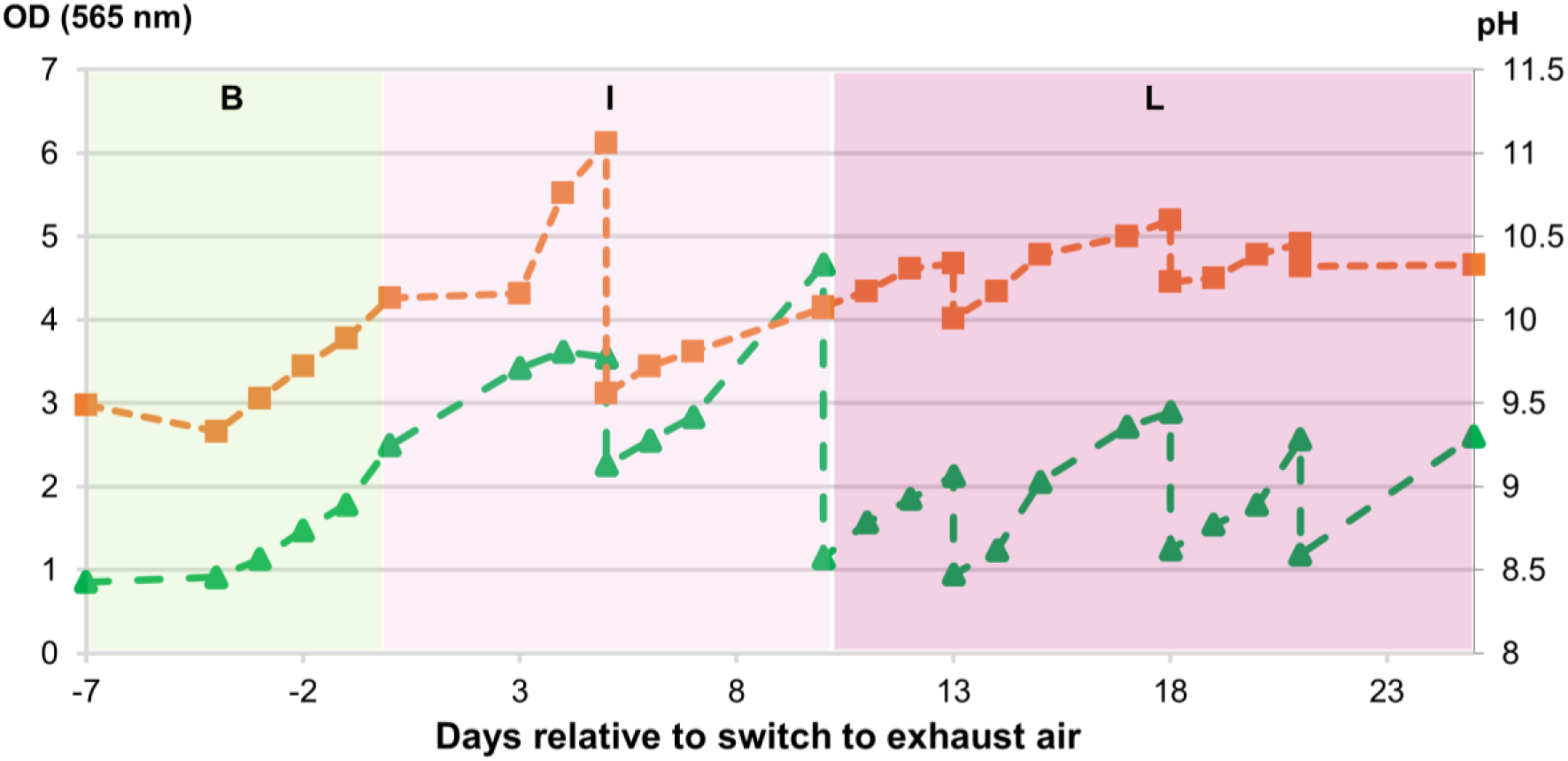
Cultivation of A. platensis under aeration with poultry exhaust air. Optical density at 565 nm (OD₅₆₅, green, left axis) and pH (orange, right axis) are shown over the course of the experiment. Aeration with exhaust air was initiated after a 7-day acclimation period (day 0) and maintained continuously for 25 days (see Figure S2 for detailed temporal profiles). OD_565_ was used to estimate biomass concentration based on its experimentally validated relationship with gravimetric dry biomass; an OD₅₆₅ of 1 corresponds to approximately 400 mg L^-1^ dry biomass of *A. platensis* under the conditions of this system (Figure S1). Shaded areas indicate the temporal sampling windows B (green), I (light pink), and L (dark pink), as defined in Figure 1.

### 3.2. Shifts in microbial diversity and enrichment of taxa in 16S rRNA amplicons

To characterize microbial community dynamics during reactor operation, 16S rRNA amplicon sequencing was performed on suspension samples collected across three sampling windows (before, initial, and later; Figure 1B). Because exhaust-air aeration was maintained continuously from day 0 onwards, these windows represent successive stages of reactor operation rather than distinct exhaust-air treatments. They therefore differ not only in duration of exhaust-air exposure, but also in elapsed time and the cumulative number of harvesting and re-fertilization events. Accordingly, the observed community differences are interpreted as temporal dynamics under continuous reactor operation rather than as effects attributable to exhaust-air alone. Changes in alpha and beta diversity were evaluated to characterize overall community restructuring (Supplementary Figure S3).

Alpha diversity showed a decreasing trend over the course of reactor operation, although no significant overall differences were detected for Shannon diversity (p = 0.25), ACE (p = 0.45), or Faith’s phylogenetic diversity (p = 0.051; Supplementary Figure S3A-C). Beta-diversity ordinations suggested temporal restructuring, with later samples showing greater separation from earlier sampling windows. For Jaccard distances, sampling window was associated with 27.7% of the observed compositional variation (PERMANOVA: R² = 0.277, p < 0.001), although dispersion also differed significantly (PERMDISP: p = 0.024), requiring cautious interpretation. Weighted UniFrac showed no significant difference among sampling windows (PERMANOVA: R² = 0.171, p = 0.209; Supplementary Figure S3D-E).

To further resolve taxonomic differences among sampling windows, differential abundance analysis was performed using ANCOM-BC^20, 21^. In total, 543 taxa were identified as differentially abundant among sampling windows (Supplementary Table S2), with selected strongly enriched taxa shown in Figure 3.

**Figure 3:**
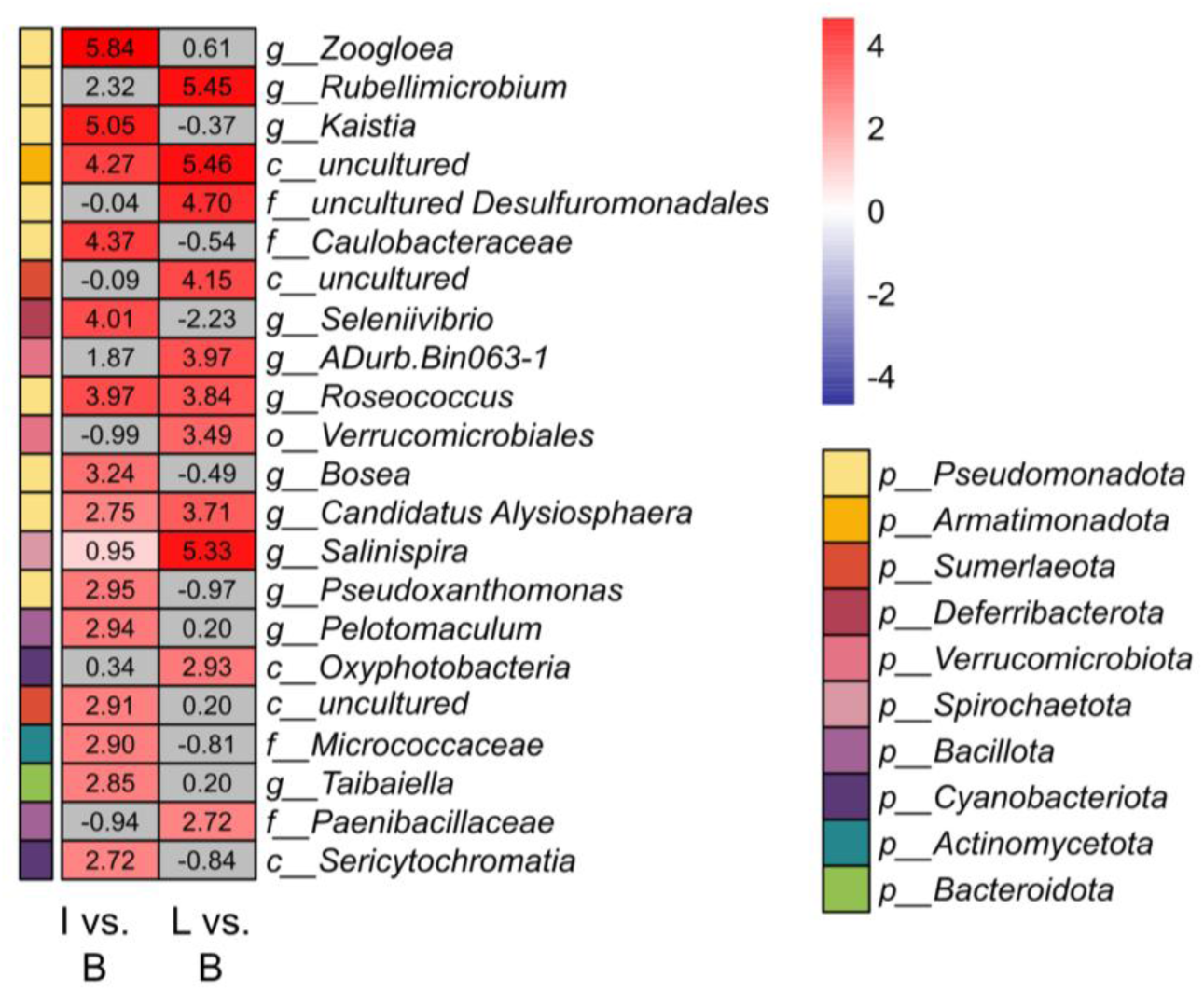
Differentially abundant taxa were identified using Analysis of Compositions of Microbiomes with Bias Correction (ANCOM-BC). Log-fold changes were calculated at the genus level for the sampling-window comparisons I vs B (initial vs before) and L vs B (later vs before). The initial window comprised days 1-10 and the later window days 14-25 under continuous poultry-exhaust exposure, while the before window comprised samples collected during ambient-air acclimation and prior to poultry-exhaust exposure (days −7 to 0). These labels therefore denote temporal sampling windows within a single continuously operated reactor campaign rather than distinct aeration treatments. Phyla are indicated by colored boxes to the left of the data columns. Where a taxon could not be resolved to genus level, the lowest assigned taxonomic rank is shown using the prefixes g for genus, f for family, o for order, and c for class. Taxa with significant differential abundance (p < 0.05) are shown in red, whereas non-significant taxa are shown in grey.

During the initial aeration phase, several genera associated with heterotrophic carbon turnover and nitrogen transformations were enriched, including *Zoogloea*^22, 23^, *Kaistia*^24, 25^, and *Bosea*^26–28^. These taxa are linked to processes such as aerobic carbon oxidation, nitrate reduction, and denitrification. In addition, *Seleniivibrio*^29^ was also enriched. Members of this genus have been reported to respire selenate and arsenate,^29^ but neither these nor other alternative electron acceptors were measured here, and the enrichment is therefore reported descriptively without a functional inference.

In the later aeration phase, the community shifted toward taxa associated with autotrophic and phototrophic processes. Notably, *Rubellimicrobium*^30, 31^ and unclassified Oxophotobacteria^32, 33^ were enriched, indicating increased contributions of oxygenic photosynthesis and CO_2_ fixation.

Several taxa persisted across both phases, including *Roseococcus*^34^ and *Candidatus Alysiosphaera*^35^, suggesting metabolic flexibility under changing conditions. In contrast, *Salinispira* showed stronger enrichment during the later sampling window. Members of this genus have been described as anaerobic, halophilic fermenters capable of utilizing carbohydrates.^36, 37^ Such a physiology could be compatible with organic substrates derived from algal biomass in the reactor; however, dissolved organic carbon was not measured, and the trophic basis of the observed enrichment therefore remains unresolved.

In addition to these characterized taxa, a substantial fraction of enriched lineages remained taxonomically unresolved, including members of *Pseudomonadota*, *Desulfobacterota*, and *Sumerlaeota*, highlighting the potential role of underexplored microbial groups in carbon and nitrogen cycling within the system.

Together, these results indicate temporal restructuring of the microbial community during reactor operation, with enrichment patterns differing between the initial and later sampling windows, despite no significant overall differences in alpha diversity or abundance-weighted phylogenetic composition. However, because exhaust-air exposure increased concurrently with elapsed time and repeated harvesting and re-fertilization events, the individual contribution of these factors cannot be disentangled within the present experimental design.

### 3.3. Metagenomic insights into taxonomic composition

To assess the taxonomic composition of the microbial community, metagenomic sequencing reads were assembled and ribosomal RNA genes were identified and classified using the SILVA 138.1 database, enabling taxonomic assignment across both prokaryotic and eukaryotic domains (Figure 4).

**Figure 4:**
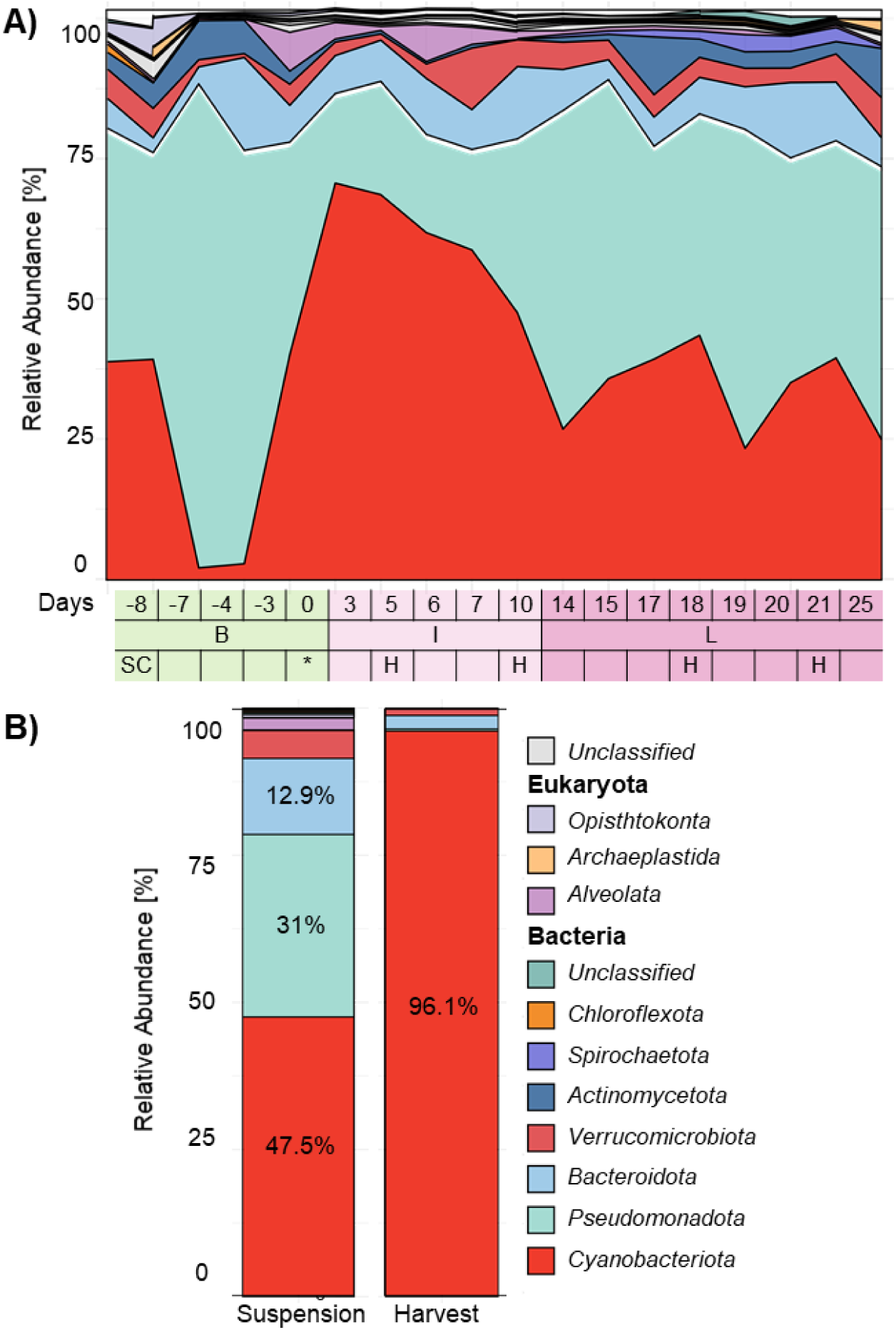
Metagenomic taxonomic composition of the photobioreactor community. (A) Relative abundance of the major bacterial phyla across all sampling time points, grouped into the temporal sampling windows before exhaust-air exposure (B, green), initial sampling window under continuous exhaust-air exposure (I, light pink), and later sampling window under continuous exhaust-air exposure (L, dark pink). Numbers on the x-axis indicate sampling days relative to the onset of exhaust-air exposure. Aeration with poultry-house exhaust air was initiated on day 0 (*) and maintained continuously thereafter; biomass harvesting events are indicated by H. (B) Comparison of the phylum-level composition of the reactor suspension and the harvested biomass fraction collected on day 10. Figure 4 presents taxonomic composition at the phylum level; the genus Arthrospira is therefore represented within *Cyanobacteriota*. Genus-level taxonomic profiles, including *Arthrospira*, are presented in Supplementary Figure S4.

At the phylum level, the community showed temporal restructuring during reactor operation. Early samples displayed transient fluctuations, including a temporary increase in *Pseudomonadota*, whereas later samples were increasingly dominated by *Cyanobacteriota* (Figure 4A).

At genus level, *Arthrospira* accounted for up to 60% of the total community in suspension samples (Supplementary Figure S4), indicating the successful persistence and proliferation of the target phototrophic organism.

Biomass harvesting events were accompanied by transient decreases in the relative abundance of *Cyanobacteriota* in the suspension, consistent with periodic removal of *Arthrospira*-rich biomass. The harvested fraction collected on day 10 contained more than 96% *Cyanobacteriota* (Figure 4B), consistent with preferential retention of filamentous *Arthrospira* during rotational filtration. Individual harvesting events removed approximately 36-76% of the suspended biomass and were followed by re-fertilization of the returned medium (Section 2.1; Supplementary Figure S2). The resulting selective biomass removal and nutrient pulse provide a plausible mechanism for the transient compositional shifts associated with harvesting events; however, because no harvesting-free control was included, this interpretation remains a hypothesis rather than a demonstrated causal effect. The difference between suspension and harvested fractions therefore primarily reflects size-selective biomass separation rather than biomass originating from a distinct biofilm compartment. Because the harvested fraction was strongly dominated by *Cyanobacteriota*, nitrogen exported with harvested biomass was predominantly associated with the *Arthrospira*-rich fraction and was accounted for accordingly in the estimation of biomass-associated nitrogen (Section 3.6).

In addition to prokaryotic taxa, microbial eukaryotes contributed to the temporal community patterns observed in the metagenomic rRNA data. *Alveolata* increased during the initial sampling window and declined again in the later sampling window (Figure 4A), whereas *Opisthokonta* were detected only at early time points. Alveolates include ciliates with established roles in nutrient cycling in microbial ecosystems.^38^ However, their functional contribution was not assessed here, and the ecological drivers underlying these temporal patterns remain unresolved.

Together, these results demonstrate that while the switch to poultry-house exhaust introduced new environmental pressures, the photobioreactor maintained a stable enrichment of the target phototrophic organism, with transient dynamics in associated microbial populations. This balance between stability and adaptability provides a foundation for sustained biomass production under fluctuating conditions. To further investigate the functional implications of these community dynamics, we next analyzed metagenome-assembled genomes (MAGs) and the metabolic potential of the system.

### 3.4. Reconstruction of Metagenome-Assembled Genomes and Functional Annotation

Metagenome-assembled genomes (MAGs) were reconstructed and filtered to retain high-quality genomes (>50% completeness, <5% contamination) classified to at least the genus level (Supplementary Table S2). In total, 38 stage-specific MAGs were identified across the three aeration stages, comprising 19, 8 and 11 MAGs uniquely associated with the before,initial and later stages, respectively (Supplementary Figure S5). In addition, 33 MAGs were detected across multiple stages, indicating partial persistence of certain populations alongside pronounced taxonomic turnover over time.

Despite these shifts in community composition, analysis of metabolic pathway completeness across MAGs revealed largely conserved functional potential (Supplementary Figure S6). Although some variation was observed, amino-acid biosynthesis and other core metabolic pathways were broadly represented across the recovered MAGs, without a consistent sampling-window-specific pattern. We therefore interpret these results as broadly conserved biosynthetic potential rather than evidence of specific metabolic interactions such as cross-feeding. The representation of similar core metabolic functions across taxonomically distinct MAGs is consistent with functional redundancy at the level of encoded metabolic potential.^39, 40^

To describe the functional gene potential of the community beyond genome-level pathway completeness, gene-centric analyses were performed across the metagenomes. Genes involved in carbon fixation and nitrogen metabolism (Supplementary Table S4) were quantified based on normalized coverage and aggregated at the pathway level (Figure 5).

**Figure 5:**
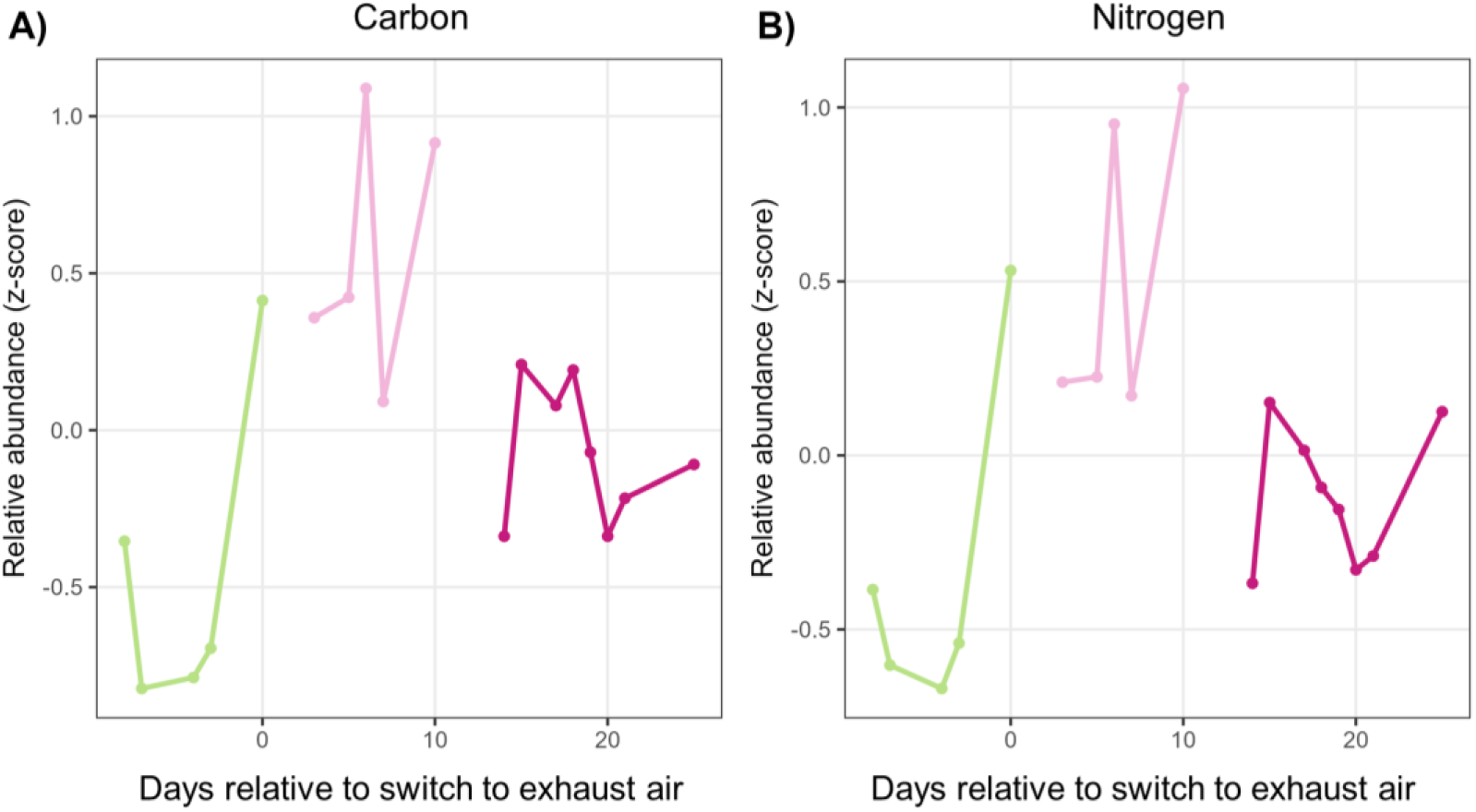
Temporal dynamics of carbon- and nitrogen-related functional potential across sampling windows. Mean normalized gene abundances (z-scores) are shown for genes associated with (A) carbon and (B) nitrogen metabolism. Each point represents the mean z-score of genes assigned to the respective functional category at a given sampling time. Gene abundances were standardized across samples for each gene, such that positive and negative z-scores indicate relative enrichment or depletion, respectively, compared with that gene’s mean abundance across the dataset. The profiles therefore describe temporal variation in the community-level representation of metabolic genetic potential, complementing the taxonomic analyses, and should not be interpreted as direct measures of transcriptional activity, metabolic rates, or process fluxes. Gene-specific patterns are provided in Supporting Information (Supplementary Figure S7).

This analysis revealed temporal shifts in the relative abundance of genes associated with carbon and nitrogen metabolism. During the initial phase, genes involved in central carbon metabolism (e.g., *pgk*, *fumC*) and nitrogen transformation pathways, including denitrification-associated genes (e.g., *narG*, *nosZ*), showed increased relative representation in the assembled metagenomes (Figure 5; Supplementary Figure S7). In the later phase, their relative abundances declined from the initial peak and stabilized closer to the overall mean. Genes associated with autotrophic carbon fixation, including components of the Calvin-Benson-Bassham cycle (e.g., *rbcL*, *prk*), remained consistently represented across sampling windows (Supplementary Figure S7).

Together, these results describe a temporal shift in the functional gene potential of the community. Because metagenomic gene abundance reflects genetic potential rather than expression or flux, these patterns are interpreted only at the level of functional gene representation and not as evidence of corresponding metabolic rates.

### 3.5. Impact of cultivation matrices on the microbiome

Building on the functional transitions observed in suspension cultures, we investigated whether defined cultivation matrices could provide additional insight into microbiome organization and ammonia-related processes in spatially structured niches.^11, 12, 41^ For this purpose, two carrier materials were introduced into the photobioreactor: an in-house developed macroporous elastomeric silicone foam (MESIF), previously shown to support the sampling of complex environmental microbiomes,^19^ and the commercially available MUTAG™ biocarrier. The matrices were introduced at low loading as standardized colonization substrates rather than as a process-scale biofilm module. They provided localized attached-growth niches for characterizing microorganisms that preferentially established on structured surfaces relative to the circulating suspension.

Microbial community profiling across defined time points before (−7 to 0) and after aeration (3 to 25 days; Supplementary Table S5) showed that matrix-associated communities differed from the suspension and exhibited greater structural stability across aeration stages, as reflected in diversity patterns (Supplementary Figure S8). In particular, suspension communities showed stronger aeration-associated shifts, whereas MESIF- and MUTAG™-associated communities clustered primarily by aeration stage rather than by carrier type, indicating that matrices captured related but more buffered surface-associated microbiomes.

Taxonomic profiling based on 16S rRNA gene amplicons revealed pronounced enrichment of members of the candidate phyla *Eremiobacteriota* and *Hydrogenedentota*, particularly after prolonged aeration (Supplementary Figure S9). Differential abundance analysis further identified clear taxonomic differences between matrix-associated and planktonic communities (Supplementary Figure S10A). Notably, genera involved in nitrogen transformation, including *Nitrosomonas*, were enriched in both MESIF and MUTAG™ relative to the suspension, consistent with preferential establishment of nitrifying microorganisms in surface-associated niches.^13–15, 17^ In addition, several enriched genera with known surface-associated lifestyles, including *Hyphomonas*^42^ and *Pirellula*,^43^ supported selective enrichment of biofilm-associated communities.

Comparison between carrier types showed that MESIF enriched a broader range of taxa than MUTAG™ (Supplementary Figure S10B-C), including *Methylobacterium*, associated with metabolically versatile lifestyles and diverse nitrogen utilization strategies,^44^ and *Halomonas*, frequently linked to nitrogen transformation processes under variable environmental conditions.^45, 46^ These patterns suggest that MESIF provides more diverse microenvironments that support specialized microbial populations.

To investigate whether these enrichment patterns were also reflected at the metagenomic level, selected matrix-associated samples were sequenced and compared with suspension communities (Figure 6). Both matrices showed enrichment of eukaryotic members, particularly *Opisthokonta* and *Alveolata*, at later time points, whereas suspension communities became more strongly dominated by prokaryotic taxa. In addition, enrichment of *Eremiobacteriota* and *Hydrogenedentota* in matrix-associated samples confirmed the trends observed in the amplicon data. In addition, matrix-associated communities showed a relative enrichment of *Pseudomonadota* and a concomitant decrease in *Cyanobacteriota* compared to suspension samples, indicating a clear separation between surface-associated heterotrophic populations and planktonic phototrophic biomass (*Arthrospira*).

**Figure 6:**
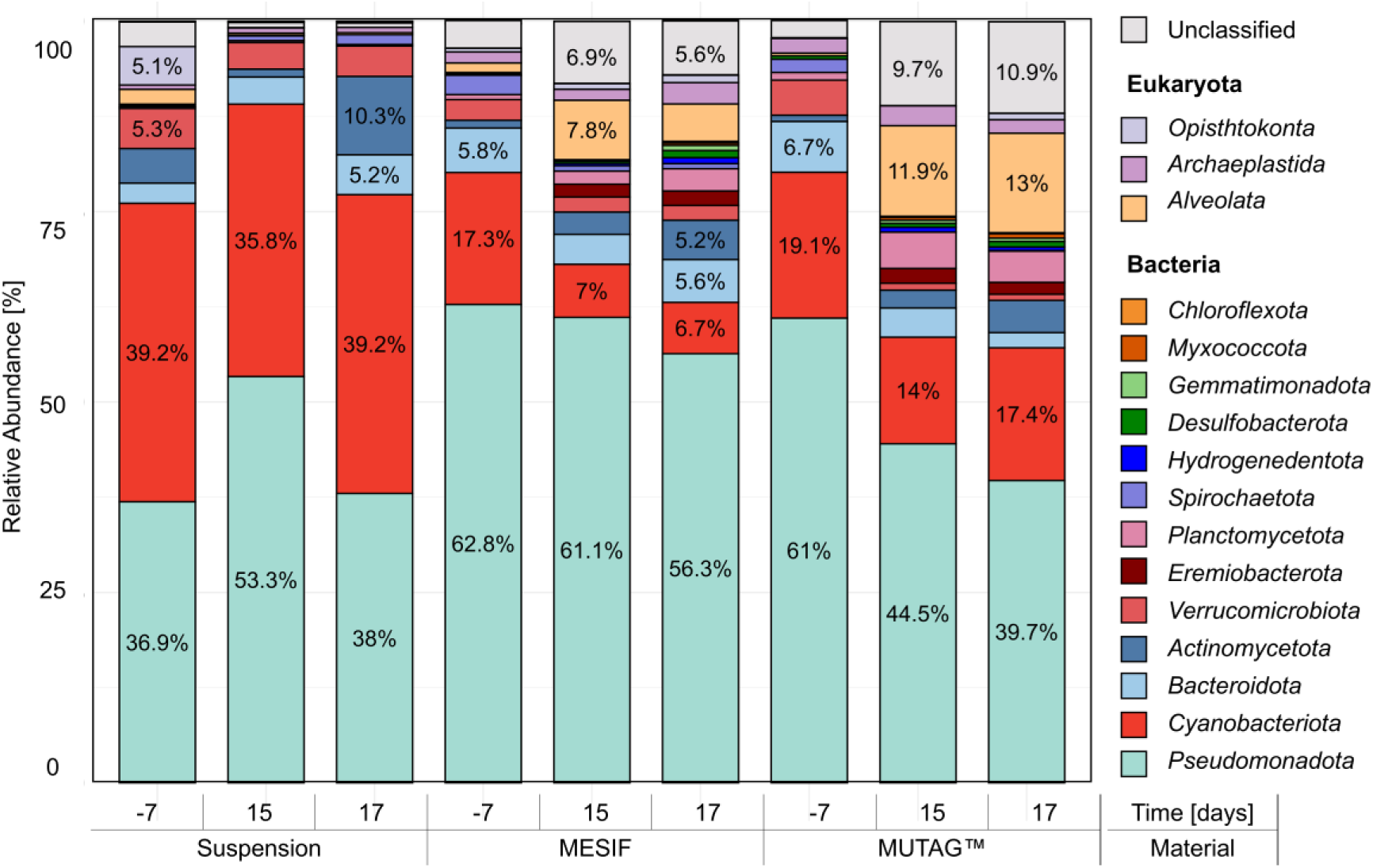
Phylum-level taxonomic composition of microbial communities in suspension, MESIF, and MUTAG™ samples across three time points. Relative abundances are shown for phyla representing more than 5% of the community.

To further assess whether these taxonomic enrichments were accompanied by distinct genome-resolved populations, MAGs were reconstructed from matrix-associated samples. After quality filtering and removal of well-characterized genomes, 50 unique MAGs were obtained exclusively from matrix-associated communities, including 10 unique to MESIF, 7 unique to MUTAG™, and 33 shared between both matrices (Supplementary Tables S6-S7). Matrix-associated MAGs included representatives of *Eremiobacteriota* and *Hydrogenedentota*, providing genome-level support for the presence of these underexplored lineages.

While predicted metabolic profiles were broadly similar across MAGs (Supplementary Figure S11), their relative abundances differed markedly when mapped across all metagenomic assemblies (Figure 7). Most MAGs were strongly enriched within the matrices and increased in abundance over time, whereas only a subset was detected in suspension samples.

**Figure 7.**
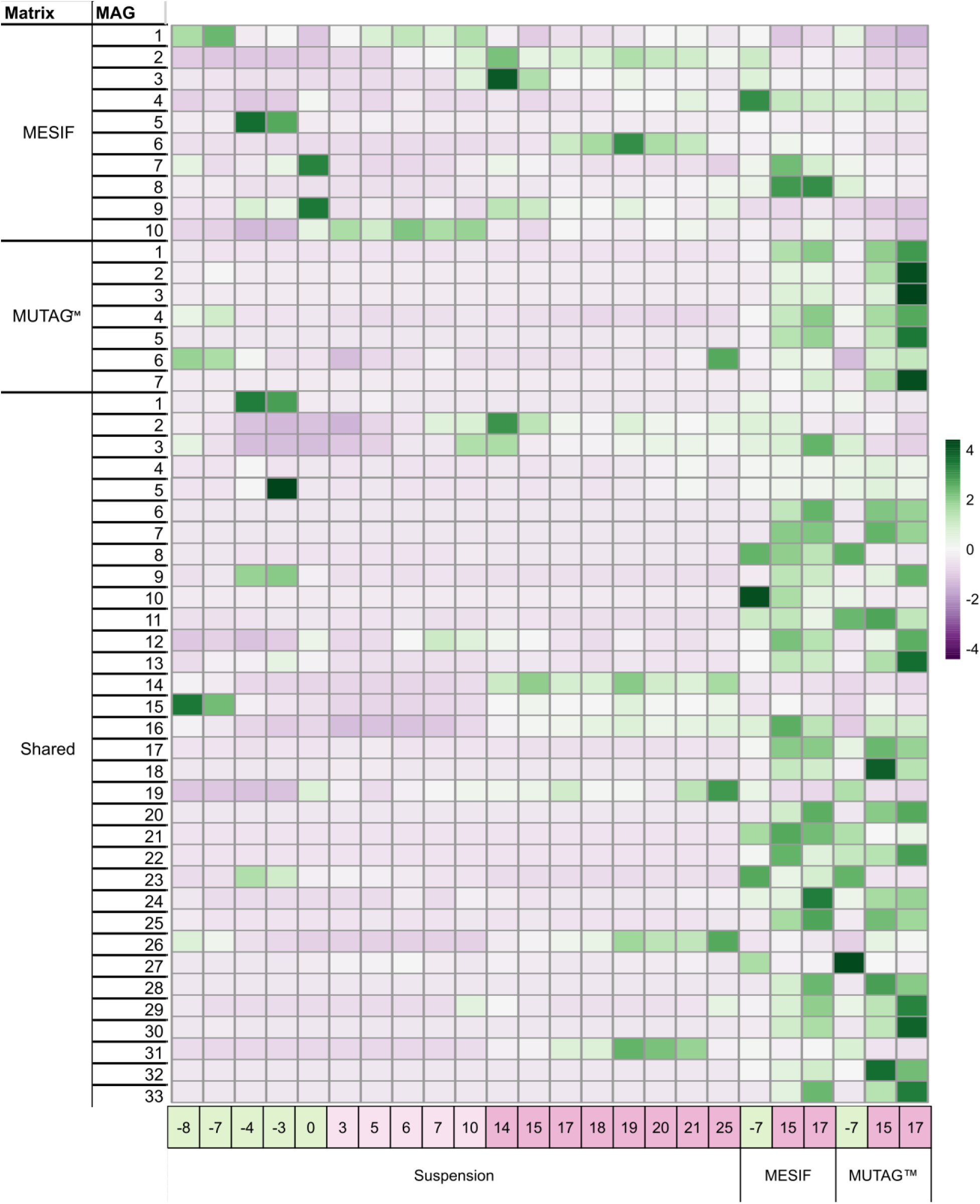
Normalized relative abundance of matrix-derived metagenome-assembled genomes (MAGs) across all metagenomic assemblies. MAGs reconstructed exclusively from MESIF, exclusively from MUTAG™, or shared between both matrices are shown across suspension and matrix-associated samples. Color intensity indicates relative abundance (green: higher; purple: lower). Note that MAGs are predominantly detected in matrix-associated samples and show increasing abundance over time, whereas only a subset is present in suspension samples, indicating matrix-specific enrichment patterns.

Together, these results indicate that the cultivation matrices act as ecological filters by providing local habitat conditions, including surface attachment and reduced light exposure, that selectively favor the establishment or persistence of particular taxa relative to the bulk suspension. This promotes spatially structured microbial communities in which taxa underrepresented in the suspension can become enriched. While biofilm formation was not directly quantified (e.g., via extracellular polymeric substance (EPS) measurements), the consistent enrichment of surface-associated taxa and clear compositional separation from suspension communities support the presence of biofilm-like communities on the carrier materials.

Despite their different material properties and geometries, MESIF and MUTAG™ supported largely similar microbial communities, as also apparent from their taxonomic profiles (Supplementary Figure S9). Samples clustered primarily by sampling window rather than by carrier material (Supplementary Figure S8), 33 of the 50 carrier-derived MAGs were shared between MESIF and MUTAG™ (Supplementary Tables S5-S6), and only 12 genera differed significantly between the two carrier types (Supplementary Figure S10B-C). Given that both carrier types were exposed within the same reservoir compartment without direct artificial illumination, these findings indicate that carrier identity had a comparatively limited influence on overall attached-community composition under the conditions investigated here.

### 3.6. Nitrogen dynamics and carrier-associated ammonia-oxidizing potential

To complement the direct exhaust-aeration campaign, a separate experiment was performed using air-scrubber process water as a defined liquid nitrogen input. The process water had already undergone processing within the air-scrubbing system before addition to the photobioreactor. Biological livestock air scrubbers can both capture gaseous NH_3_ into the aqueous phase and biologically convert part of the ammoniacal nitrogen to nitrate.^18^ Accordingly, nitrate was already present in the supplied process water and was reintroduced during successive replenishment cycles. The experiment was therefore used to evaluate changes in measured dissolved inorganic nitrogen (DIN) and its recovery in biomass rather than to infer nitrogen transfer directly from the gas phase or in-reactor nitrification.

Across all cycles, DIN decreased consistently (Figure 8B), corresponding to removal efficiencies between approximately 56% and 80% (mean ∼68%, Figure 8B). Only ∼24-58% of this decrease was recovered as biomass-associated nitrogen, indicating that the decrease in measured DIN was only partially recovered in harvested biomass. The fate of the remaining nitrogen cannot be resolved from the present measurements and may include unmeasured nitrogen pools and gaseous losses.

**Figure 8:**
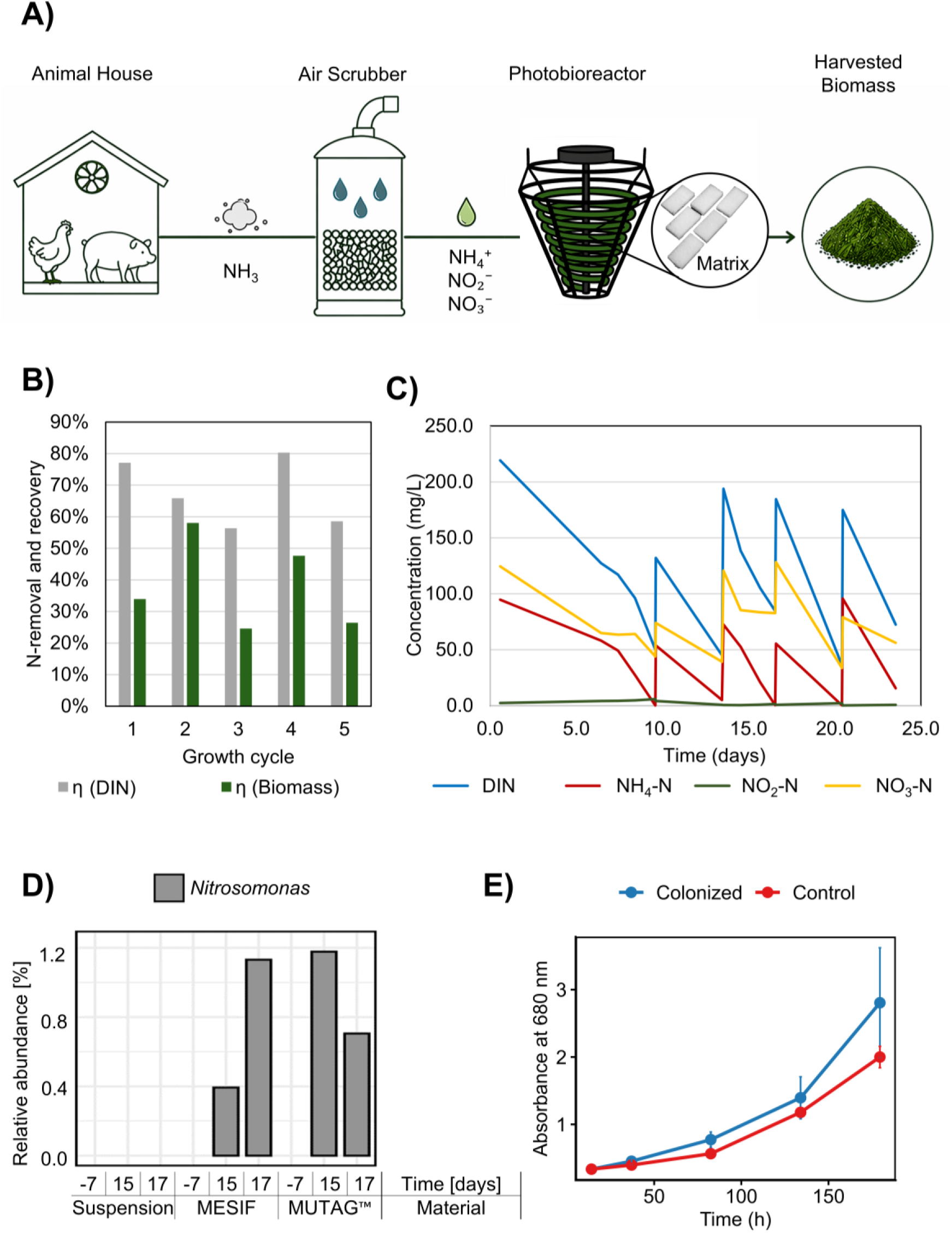
Nitrogen partitioning and ammonia-oxidizing microorganisms in spatially structured niches within the photobioreactor system. (A) Schematic representation of nitrogen transfer from livestock exhaust via an air scrubber into the liquid phase and subsequent input into the photobioreactor. (B) Cycle-based comparison of the decrease in dissolved inorganic nitrogen (DIN; NH_4_^+^ + NO_2_^-^ + NO_3_^-^) (grey) and nitrogen recovered in harvested biomass (green), showing that only part of the decrease in measured DIN was recovered in biomass. (C) Temporal profiles of dissolved inorganic nitrogen species showing rapid depletion of NH₄⁺ (red), transient accumulation of NO₂⁻ (green), and persistence of NO₃⁻ (yellow). (D) Relative abundance of *Nitrosomonas* in matrix-associated and suspension communities. (E) Re-inoculation experiment showing enhanced early biomass development and a shorter lag phase in initially axenic *A. maxima* cultures supplemented with pre-colonized MESIF compared with non-colonized controls.

Analysis of individual nitrogen species revealed rapid depletion of NH₄⁺ to near-zero concentrations and minimal, transient NO_2_^-^ accumulation, while NO_3_^-^ remained present throughout the cultivation cycles (Figure 8C). Because nitrate was already present in the scrubber-derived input and replenished during successive cycles, these concentration profiles cannot be interpreted as evidence for in-reactor nitrification. Moreover, the decrease in DIN cannot be attributed to nitrification, which redistributes NH_4_^+^ among dissolved inorganic nitrogen species rather than removing nitrogen from the DIN pool.

The ammonia-oxidizing genus *Nitrosomonas* was preferentially associated with both carrier materials and absent from suspension samples (Figure 8D; Figure S10A). A *Nitrosomonas*-affiliated MAG (Shared MAG17) was likewise recovered exclusively from matrix samples (Figure 7). This enrichment within the low-light attached niche is consistent with previous observations that nitrifying microorganisms can be inhibited by high irradiance and may benefit from protected attached-growth habitats.^13, 16^

Carrier-derived MAGs also revealed genetic potential for nitrate reduction: 19 of 47 MAGs (40.4%) encoded at least one nitrate reductase gene (*narG*, *narZ*, or *napA*) (Supplementary Figure S12; Table S8). MAGs encoding these genes were predominantly shared between MESIF and MUTAG™, and no individual MAG contained the complete set of genes examined for canonical denitrification. Thus, the carrier-associated communities contained genetic potential for both ammonia oxidation and nitrate reduction, but these genomic observations indicate functional potential rather than pathway activity or quantitative nitrogen flux.

Dissolved oxygen was not monitored directly. Although the continuously aerated airlift configuration and additional aeration of the reservoir make persistent anaerobic conditions in the bulk liquid unlikely, local oxygen gradients within attached communities cannot be excluded. Consequently, the occurrence and quantitative contribution of nitrification, nitrate reduction, or coupled nitrification-denitrification cannot be established from the present data.

To assess the transferability of matrix-associated communities, re-inoculation experiments were performed using pre-colonized MESIF carriers and an initially axenic *Arthrospira maxima* culture. Cultures supplemented with pre-colonized MESIF exhibited faster early biomass development and a shorter lag phase than controls containing non-colonized matrices (Figure 8E; Supplementary Figure S13). The effect was most pronounced during the first ∼200 h, indicating that transfer of pre-established matrix-associated communities can facilitate early culture establishment.

## 4. Implications

This study links microbial-community dynamics and spatial niche differentiation in an operational photobioreactor to complementary measurements of nitrogen partitioning under controlled nitrogen supply. During cultivation with air-scrubber process water, the decrease in measured dissolved inorganic nitrogen (DIN) exceeded the amount of nitrogen recovered in harvested biomass, demonstrating that only part of the decrease in DIN was accounted for by biomass production. The fate of the remaining nitrogen cannot be resolved from the present measurements and, importantly, cannot be attributed to nitrification, which redistributes nitrogen among dissolved inorganic species rather than removing it from the DIN pool.

Under the alkaline and continuously aerated cultivation conditions, NH_3_ volatilization represents a plausible pathway contributing to the unresolved nitrogen fraction. At 25 °C, the pK`ₐ` of the NH_4_^+^/NH_3_ equilibrium is approximately 9.25, corresponding theoretically to approximately 64% of total ammoniacal nitrogen being present as un-ionized NH_3_ at pH 9.5 and approximately 97% at pH 10.7.^47^ Continuous aeration may therefore promote NH_3_ stripping. However, these equilibrium fractions do not quantify the actual volatilized nitrogen flux, which was not measured in the scrubber-water experiment. Ammonia volatilization has been demonstrated experimentally under alkaline wastewater-treatment conditions,^48^ supporting its consideration as a plausible contributor to the unresolved nitrogen fraction rather than a quantitatively established loss pathway.

From a process-integration perspective, the large mismatch between livestock ventilation rates and the gas throughput of the photobioreactor highlights a fundamental scaling challenge for direct exhaust-air coupling. In the present setup, approximately 1 m^3^ h^-1^ of a total poultry-house ventilation flow of approximately 175 m^3^ h^-1^ was directed through the 150 L reactor, corresponding to only ∼0.6% of the total exhaust stream. At practically relevant scales, an intermediate concentration step such as air scrubbing may therefore be required to decouple treatment of large volumes of dilute agricultural exhaust from nitrogen supply to the photobioreactor. By concentrating emission-derived nitrogen into a substantially smaller aqueous stream, air scrubbing could enable controlled nitrogen dosing without requiring the entire ventilation flow to pass through the cultivation system

At the microbial-community level, *Nitrosomonas* was preferentially associated with the carrier materials, indicating that ammonia-oxidizing microorganisms established within the attached, low-light niche. This spatial association is consistent with the known sensitivity of nitrifiers to high irradiance and their preferential establishment in protected attached-growth environments.^13, 16^ However, the quantitative contribution of these microorganisms to reactor-scale nitrogen transformation was not determined.

Similar changes in dissolved nitrogen species concentrations were also observed during reactor operation in the absence of added carrier materials, indicating that these patterns were not dependent on the presence of the carriers. During prolonged reactor operation, thin, visibly adherent deposits with a biofilm-like appearance were observed on reactor surfaces, particularly in the reservoir compartment without direct artificial illumination, suggesting the presence of additional surface-associated microbial niches. However, these deposits were not systematically sampled, quantified, or microbiologically characterized in the present study, and their contribution to the observed nitrogen concentration dynamics therefore remains unresolved.

The preferential association of ammonia-oxidizing microorganisms with the carrier communities is consistent with previous reports that nitrifiers can be inhibited by high irradiance and benefit from protected attached-growth environments in illuminated algal-bacterial systems.^13–16^ In the present system, the carriers were confined to the reservoir compartment without direct artificial illumination; surface attachment and reduced light availability therefore co-varied. The enrichment of *Nitrosomonas* should consequently be interpreted in the context of this combined attached, low-light niche rather than as an effect of carrier attachment alone. Importantly, taxonomic enrichment indicates the presence of ammonia-oxidizing microorganisms but does not establish their quantitative contribution to reactor-scale nitrogen conversion.

These findings highlight an important limitation of suspension-based analyses in photobioreactor systems, as key functional processes may remain undetected without resolving spatial heterogeneity. The use of structured carrier materials such as MESIF and MUTAG™ therefore provides a means to recover and characterize microbial populations associated with localized attached-growth niches that may remain underrepresented in suspension-based analyses. Beyond enabling taxonomic discovery, including enrichment of underexplored lineages such as *Eremiobacteriota* and *Hydrogenedentota*, these matrices provide ecological niches that support microorganisms sensitive to planktonic and illuminated conditions.^16^

Importantly, the functional relevance of these matrix-associated communities was further supported by re-inoculation experiments, which demonstrated that pre-colonized carrier materials can enhance culture establishment and biomass development in independent systems using a different *Arthrospira* strain. This suggests that structured matrices can serve as reservoirs of pre-adapted microbial consortia and enable transfer of functional potential across cultivation setups.

Taken together, our results highlight how spatial organization and microbial niche differentiation contribute to the ecological structure of photobioreactor communities. This has implications for the design of bio-based production systems, as the targeted integration of structured carrier materials may provide niches for specific microbial guilds, although their contribution to reactor-scale processes remains to be demonstrated. From an application perspective, valorization of the harvested *Arthrospira*-rich biomass as a protein-rich feed ingredient or supplement represents a potential route for local nutrient recovery and recirculation within agricultural systems. Previous studies have evaluated the amino-acid composition, mineral and heavy-metal content, and microbiological quality of microalgal biomass cultivated using poultry-house exhaust air, providing an initial basis for considering this valorization route.^7, 9, 10^ However, the present study did not assess feed or food safety, and any such application would require appropriate compositional, toxicological, microbiological, and regulatory evaluation.

More broadly, this work demonstrates how combining systems-level measurements with spatially resolved microbial ecology can bridge the gap between descriptive community analysis and mechanistic understanding. Such approaches are essential for advancing the rational design of microbial consortia for sustainable biotechnology applications, particularly in systems that integrate waste valorization with biomass production.

Several limitations should be noted. The study was conducted using a single 150 L reactor operated once; the reported samples therefore represent repeated measurements from the same system rather than independent biological units, and differences between sampling windows are described rather than tested as reactor-level effects. Moreover, because exhaust-air aeration was applied continuously from day 0, temporal changes during exposure are inherently confounded with elapsed time and cumulative reactor operation, including periodic harvesting and re-fertilization. Indeed, harvesting events coincided with larger short-term shifts in community composition than the onset of aeration itself (Figure 4A, “H”). Consequently, differences between sampling windows cannot be attributed specifically to the duration of exhaust-air exposure.

In addition, the carriers were confined to the reservoir compartment without direct artificial illumination, meaning that surface attachment and reduced light availability co-varied in this design. Given the known light sensitivity of nitrifying microorganisms,^13, 16^ the enrichment of *Nitrosomonas* and other carrier-associated taxa should therefore be interpreted as a response to this combined attached, low-light niche rather than to surface attachment alone.

During the poultry-exhaust-air experiment, dissolved nitrogen species were not systematically quantified and a complete nitrogen balance was therefore not determined. Gas-phase NH_3_ concentrations were assessed manually at both the reactor inlet and outlet using colorimetric test strips. Although these measurements indicated lower NH_3_ concentrations at the outlet than at the inlet, the limited quantitative resolution of the method, together with substantial temporal variability in exhaust-air NH_3_ concentrations, precluded reliable determination of NH_3_ removal efficiency or a quantitative gas-phase nitrogen balance.

Dissolved oxygen was not monitored directly. Although continuous aeration makes persistent anaerobic conditions in the bulk liquid unlikely, local oxygen gradients within attached communities cannot be excluded. Moreover, metagenomic detection of ammonia-oxidizing taxa and nitrogen-cycle genes indicates functional potential rather than activity or quantitative flux. Consequently, the occurrence and reactor-scale contribution of nitrification, nitrate reduction, or coupled nitrification-denitrification cannot be determined from the present data.

## Acknowledgements

This work was financially supported through the Helmholtz program “Materials Systems Engineering” under the topic “Adaptive and Bioinstructive Materials Systems” (43.33.11), KIT EXU project DigiteLiSE, BMBF project 161L0284A MicroMatrix, and BMBF project 031B0915U1 UrbanAqua under the umbrella “Innovationsraum Bioökonomie auf Marinen Standorten”. The authors thank Anna Edinger, David Thiele and Simone Weigel for experimental help.

## Supporting Information

Supporting Information (SI) includes Extended Methods (Sections S1-S4), Figures S1-S13, Tables S1-S8, and additional references (49-82). Section S1 provides detailed descriptions of microalgae cultivation, reactor operation, sampling procedures, and exhaust gas characterization. Section S2 describes nitrogen mass balance calculations and analytical methods. Section S3 includes detailed protocols for carrier preparation, matrix incubation, and re-inoculation experiments. Section S4 outlines DNA extraction, sequencing, and bioinformatic analyses.

Figure S1 shows the correlation between optical density (OD₅₆₅) and dry biomass of *A. platensis*. Figure S2 presents temporal CO_2_ profiles of the supplied exhaust air together with biomass dynamics. Figure S3 shows alpha and beta diversity analyses of suspension communities. Figure S4 displays metagenomic taxonomic composition across aeration stages. Figure S5 summarizes metagenome-assembled genomes (MAGs) across conditions. Figure S6 presents metabolic pathway completeness of MAGs. Figure S7 shows functional gene dynamics related to carbon and nitrogen metabolism. Figure S8 presents alpha and beta diversity analyses of matrix-associated communities. Figure S9 shows taxonomic composition of matrix-associated microbiomes. Figure S10 presents differential abundance analysis of matrix-associated communities. Figure S11 summarizes metabolic profiles of matrix-derived MAGs. Figure S12 presents the distribution of denitrification-associated genes. Figure S13 shows results of re-inoculation experiments.

Table S1 provides a sampling and sequencing overview of the experiment. Table S2 lists differentially abundant taxa identified by ANCOM-BC. Table S3 summarizes unique MAGs detected in suspension samples. Table S4 provides the list of genes associated with carbon fixation and nitrogen metabolism. Table S5 provides an overview of sampling time points for photobioreactors with cultivation matrices. Table S6 contains unique MAGs recovered from MESIF and MUTAG matrices. Table S7 lists shared MAGs recovered from both cultivation matrices. Table S8 shows the detailed annotation of denitrification-derived hits within the carrier-associated MAGs.

