## Supplementary Information for "Integrating Microbial Ecology and Sustainability: Linking Agricultural Waste Gas Streams to Microbial Community Shifts in a Photobioreactor"

### Supporting Information

Marta Velaz Martín[1], Till Glockow[2], Laura Meisch[1], John Vollmers[3], Anne-Kristin Kaster[3], Kersten S. Rabe[1], Christof M. Niemeyer[1]

[1] Karlsruhe Institute of Technology (KIT), Institute for Biological Interfaces 1 (IBG-1), Biomolecular Micro- and Nanostructures, Hermann-von-Helmholtz-Platz 1, D-76344 Eggenstein-Leopoldshafen, Germany.

[2] Acheron GmbH, Auf der Muggenburg 30, 28217, Bremen, Germany

[3] Karlsruhe Institute of Technology (KIT), Institute for Biological Interfaces 5 (IBG-5), Biotechnology and Microbial Genetics, Hermann-von-Helmholtz-Platz 1, D-76344 Eggenstein-Leopoldshafen, Germany.

### 1. Extended Methods

#### SI Section S1 - 1.1. Microalgae cultivation; animal housing setup; sample preparation

A mixed culture of *Arthrospira platensis* (*A. platensis*) originating from an open pond (ROVAL GmbH, Germany) was first propagated in 10-L algal bags and subsequently used to inoculate a closed photobioreactor (~150 L), as reported previously.[7] In short, the saline mineral medium was prepared with tap water supplemented with 1.45 g/L Agrolution 324 fertilizer, 0.19 g/L KALISOP, 0.033 g/L Micromax WS Fe Chelate, 1 g/L Canna Trace Mix Mono, 5.5 g/L sodium chloride, and 9 g/L sodium bicarbonate, achieving final concentrations of 203 mg/L nitrogen, 51 mg/L phosphorus, 344 mg/L potassium, 35 mg/L sulfur, 4.3 mg/L iron, and 17 mg/L magnesium. The photobioreactor was operated with continuous aeration (15 L $min^{-1}$) and LED lighting (Figure 1A). Following transfer from the algal bags, the photobioreactor was operated for 7 days with continuous ambient-air aeration (15 L $min^{-1}$; ~400 ppm $CO_2$; $NH_3$ not detectable) to allow acclimation of the culture to the airlift photobioreactor. On day 0, the gas source was switched from ambient air to poultry-house exhaust air, which was then supplied continuously until the end of the experiment.

$CO_2$ concentrations in the supplied poultry-house exhaust (reactor inlet) and after passage through the photobioreactor (reactor outlet) were measured using T6545 meters (COMET SYSTEM, Bezrucova, Czech Republic). $CO_2$ inlet concentrations varied between day and night due to chickens being outdoors during the day and indoors at night, with average concentrations of 1306 ppm during the day, 2119 ppm at night, and a combined average of 1716 ppm.

The photobioreactor system included a connected dark reservoir compartment (approx. 30-40 L), which was not directly exposed to light and provided a low-light environment distinct from the illuminated tubular reactor.

Biomass was harvested from the suspended culture via rotational filtration; carrier-associated biomass was not harvested. Biomass concentration was monitored by optical density at 565 nm ($OD_{565}$) using a previously established correlation derived from the same reactor system,[7] which was validated by gravimetric dry weight measurements under the conditions of this study ($R^2$ = 0.9973; Figure S1). Based on biomass concentrations immediately before and after harvesting, individual harvesting events reduced the suspended biomass concentration by approximately 36–76% (Figure S2). The clarified medium obtained after rotational filtration was post-fertilized with 720 g/kg harvested dry weight Agrolution 324, 65 g/kg KALISOP, 10 g/kg Micromax WS Fe chelate, 100 g/kg Canna Trace Mix Mono, and 2000 g/kg sodium bicarbonate before being returned to the reactor.

The chicken coop (16 $m^2$, 54 $m^3$) housed 30 laying hens and one rooster, with daytime access to an outdoor area via an automatic flap. A ventilation system extracted 175 $m^3$/h of exhaust air, with 1 $m^3$/h redirected into the helical tube photobioreactor through a pump. Thus, approximately 0.6% of the total poultry-house ventilation stream was directed through the photobioreactor.

Ammonia concentrations in the reactor inlet and outlet gas were assessed manually using colorimetric test strips. Measurements indicated $NH_3$ concentrations of approximately 35–40 ppm in the supplied exhaust air and approximately 0–5 ppm in the reactor outlet. Because of the semi-quantitative nature of the test strips and the pronounced temporal variability of the exhaust-air composition, these measurements were used for indicative monitoring rather than for calculation of gas-phase removal efficiencies.

Samples of the algal suspension were collected over 33 days and assigned to three temporal sampling windows relative to the switch from ambient air to poultry-house exhaust air: before exhaust-air exposure (B; during the 7-day ambient-air acclimation period), initial exhaust-air exposure (I; days 1–10), and later exhaust-air exposure (L; days 14–25) (Figure 1B, main manuscript). These categories represent temporal sampling windows rather than distinct aeration regimes; the reactor remained continuously aerated throughout the experiment.

Suspension samples (aliquots obtained directly from the circulating culture) were withdrawn from the circulating culture via a lateral sampling port of the helical tube reactor. To minimize potential sedimentation effects, an initial volume (~100 mL) was discarded prior to sampling. Subsequently, ~100 mL of culture were collected and divided into two aliquots. One aliquot (50 mL) was used immediately for optical density measurements at 565 nm ($OD_{565}$), performed in triplicate using an IRIS photometer (Hanna Instruments, Germany). The second aliquot (50 mL) was stored at −28 °C for downstream analyses. Moreover, harvested biomass samples were withdrawn from the concentrated biomass fraction obtained during routine harvesting events, in which a defined portion of the culture was removed and subjected to biomass separation.

Reactor performance and biomass development were monitored over time using optical density measurements ($OD_{565}$). The culture exhibited stable growth under semi-continuous operation with periodic harvesting and medium replenishment. Detailed temporal profiles of biomass development, including growth rates and harvesting cycles, are provided in Figure S2.

### SI Section S2 - Nitrogen Mass Balance Estimations

Air-scrubber process water (ASW) was obtained from a biological air scrubber connected to a pig housing facility in Lower Saxony, Germany. In this system, exhaust air is passed through a packed bed and continuously irrigated with recirculating liquid, enabling the absorption of gaseous ammonia ($NH_3$) into the aqueous phase. Microbial activity within the scrubber leads to the accumulation of ammonium ($NH_4^+$) and partial oxidation to nitrite ($NO_2^-$) and nitrate ($NO_3^-$), resulting in a nitrogen-rich process water stream. Prior to use, the ASW was gravity-filtered using paper filter material to remove coarse suspended solids but was not sterilized, thereby preserving its native microbial and chemical characteristics. The ASW was used exclusively in nitrogen dynamics experiments and was not applied during the cultivation experiments described in Sections 2.1 and 3.1-3.5.

For nitrogen dynamics experiments, ASW was periodically added to the photobioreactor during semi-continuous operation as an external nitrogen source. The volume of ASW added was adjusted to achieve dissolved inorganic nitrogen concentrations (DIN) in the range of approximately 200 $mg \cdot L^{-1}$ within the reactor, enabling controlled assessment of nitrogen transformation processes.

Nitrogen removal and transformation were assessed based on DIN and individual inorganic nitrogen species ($NH_4^+$, $NO_2^-$, $NO_3^-$), quantified photometrically using an HI801 Iris Photometer (Hanna Instruments, Germany). Measurements were performed in technical triplicates, and mean values with standard deviations were calculated.

Nitrogen dynamics were evaluated both as time-resolved concentration profiles and through cycle-based mass balances. Each cycle k was defined as the period between two harvesting events.

For each cycle, the change in DIN (ΔDIN,k) was calculated as the difference between initial and final concentrations and converted to mass by multiplication with the reactor volume (145 L):

$$\Delta DIN,k = (C\_start - C\_end) \times V$$

Nitrogen incorporation into biomass (N_biomass) was estimated based on biomass production and an assumed nitrogen content of Arthrospira. Biomass concentrations were approximated using $OD_{565}$ measurements, as described in Section S1. Protein content was quantified by an external laboratory, and nitrogen content was subsequently calculated using the Jones factor (6.25 g protein per g nitrogen), a standard conversion to estimate nitrogen from total protein.

Nitrogen closure for each cycle was calculated as

$$\eta\_biomass = N\_biomass / \Delta DIN$$

The remaining fraction (1 – η_biomass) represents the decrease in measured DIN not recovered in harvested biomass and may include unmeasured nitrogen pools and gaseous nitrogen losses.

### SI Section S3 - Cultivation in matrix materials

#### S3.1 Matrix materials and preparation

To investigate the spatial organization of microbial communities into planktonic and surface-associated niches, inert carrier matrices were introduced into the photobioreactor. Macroporous elastomeric silicone foams (MESIF) and commercial biofilm carriers (MUTAG™ BioChip) were used as standardized substrates for biofilm formation. Macroporous elastomeric silicone foam (MESIF) pieces were produced produced as previously described.[19] In brief, a polydimethylsiloxane (PDMS) solution was prepared by mixing a prepolymer solution with a curing agent (Sylgard 184, Down Corning) in a 10:1 mass ratio. The mixture was degassed in a vacuum chamber for 45 min, then 0.9 mL of the mixture was poured into a polymethylmethacrylate (PMMA) mold. Sieved table salt grains (707-500 µm, ChanteSel iodized salt fine from Lidl, Germany) were added as porogens and homogeneously dispersed by centrifugation at 4600 rpm and 4 °C for 20 min (5804R, Eppendorf). The resulting mixture was cured in an oven (Heraeus, Thermo Scientific Inc.) at 70 °C for one hour. After demolding, the MESIF pieces were repeatedly pressed in a 50 °C water bath until the material reached a sponge-like structure. Finally, they were placed in a mechanical stirrer overnight to dissolve the remaining porogens and then cut into slices (10 x 10 x 3 mm). As an alternative carrier matrix, MUTAG Biochip™ (MUTAG™ BioChip, Multi Umwelttechnologie AG, Germany) was used. MUTAG™ Chips, which are made of polyethylene and have a diameter of 30 mm, were sterilized together with the MESIFs at 121 °C for 20 min (HP Medizintechnik GmbH, Germany).

#### S3.2 Deployment and sampling of matrices

Sterilized MESIF and MUTAG™ matrices were introduced into the photobioreactor under standard cultivation conditions. MESIF materials were manually compressed after insertion to ensure complete saturation of internal pore structures with culture medium. Matrices were incubated for defined time periods (Table 1) to allow microbial colonization and biofilm formation. At designated sampling time points, matrices were removed and processed for downstream analyses. Sampling durations and conditions are summarized in Table 1.

#### S3.3 Re-inoculation experiments

To assess whether matrix-associated microbial communities contribute to culture establishment and biomass development, re-inoculation experiments were performed using MESIF carrier matrices. Pre-colonized MESIF materials obtained from previous photobioreactor operation were used as inoculated carriers, while non-colonized MESIFs served as controls. Experiments were conducted using a different laboratory strain (*Arthrospira maxima*) to evaluate the transferability of matrix-associated effects across related cyanobacterial systems. Cultures were established in parallel under identical conditions in 1 L Schott flasks and aerated with exhaust air. Biomass development was monitored over time using absorbance measurements at multiple wavelengths associated with photosynthetic pigments (565 nm for phycocyanin and phycoerythrin, 680 nm for chlorophyll a, and 780 nm as a reference for biomass).

### SI Section S4 - Sequencing and Bioinformatics

#### S4.1 DNA extraction

Genomic DNA (gDNA) was extracted from 2 mL of algal suspension by centrifugation at 10,000 × g for 10 min, followed by resuspension in CD1 lysis buffer (DNeasy PowerSoil Pro Kit, QIAGEN, Germany). Extraction was performed according to the manufacturer's instructions. gDNA concentration was quantified using NanoDrop OneC (Thermo Scientific Inc.) and Qubit 3 fluorometric measurements (Thermo Scientific Inc.). The same extraction protocol was applied to matrix samples (MESIF and MUTAG™), where half of each carrier was processed under sterile conditions. To ensure pore saturation, matrices were compressed in lysis buffer prior to extraction.

**Table S1.** Sampling and sequencing overview of the photobioreactor experiment. Sampling days are given relative to the switch from ambient air to poultry-house exhaust air (day 0), and 'before', 'initial', and 'later' denote temporal sampling windows within the continuously operated reactor rather than distinct aeration regimes. Sample type abbreviations indicate suspension (SU), MESIF (ME), and MUTAG™ (MU). Suspension samples were analyzed by both 16S rRNA gene amplicon sequencing and shotgun metagenomics (n = 18). MESIF and MUTAG™ samples were analyzed primarily by 16S rRNA gene amplicon sequencing (n = 12 per carrier type), with shotgun metagenomic sequencing additionally performed for carrier samples collected at days −7, 21, and 25 (n = 3 per carrier type).

| Days relative to exhaust aeration | Sample type | Sampling window |
|---|---|---|
| -8 | SU | Before (B) |
| -7 | SU, ME, MU | |
| -4 | SU, ME, MU | |
| -3 | SU, ME, MU | |
| 0 | SU, ME, MU | |
| 3 | SU, ME, MU | Initial (I) |
| 5 | SU, ME, MU | |
| 6 | SU | |
| 7 | SU | |
| 10 | SU, ME, MU | |
| 14 | SU, ME, MU | Later (L) |
| 15 | SU | |
| 17 | SU, ME, MU | |
| 18 | SU | |
| 19 | SU, ME, MU | |
| 20 | SU | |
| 21 | SU, ME, MU | |
| 25 | SU, ME, MU | |

#### S4.2 Amplicon sequencing (16S rRNA gene)

16S rRNA gene amplicons targeting the V3 region were generated using primers 341bf (ACACTCTTTCCCTACACGACGCTCTTCCGATCTCCTACGGGNGGCWGCAG) and 518r (GACTGGAGTTCAGACGTGTGCTCTTCCGATCTWTTACCGCRGCTGCTGG) with Illumina TruSeq adapters.

PCR reactions were performed in 25 µL volumes with 0.04 U/µL Q5 High-Fidelity DNA Polymerase, 1× Q5 reaction buffer, 1× High GC enhancer, 0.25 µM of each primer, 0.2 mM dNTPs, and 100 ng of template DNA. A touchdown PCR protocol was used: initial denaturation at 95°C for 2 min, followed by 4 cycles of 95°C for 9 s, 56-52°C for 40 s, and 72°C for 40 s; then

25 cycles at 95°C for 40 s, 51°C for 40 s, and 72°C for 40 s, with a final extension at 72°C for 5 min and a hold at 8°C. Amplicons were purified using the Clean & Concentrate 5 Kit (Zymo Research, Germany).

#### S4.3 Library preparation and sequencing

Sequencing libraries were prepared from both gDNA (for metagenomics) and 16S rRNA gene amplicons using the NEBNext Ultra II FS DNA Library Prep Kit (New England BioLabs, Germany), following the protocol for DNA input ≥100 ng, applying an incubation time of 14 min for the fragmentation step and 6 PCR cycles for the indexing step. Library quality was assessed with the Agilent High Sensitivity DNA Kit on an Agilent 2100 Bioanalyzer (Agilent Technologies, Germany). Paired-end sequencing was performed on an Illumina NextSeq 550 (Illumina, USA) using a High Output Kit v2.5 (300 Cycles).

#### S4.4 16S rRNA data processing and analysis

The 16S rRNA amplicons were processed using QIIME2 2021.8.0.[49] Raw sequencing reads were trimmed with fastp[50] to remove adapter sequences, and merged with FLASH [51] to have merged reads, which were subsequently imported to QIIME2. From there, the remaining primer sequences were removed with Cutadapt v1.1.4[52] and denoising performed with dada2[53] to generate amplicon sequence variants (ASVs). These ASVs were checked for chimeric reads with help of vsearch[54] and then aligned using mafft [55] (via q2-alignment), and a phylogenetic tree was constructed with fasttree2 [56] (via q2-phylogeny). ASVs were rarefied to 250000 sequences per sample for diversity metric calculations.

Alpha diversity metrics (Shannon [57], Faith's PD [58,59], and ACE [60],) were calculated using the R package phyloseq v1.48.0 [61,62] and picante v.1.8.2 [63],.while beta diversity metrics (Weighted UniFrac [64] and Jaccard distance [65]) were estimated using microbiome v1.26.0 [66] and phyloseq v1.48.0. Taxonomy was assigned to ASVs with the q2-feature-classifier [49] classify-sklearn naive Bayes classifier, using the Silva 138 SSURef NR99 full-length taxonomy database [67] trained on primers 341bf and 518r. The resulting taxonomy was used to compute relative abundances in R [68] with qiime2R v0.99.6 [69].

Differences in community composition among sampling windows were tested by permutational multivariate analysis of variance (PERMANOVA) using adonis2 (vegan v2.7) on Jaccard and weighted UniFrac distance matrices with 999 permutations. Homogeneity of multivariate dispersions was assessed with betadisper and permutest (PERMDISP). Because all samples derive from a single reactor run, these tests evaluate compositional differences between temporal sampling windows and not independently replicated reactor-level effects.

Differentially abundant taxa were identified using ANCOMBC v2.6.0[20,21]. Logarithmic fold differences in feature abundance were calculated using suspension samples 'before' aeration as the reference, with the p-value adjustment method set to 'Holm'.

#### S4.5 Shotgun metagenomic analysis

The raw metagenomic sequences were sequentially filtered to remove adapter and low-quality reads using Trimmomatic v0.36 [70], BBDuk v35.69 (http://jgi.doe.gov/data-and-tools/bb-tools/), and Cutadapt v1.1.4 [52]. High-quality reads were then merged with FLASH v1.2.11 [51] and assembled into long contiguous sequences (contigs) using MEGAHIT v1.2.9 [71]. Coverage profiles were generated with CoverM v0.7.0 [72] by mapping the merged reads to their assembly.

In parallel, coassembly of all the metagenomes from the suspensions belonging to the corresponding condition ('before', 'initial' and 'later') was performed with MEGAHIT to improve metagenome-assembled genomes (MAGs) reconstruction. From there, marker genes were identified using the get_markers function of MDMcleaner v0.8.7 [73] and the predicted SSU and LSU classified with SINA v1.7.2 [67]. The classified sequences were combined with their coverage profiles to calculate relative abundances in R v4.4.1 [68]. In parallel, the assembled metagenomes were annotated with the eggNOG mapper v2.1.6 [74] and combined with their coverage profiles to screen for interesting functional shifts in carbon and nitrogen metabolism based on the KEGG database [75, 76] with R v4.4.1. Gene coverage was estimated by mapping genes back to the assemblies using CoverM v0.7.0 [72]. Gene depths were then normalized for each gene, with further normalization performed in R. To allow comparison of genes with different absolute coverage levels, normalized relative gene abundances were transformed into z-scores (i.e., the deviation of each observation from the gene's mean abundance across samples). This approach shows relative enrichment or depletion patterns within each gene, independent of absolute abundance, and thereby enables direct comparison across samples and conditions. Temporal dynamics of carbon- and nitrogen-related pathways were summarized as line plots showing mean z-scores of all genes within each category across aeration phases. In parallel, detailed gene-level patterns were visualized as heatmaps, where rows represent individual genes and columns represent sampling time points, with color intensity reflecting relative abundance.

**S4.6 MAG reconstruction and analysis**

For metagenome-assembled genomes (MAGs), metagenomic bins were generated using MaxBin2 v2.2.7 [77], MetaBAT v2.12.1 [78], and CONCOCT v1.1.0 [79], followed by refinement with DAS_Tool v1.1.6 [80]. All the reconstructed MAGs were further refined by dereplication with DRep v3.0.0 [81]. The dereplicated MAGs were filtered, retaining only those with >50% completeness and <5% contamination, which were predicted with CheckM2 for further taxonomic classification with GTDB-Tk v2.4.0 [82]. Only MAGs classified up to the genus level were included in further analyses. Relative abundances of MAGs were estimated using CoverM (v.0.7.0) in genome mode by mapping quality-filtered paired-end metagenomic reads to dereplicated MAGs. The relative abundance method was applied, with a minimum read identity threshold of 95% and a minimum alignment coverage of 75% per read. The resulting abundance matrix was normalized by genome length and total mapped reads per sample.

High-quality unclassified MAGs were further processed with the clean function of MDMcleaner v0.8.7 [73]. Predicted proteins were annotated using eggNOG mapper v2.1.6 [74] and analyzed with KEGGDecoder v1.3 for functional analysis, with pathway completeness calculated in R v4.4.1.

### 2. Supplementary Figures

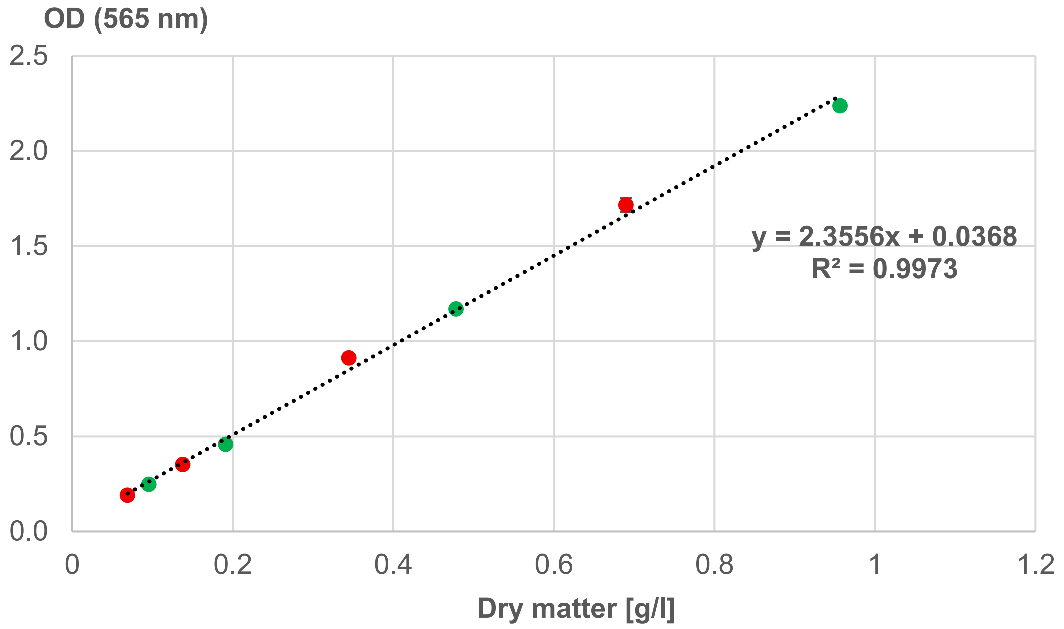


**Figure S1:** Correlation between optical density at 565 nm ($OD_{565}$) and dry biomass concentration of *A. platensis* (triplicate measurements). The linear regression ($R^2$ = 0.9973) demonstrates a strong relationship between $OD_{565}$ and biomass concentration, supporting the use of $OD_{565}$ as a proxy for biomass estimation under the conditions of this study. The calibration is based on parallel measurements of optical density, gravimetric dry weight, and phototrophic pigment-associated absorbance, including wavelengths characteristic for phycocyanin and chlorophyll. Data points represent measurements obtained during the present study, with early (t = −7) and late (t = 23) time points indicated in green and red, respectively.

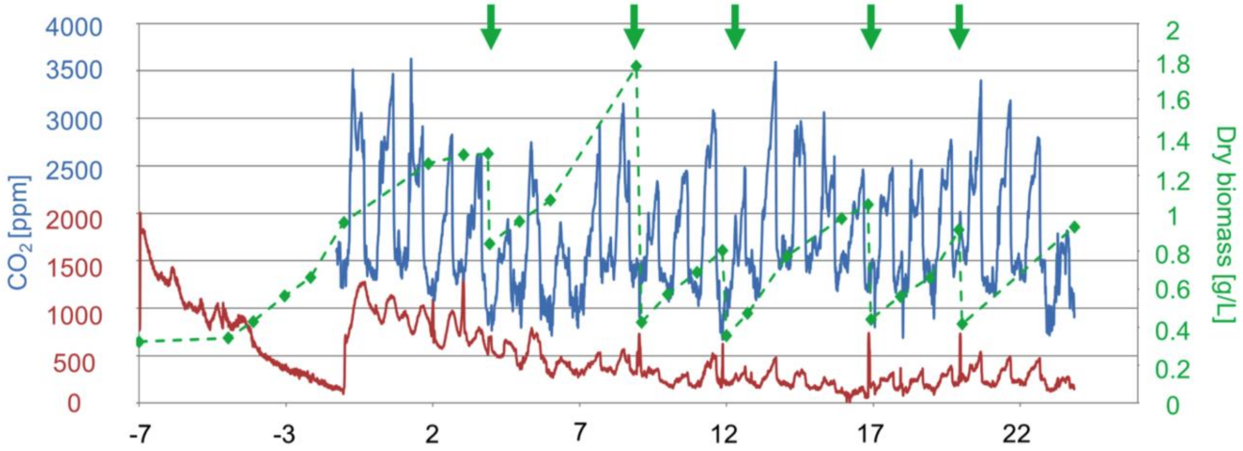


**Figure S2:** Temporal profiles of $CO_2$ concentrations measured at the photobioreactor inlet (Input, blue) and outlet (Output, red), together with suspended biomass concentration estimated from optical density at 565 nm ($OD_{565}$, green) using the experimentally determined $OD_{565}$–dry biomass calibration (Figure S1). During the 7-day acclimation period preceding day 0, the reactor was operated in airlift mode with ambient air (~400 ppm $CO_2$; $NH_3$ not detectable). On day 0, the gas supply was switched to poultry-house exhaust air and maintained continuously thereafter. The pronounced diurnal variation in inlet $CO_2$ concentrations reflects animal housing patterns, with lower daytime concentrations resulting from reduced indoor emissions when the animals accessed the outdoor area via an open flap. Green arrows indicate biomass harvesting events during semi-continuous reactor operation. Harvesting resulted in distinct reductions in suspended biomass concentration, followed by renewed biomass accumulation during subsequent cultivation intervals. Apparent volumetric biomass productivity was calculated from the increase in $OD_{565}$-derived dry biomass concentration between successive harvesting events, normalized to the duration of the respective cultivation interval. Despite repeated harvest-associated reductions, biomass consistently recovered during subsequent cultivation intervals, with and no indication of sustained growth inhibition following harvesting.

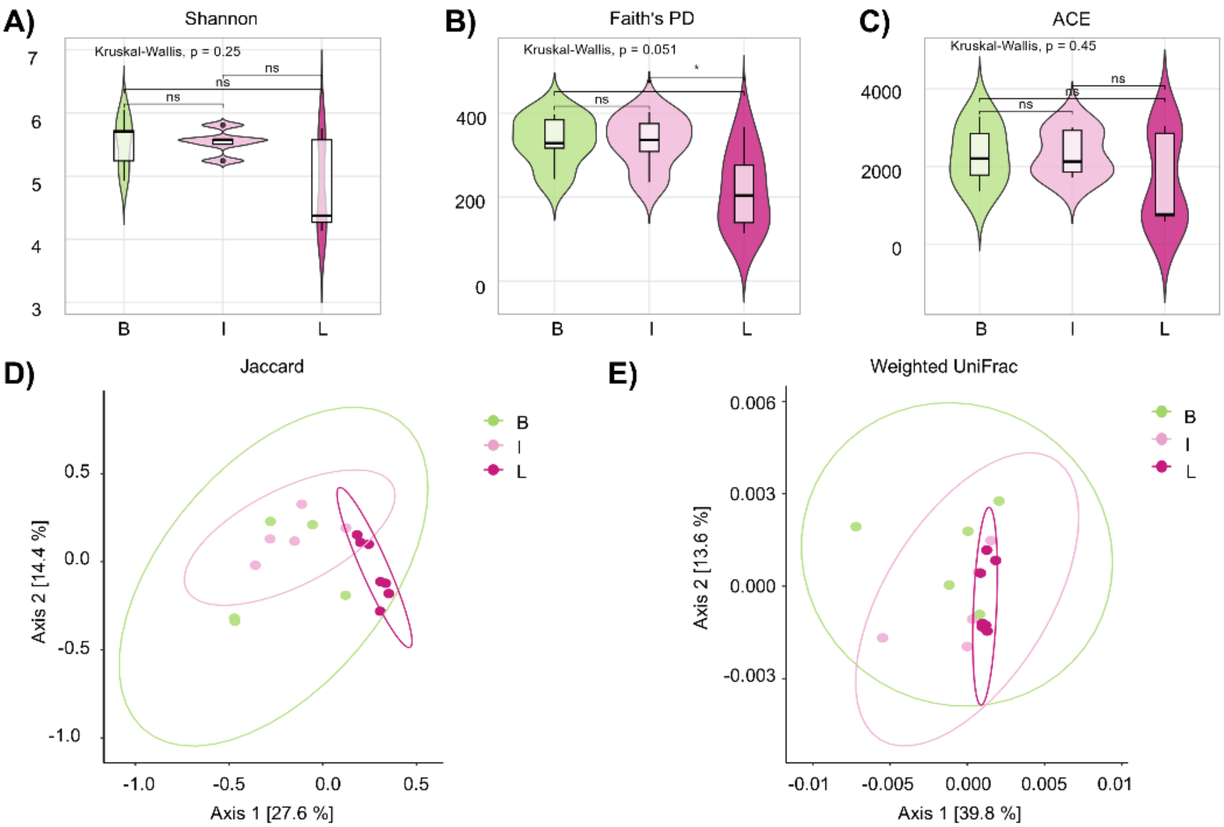


**Figure S3**: Alpha and beta diversity analyses of microbial communities across temporal sampling windows (‘before’, ‘initial’, and ‘later’) during reactor operation. Alpha diversity is represented by (A) Shannon index, (B) Faith’s phylogenetic diversity (PD), and (C) abundance-based coverage estimator (ACE), visualized as violin plots showing the distribution of values across samples. Beta diversity was assessed using (D) Jaccard distance and (E) weighted UniFrac and visualized as principal coordinates analysis (PCoA) plots. Each point represents an individual sample grouped by sampling window (‘before’, green; ‘initial’, light pink; ‘later’, dark pink), with ellipses indicating 95% confidence intervals assuming a multivariate normal distribution. Statistical significance of alpha-diversity comparisons was assessed using Wilcoxon rank-sum tests; non-significant differences are indicated as ns, while * indicates $p < 0.05$. Beta-diversity differences among sampling windows were assessed using PERMANOVA, and homogeneity of multivariate dispersion was evaluated using PERMDISP.

The selected alpha diversity metrics capture complementary aspects of community structure: the Shannon index reflects both abundance and evenness, Faith’s PD summarizes phylogenetic branch lengths, and ACE emphasizes richness with sensitivity to rare taxa. In contrast, beta diversity metrics compare community composition between samples, with weighted UniFrac incorporating phylogenetic relationships and relative abundances, while Jaccard is based solely on presence–absence data.[57-60, 64].

Alpha diversity showed a decreasing temporal trend, with the highest diversity observed in the ‘before’ sampling window and lower values in the ‘initial’ and ‘later’ windows. However, these differences were not statistically significant. Faith’s PD showed the strongest trend between the ‘initial’ and ‘later’ sampling windows ($p = 0.051$), but the difference was not statistically significant.

Beta-diversity ordinations suggested temporal restructuring of the microbial community, with ‘later’ samples showing greater separation from the earlier sampling windows. For Jaccard distances, sampling window was associated with 27.7% of the observed compositional variation (PERMANOVA: $F = 2.88$, $R^2 = 0.277$, $p < 0.001$). However, multivariate dispersion also differed significantly among sampling windows (PERMDISP: $F = 4.92$, $p = 0.024$), and the Jaccard result should therefore be interpreted cautiously. In contrast, weighted UniFrac did not differ significantly among sampling windows (PERMANOVA: $F = 1.55$, $R^2 = 0.171$, $p = 0.209$), with no significant difference in dispersion (PERMDISP: $F = 2.12$, $p = 0.143$).

Together, these analyses indicate temporal variation in community composition within the single reactor campaign, particularly in presence–absence patterns, while alpha diversity and abundance-weighted phylogenetic composition showed no significant overall differences among sampling windows. The considerable within-window variability is consistent with the dynamic, semi-continuous operation of the reactor; however, because elapsed time, exhaust-air exposure, harvesting, and re-fertilization co-varied, their individual contributions to these patterns cannot be resolved.

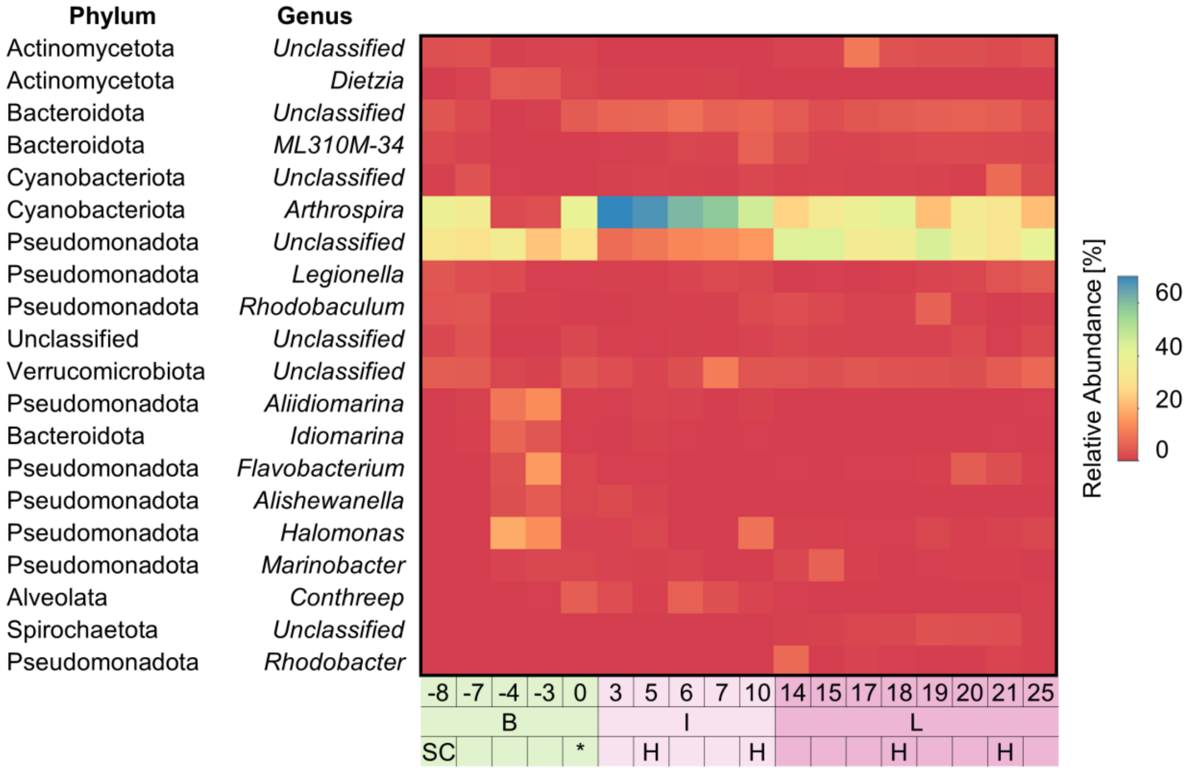


**Figure S4**: Heatmap of the relative abundance of the 20 most abundant genera across all metagenomic samples. Rows represent genera and columns represent sampling time points. Colors indicate relative abundance levels, with low abundance shown in red and high abundance in blue. Samples are grouped by aeration stage (‘before’, B; ‘initial’, I; ‘later’, L) and corresponding sampling days. The heatmap highlights the dominance of *Arthrospira* across the later stages of cultivation, as well as temporal shifts in associated microbial taxa. While some genera persist across all stages, others show stage-specific enrichment patterns, reflecting dynamic community restructuring under aeration and semi-continuous operation, consistent with the progressive restructuring of the microbial community observed in differential abundance analyses (Figure 4).

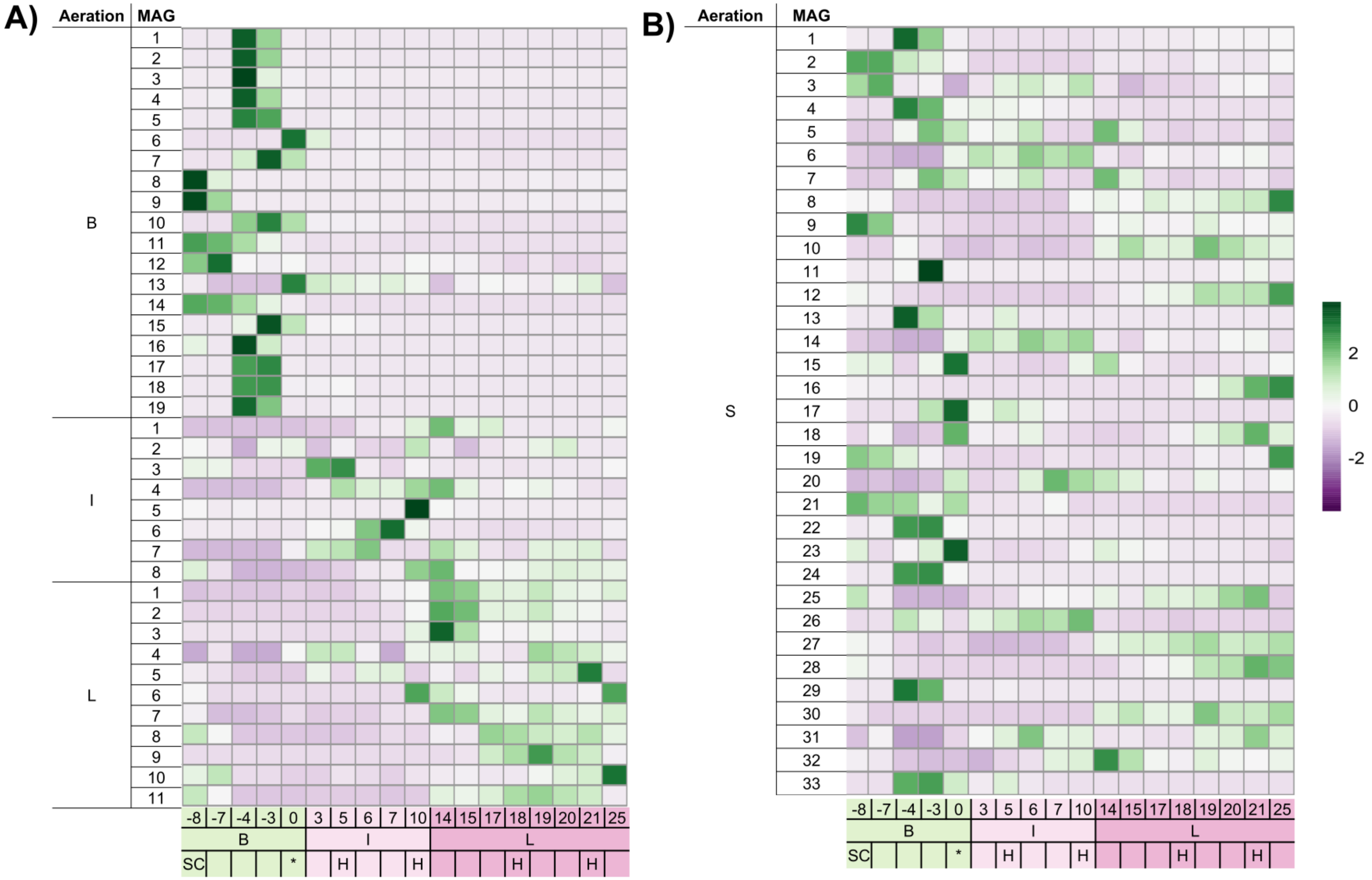


**Figure S5:** Heatmaps showing normalized relative abundances of metagenome-assembled genomes (MAGs) reconstructed from suspension samples. (A) MAGs detected exclusively under specific aeration conditions (‘before’, B; ‘initial’, I; ‘later’, L), highlighting condition-specific taxa. (B) MAGs detected across multiple conditions (shared MAGs), representing persistent members of the microbial community. Colors indicate normalized relative abundance, with purple representing low abundance and green high abundance. Samples are ordered by aeration stage and corresponding time points (days). The separation of condition-specific and shared MAGs illustrates both transient and persistent community members, indicating niche differentiation and adaptive responses to changing environmental conditions during reactor operation.

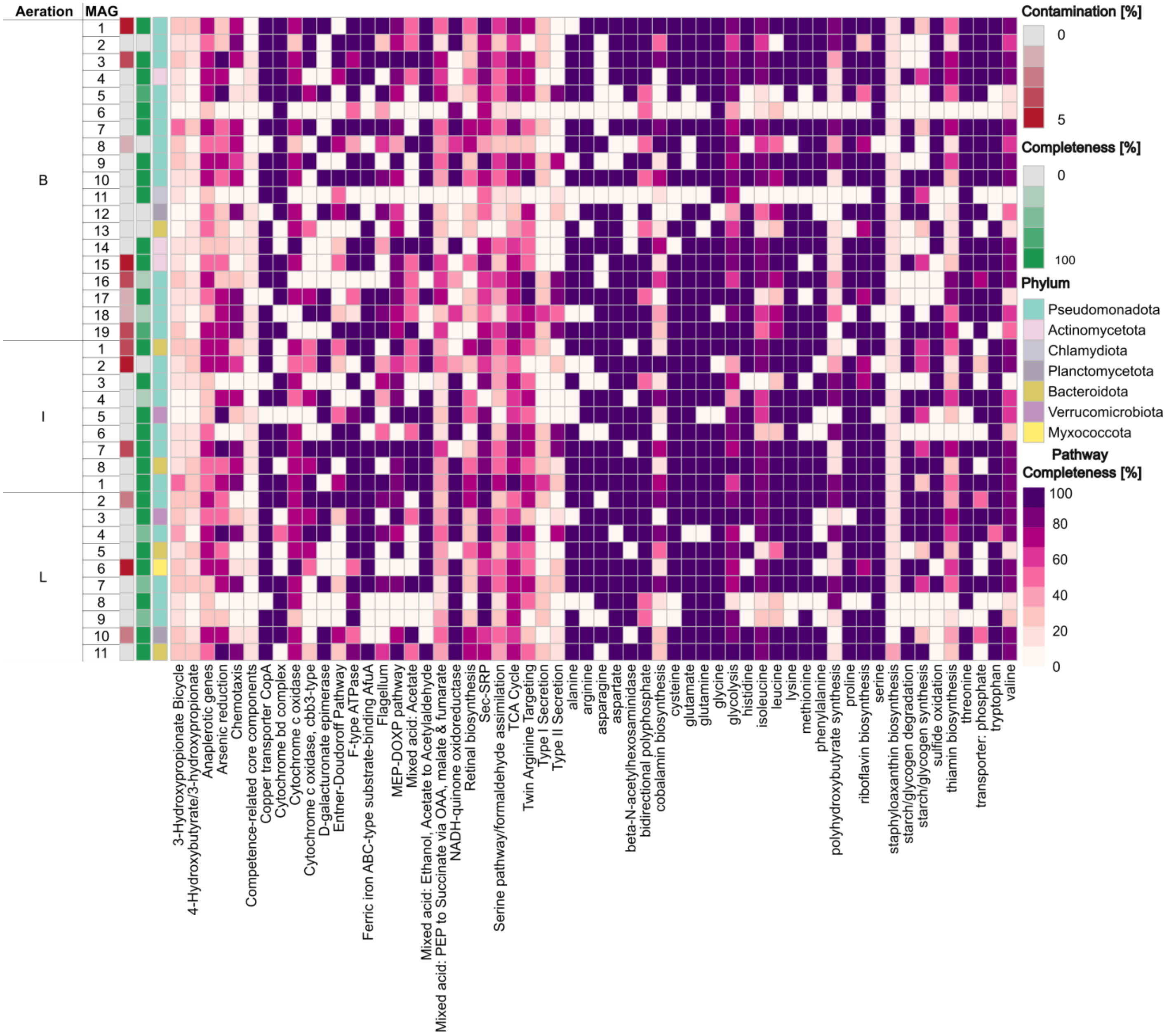
Aeration
MAG
B
I
L
Contamination [%]
0
5
Completeness [%]
0
100
Phylum
Pseudomonadota
Actinomycetota
Chlamydiota
Planctomycetota
Bacteroidota
Verrucomicrobiota
Myxococcota
Pathway Completeness [%]
100
80
60
40
20
0
3-Hydroxypropionate Bicycle
4-Hydroxybutyrate/3-hydroxypropionate
Anaplerotic genes
Arsenic reduction
Chemotaxis
Competence-related core components
Copper transporter CopA
Cytochrome bd complex
Cytochrome c oxidase
Cytochrome c oxidase, cbb3-type
D-galacturonate epimerase
Entner-Doudoroff Pathway
F-type ATPase
Ferric iron ABC-type substrate-binding AfuA
Flagellum
MEP-DOXP pathway
Mixed acid: Acetate
Mixed acid: Ethanol, Acetate to Acetylaldehyde
Mixed acid: PEP to Succinate via OAA, malate & fumarate
NADH-quinone oxidoreductase
Retinal biosynthesis
Sec-SRP
Serine pathway/formaldehyde assimilation
TCA Cycle
Twin Arginine Targeting
Type I Secretion
Type II Secretion
alanine
arginine
asparagine
aspartate
beta-N-acetylhexosaminidase
bidirectional polyphosphate
cobalamin biosynthesis
cysteine
glutamate
glutamine
glycine
glycolysis
histidine
isoleucine
leucine
lysine
methionine
phenylalanine
polyhydroxybutyrate synthesis
proline
riboflavin biosynthesis
serine
staphyloaxanthin biosynthesis
starch/glycogen degradation
starch/glycogen synthesis
sulfide oxidation
thiamin biosynthesis
threonine
transporter: phosphate
tryptophan
valine

**Figure S6:** Functional profiles of condition-specific metagenome-assembled genomes (MAGs) reconstructed under the three aeration stages ('before', B; 'initial', I; 'later', L). The heatmap shows pathway completeness for selected metabolic functions across MAGs, based on genome annotations. Rows represent individual metabolic pathways and columns represent MAGs. Colors indicate pathway completeness, with higher completeness shown in darker shades. Side panels display MAG quality metrics (genome completeness in green and contamination in red) as well as phylum-level taxonomic classification. The distribution of pathway completeness across condition-specific MAGs highlights functional differences between aeration stages, suggesting shifts in metabolic potential associated with changing environmental conditions in the photobioreactor.

A)

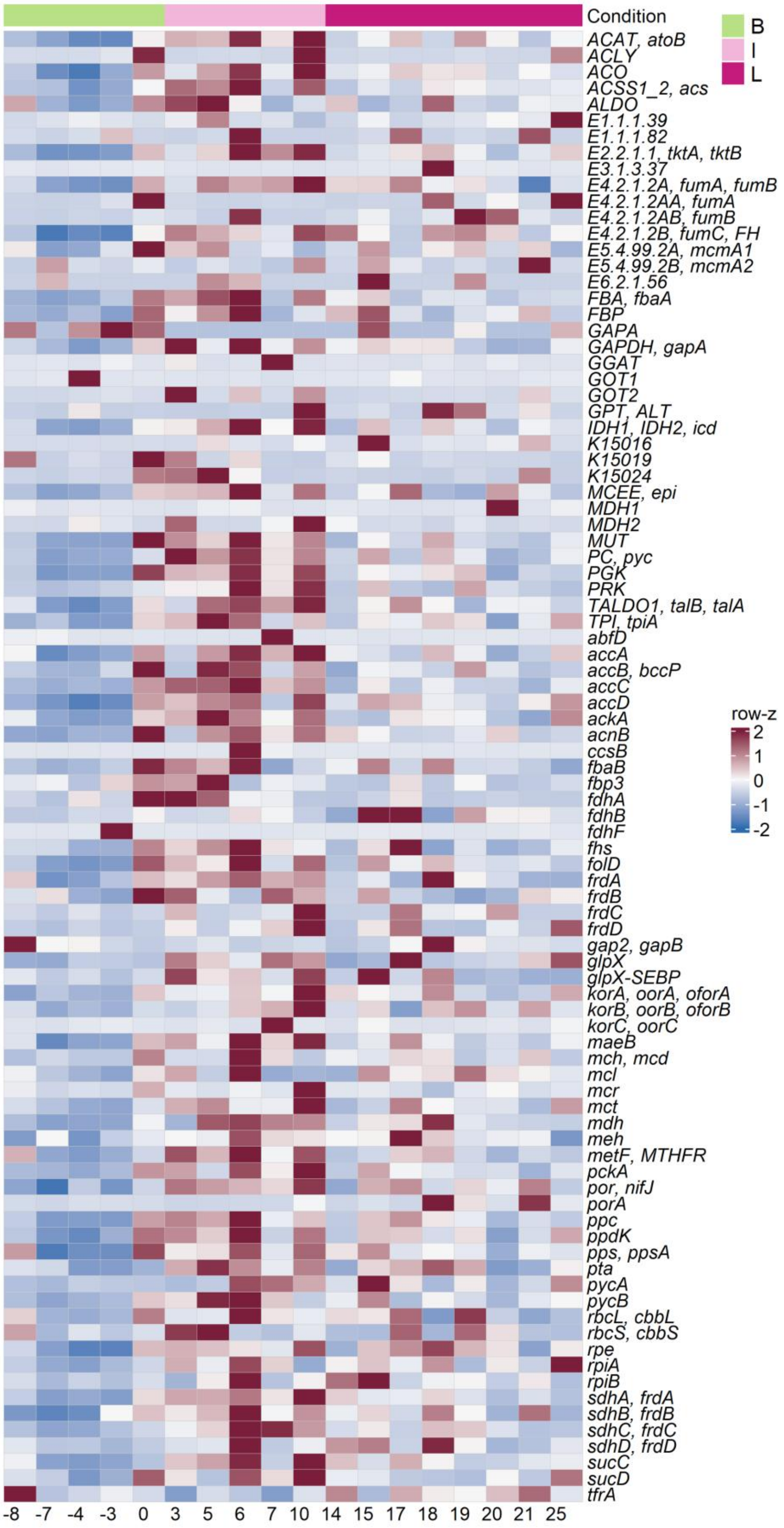

Condition
B
I
L
ACAT, atoB
ACLY
ACO
ACSS1_2, acs
ALDO
E1.1.1.39
E1.1.1.82
E2.2.1.1, tktA, tktB
E3.1.3.37
E4.2.1.2A, fumA, fumB
E4.2.1.2AA, fumA
E4.2.1.2AB, fumB
E4.2.1.2B, fumC, FH
E5.4.99.2A, mcmA1
E5.4.99.2B, mcmA2
E6.2.1.56
FBA, fbaA
FBP
GAPA
GAPDH, gapA
GGAT
GOT1
GOT2
GPT, ALT
IDH1, IDH2, icd
K15016
K15019
K15024
MCEE, epi
MDH1
MDH2
MUT
PC, pyc
PGK
PRK
TALDO1, talB, talA
TPI, tpiA
abfD
accA
accB, bccP
accC
accD
ackA
acnB
ccsB
fbaB
fbp3
fdhA
fdhB
fdhF
fhs
folD
frdA
frdB
frdC
frdD
gap2, gapB
glpX
glpX-SEBP
korA, oorA, oforA
korB, oorB, oforB
korC, oorC
maeB
mch, mcd
mcl
mcr
mct
mdh
meh
metF, MTHFR
pckA
por, nifJ
porA
ppc
ppdK
pps, ppsA
pta
pycA
pycB
rbcL, cbbL
rbcS, cbbS
rpe
rpiA
rpiB
sdhA, frdA
sdhB, frdB
sdhC, frdC
sdhD, frdD
sucC
sucD
tfrA
row-z
2
1
0
-1
-2
-8 -7 -4 -3 0 3 5 6 7 10 14 15 17 18 19 20 21 25

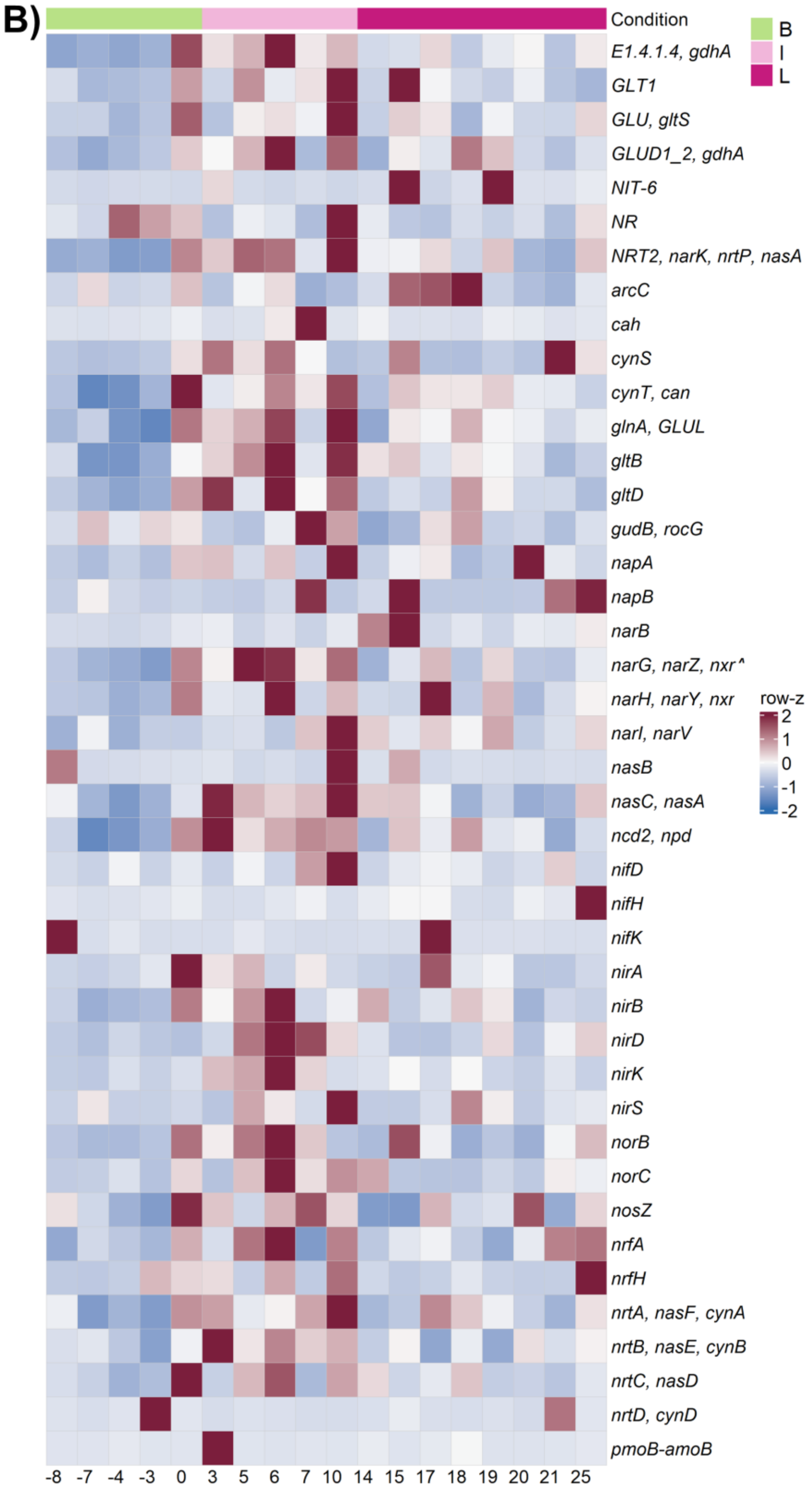


**Figure S7**: Heatmaps showing the relative representation of genes associated with (A) carbon metabolism and (B) nitrogen metabolism across suspension samples collected during the three temporal sampling windows. Rows represent individual genes and columns represent samples ordered by sampling window: before exhaust-air exposure (B, green), initial sampling window under continuous exhaust-air exposure (I, light pink), and later sampling window under continuous exhaust-air exposure (L, dark pink). Gene abundances were normalized across samples for each gene and expressed as row-wise z-scores, with red indicating above-average and blue indicating belowaverage relative abundance for the respective gene. These gene-level patterns provide detailed resolution of changes in the relative representation of genes associated with carbon and nitrogen metabolism, complementing the pathway-level analysis shown in Figure 5.

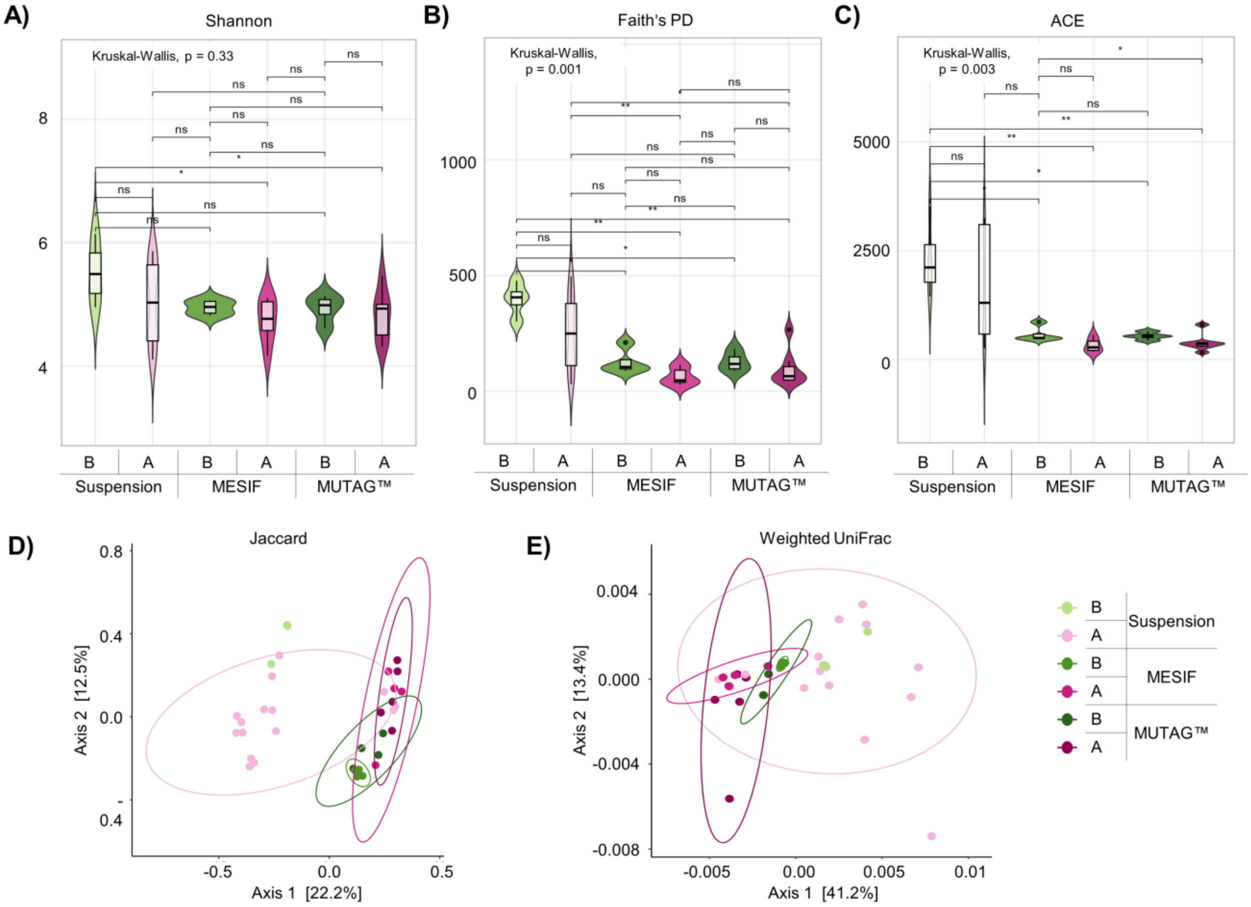


**Figure S8:** Alpha and beta diversity of microbial communities across cultivation matrices and aeration stages. (A-C) Alpha diversity is shown as violin plots for the Shannon index, Faith's phylogenetic diversity (PD), and the abundance-based coverage estimator (ACE). Shannon diversity reflects community evenness and abundance distribution, Faith's PD captures phylogenetic breadth, and ACE emphasizes richness of rare taxa. (D-E) Beta diversity is visualized as principal coordinates analysis (PCoA) based on Jaccard distance (presence/absence) and weighted UniFrac (phylogenetic distance incorporating relative abundance). Each point represents one sample, and ellipses indicate 95% confidence regions. Communities were compared across suspension, MESIF, and MUTAG™ samples before (B, green) and after exhaust-air exposure (A, pink). Statistical significance was assessed using Wilcoxon rank-sum tests (* $p < 0.05$; ns = not significant).

Alpha diversity (A-C) showed a general decline in suspension samples following aeration, with reductions in phylogenetic diversity (Faith's PD, $p < 0.01$) and richness (ACE, $p < 0.01$), while Shannon diversity remained comparatively stable (ns). In contrast, MESIF and MUTAG™-associated communities exhibited only minor changes across aeration stages, indicating that matrix-associated communities were less affected by aeration-induced shifts.

Beta diversity analyses (D, E) revealed clear separation of suspension communities between before and after aeration, whereas matrix-associated communities clustered primarily by aeration stage and remained more similar across time points, indicating greater compositional stability.

Together, these results indicate that cultivation matrices buffer microbiome dynamics and capture more stable surface-associated communities compared to the planktonic phase.

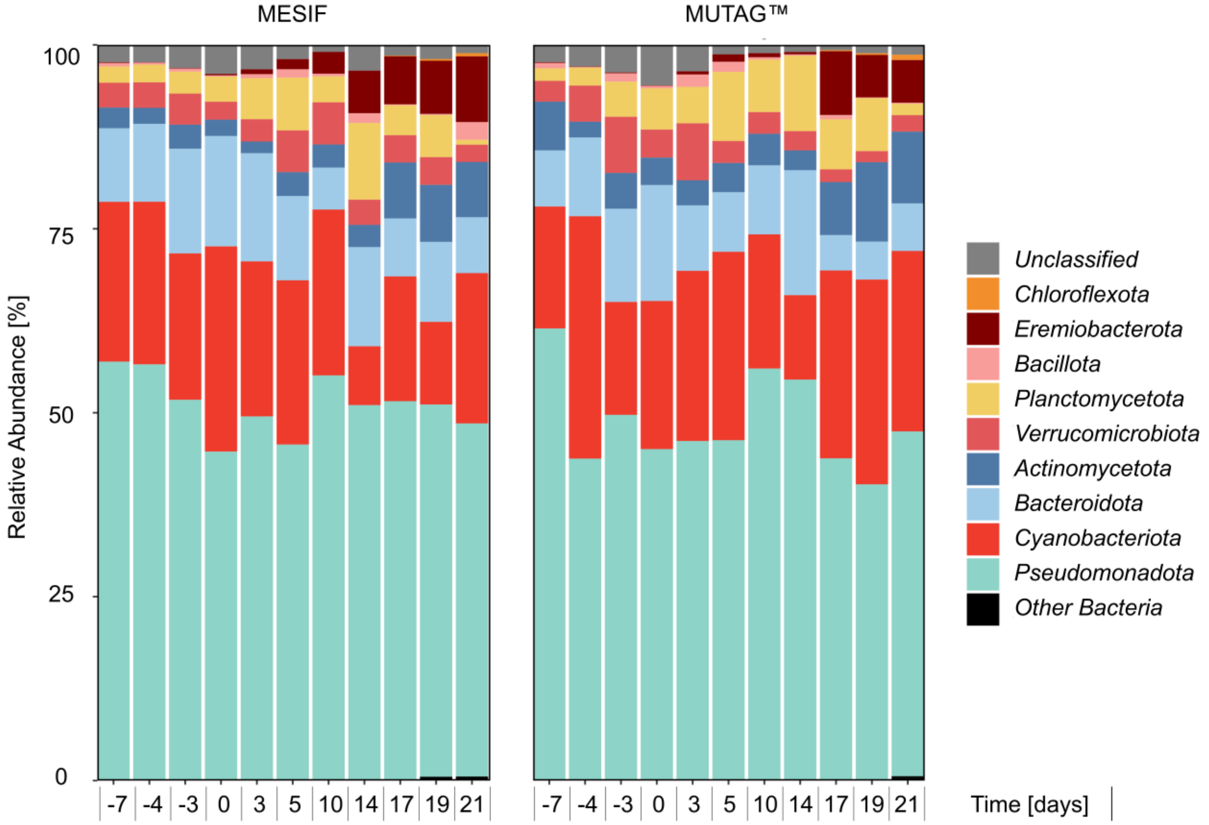


**Figure S9**: Taxonomic classification of 16S rRNA gene amplicons from MESIF and MUTAG™ matrices, showing the 10 most abundant phyla across sampling time points. Taxonomic assignment was performed using the SILVA 138 SSURef NR99 database and visualized as relative abundance bar plots to illustrate temporal shifts in matrix-associated microbial communities. Samples are ordered by time relative to the onset of exhaust-air exposure, allowing comparison of temporal community dynamics between the two carrier types. Overall, MESIF and MUTAG™ showed broadly similar phylum-level community profiles across sampling time points. Members of the candidate phyla *Eremiobacteriota* and *Hydrogenedentota* increased in relative abundance during the later sampling window under continuous exhaust-air exposure (e.g., days 15–17), indicating selective enrichment of previously underexplored microbial lineages within matrix-associated communities. These patterns support the role of cultivation matrices in capturing and enriching microbial diversity that remains underrepresented in suspension-based analyses, consistent with the formation of matrix-associated ecological niches.

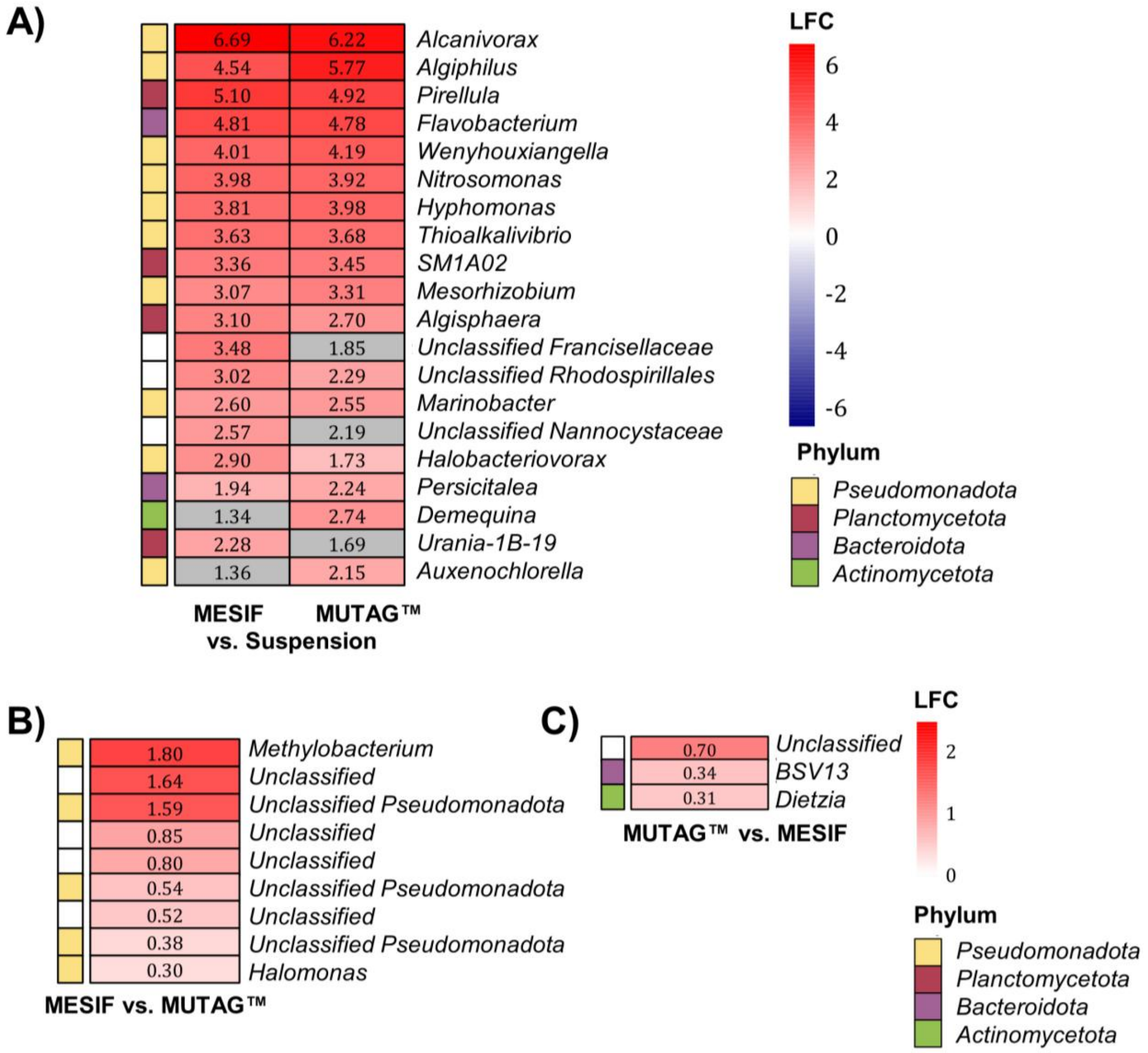


**Figure S10.** Differentially abundant genera in cultivation matrices. Log fold changes (LFC) of significantly enriched taxa are shown for (A) MESIF and MUTAG™ relative to the suspension control, (B) MESIF compared to MUTAG™, and (C) MUTAG™ compared to MESIF. Colors indicate the magnitude and direction of differential abundance (red: enriched; blue: depleted), and values represent LFC. Taxa are annotated at the genus level where possible; otherwise, higher-level classifications are shown as "unclassified." Colored side bars indicate the corresponding phylum of each taxon.

Differential abundance analysis revealed clear taxonomic distinctions between matrix-associated and planktonic communities. Notably, the ammonia-oxidizing genus *Nitrosomonas* was significantly enriched in both MESIF and MUTAG™ relative to the suspension, supporting preferential establishment of nitrifying microorganisms in surface-associated niches.

In addition, several genera with known surface-associated lifestyles, including *Hyphomonas* and *Pirellula*, were consistently enriched in matrix samples, further indicating selective colonization of biofilm-associated environments. A substantial fraction of enriched taxa remained taxonomically unresolved ("unclassified"), highlighting the recovery of previously underexplored microbial diversity within the matrices.

Comparison between carrier types showed that MESIF supported broader and stronger taxonomic enrichment than MUTAG™, with a wider range of genera displaying positive LFC values. These patterns suggest that MESIF provides more diverse microenvironments that facilitate the establishment of specialized microbial communities.

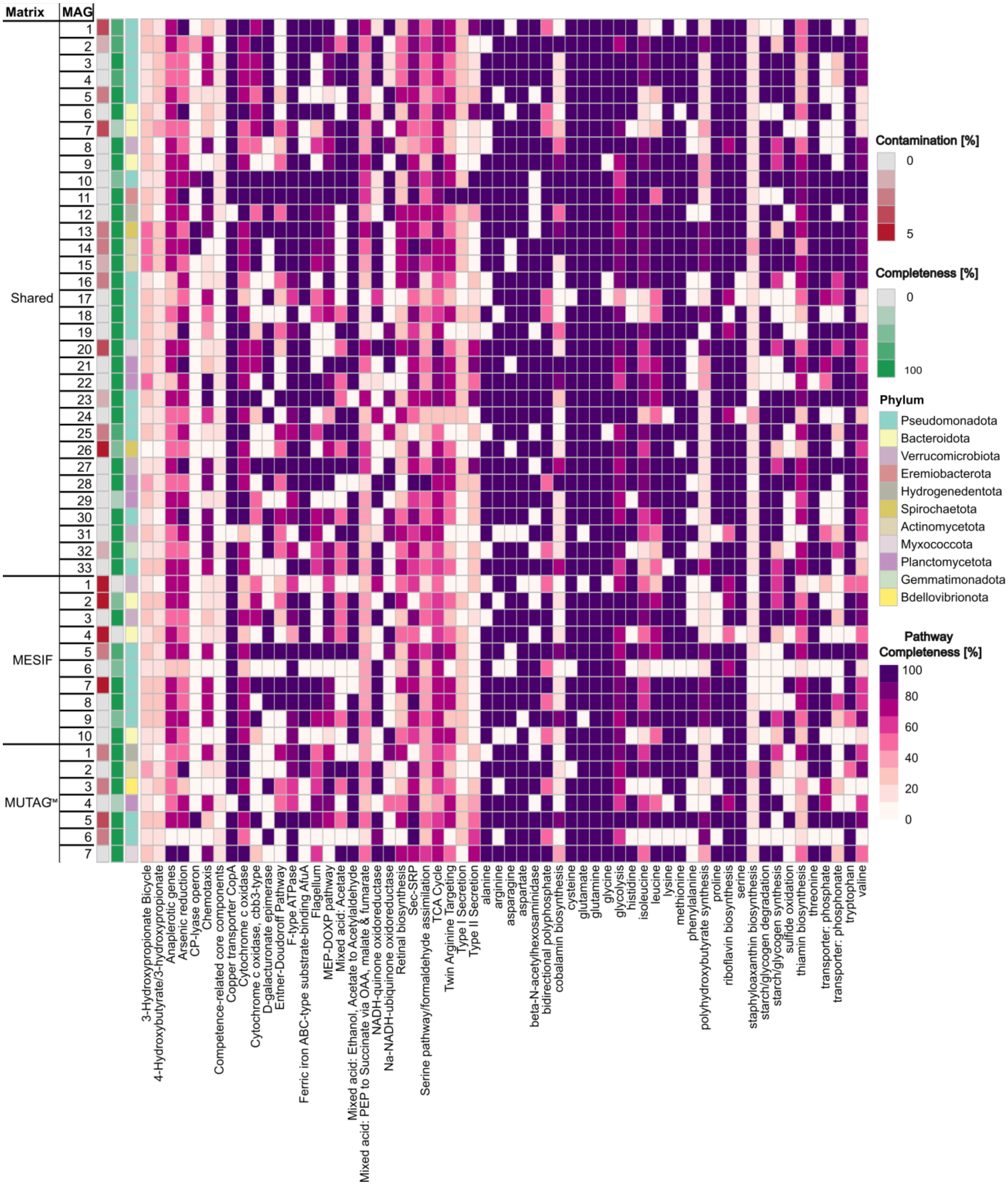


**Figure S11.** Metabolic profiles of metagenome-assembled genomes (MAGs) reconstructed from MESIF and MUTAG™ cultivation matrices. The heatmap displays pathway completeness based on KEGG annotations, with MAGs grouped according to their origin (MESIF, MUTAG™, or shared between both matrices). Side panels indicate MAG quality (completeness in green, contamination in red) and phylum-level taxonomy.

Overall, MAGs from both matrices exhibited broadly similar metabolic profiles, with only minor variation in pathway completeness across samples. No consistent differences in functional potential were observed between MESIF- and MUTAG™-derived MAGs.

These results suggest that functional capabilities are largely conserved across matrix-associated microbial communities, and that differences between matrices are primarily driven by taxonomic composition and relative abundance rather than distinct metabolic repertoires. This pattern is consistent with functional redundancy within the microbiome and supports the interpretation that matrix-specific effects arise from spatial organization and niche differentiation rather than fundamentally different metabolic capacities.

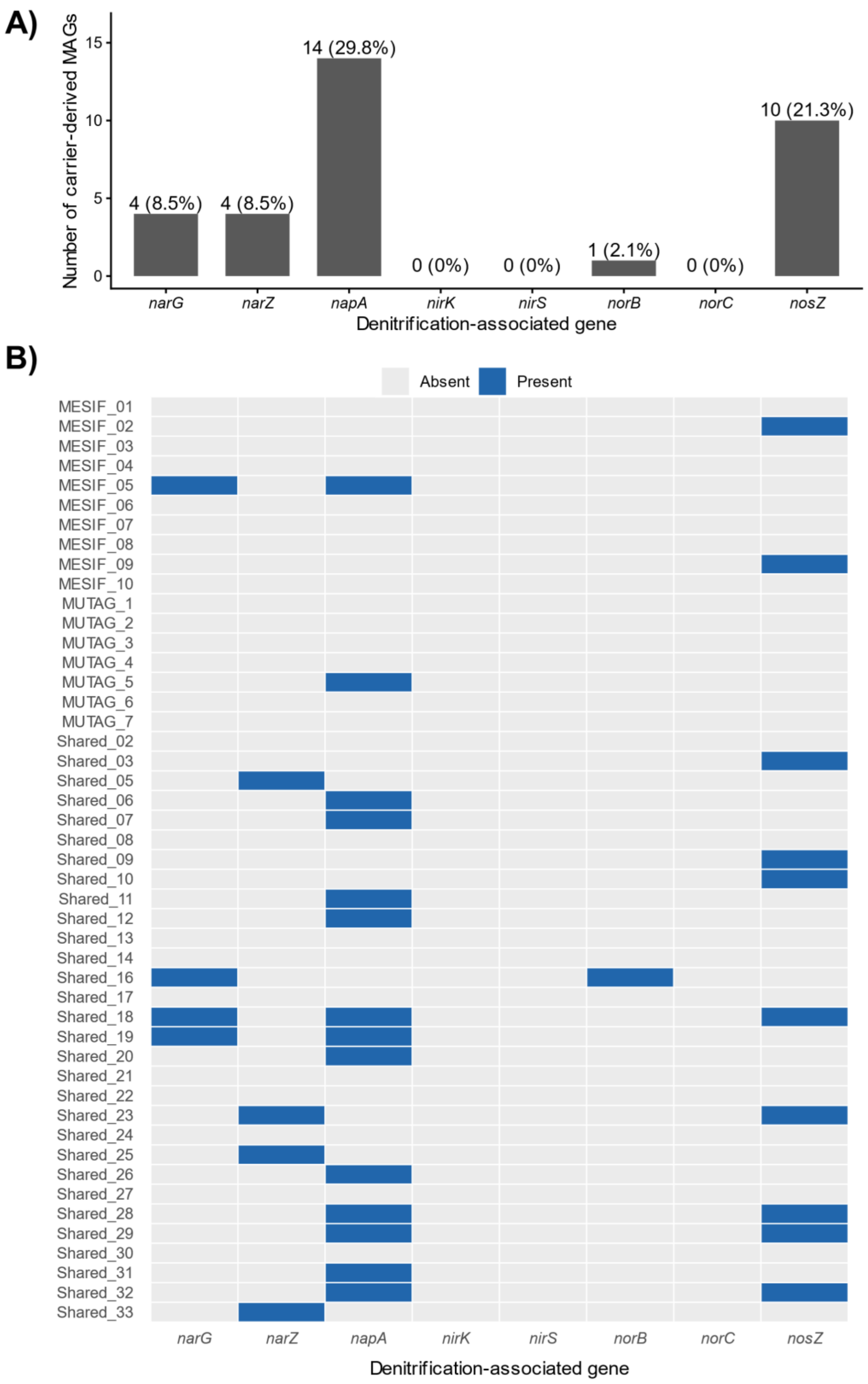


**Figure S12**. Distribution of genes associated with nitrate reduction and denitrification among carrier-derived metagenome-assembled genomes (MAGs). (A) Number and percentage of the 47 MAGs reconstructed from MESIF and MUTAG™ carrier communities containing genes associated with individual steps of nitrate reduction and denitrification. Genes encoding nitrate reductases were comparatively frequent, with *napA* detected in 14 MAGs (29.8%) and *narG* and *narZ* each detected in four MAGs (8.5%). The nitrous oxide reductase gene *nosZ* was detected in 10 MAGs (21.3%), whereas *norB* occurred in one MAG (2.1%); *nirK*, *nirS*, and *norC* were not detected. Overall, 19 of 47 MAGs (40.4%) encoded at least one nitrate reductase gene (*narG*, *narZ*, or *napA*). (B) MAG-resolved presence/absence distribution of the same genes, with MAGs grouped according to their occurrence on MESIF, MUTAG™, or both carrier types. Notably, nitrate-reduction potential was predominantly associated with MAGs shared between the two carrier materials: of the 19 MAGs encoding at least one nitrate reductase gene, 17 were shared between MESIF and MUTAG™, whereas one was MESIF-specific and one MUTAG™-specific. This pattern is consistent with the broadly convergent carrier-associated communities observed across the two materials. No individual MAG contained the complete set of genes examined here for canonical denitrification. Gene presence indicates metabolic potential rather than pathway activity or quantitative nitrogen flux; because MAGs represent incomplete genome reconstructions, absence of individual genes cannot establish their absence from the corresponding organisms. MAG-level gene annotations underlying this analysis are provided in Supplementary Table S8.

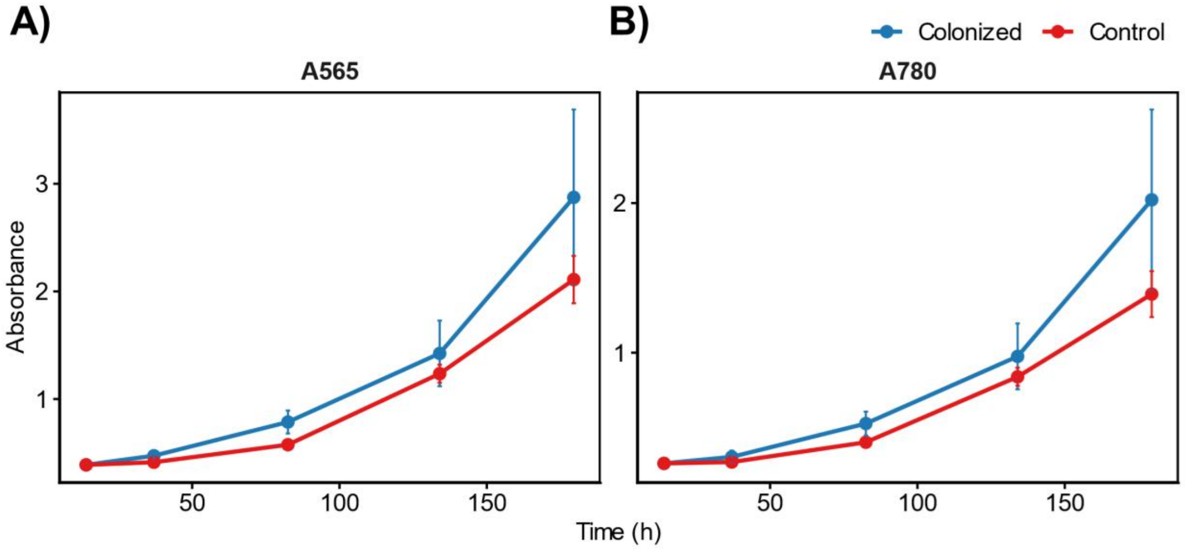


**Figure S13.** Growth dynamics of *Arthrospira maxima* cultures supplemented with pre-colonized MESIF carriers. (A-B) Optical density at 565 nm (A565) and 780 nm (A780) monitored over time (hours) in cultures supplemented with pre-colonized MESIF carriers ("Colonized") and non-colonized controls ("Control"). At both wavelengths, cultures containing pre-colonized MESIFs showed consistently higher absorbance values, indicating enhanced biomass development and faster adaptation to aerated conditions compared to controls. Differences between treatments increased over time, suggesting a sustained growth advantage associated with pre-established microbial communities on the carriers, consistent with enhanced culture establishment. Error bars represent standard deviation of two independent biological replicates

### 3. Supporting Tables

**Table S2:** Differentially abundant taxa identified with ANCOM-BC. Log-fold changes (LFC) indicate enrichment or depletion at the genus level. Significance is reported as true (T) for statistically different taxa and false (F) for non-significant taxa.

| Taxonomy | | Log-Fold Change | | Differential Significance | |
|---|---|---|---|---|---|
| **Phylum** | **Genus** | **I** | **L** | **I** | **L** |
| *Armatimonadota* | *c__uncultured* | 4.27 | 5.46 | T | T |
| *Pseudomonadota* | *g__Roseococcus* | 3.97 | 3.84 | T | T |
| *Pseudomonadota* | *g__Rubellimicrobium* | 2.32 | 5.45 | F | T |
| *Pseudomonadota* | *g__Candidatus Alysiosphaera* | 2.75 | 3.71 | T | T |
| *Pseudomonadota* | *g__Zoogloea* | 5.84 | 0.61 | T | F |
| *Spirochaetota* | *g__Salinispira* | 0.95 | 5.33 | T | T |
| *Verrucomicrobiota* | *g__ADurb,Bin063-1* | 1.87 | 3.97 | F | T |
| *Pseudomonadota* | *g__Kaistia* | 5.05 | -0.37 | T | F |
| *Pseudomonadota* | *f__uncultured Desulfuromonadales* | -0.04 | 4.70 | F | T |
| *Bacteroidota* | *g__Fontibacter* | 2.24 | 2.22 | T | T |
| *Actinomycetota* | *f__Demequinaceae* | 1.93 | 2.51 | T | T |
| *Pseudomonadota* | *g__Candidatus Trichorickettsia* | 3.83 | 0.31 | T | T |
| *Sumerlaeota* | *c__uncultured* | -0.09 | 4.15 | F | T |
| *Pseudomonadota* | *g__Candidatus Megaira* | 0.88 | 3.14 | T | T |
| *Verrucomicrobiota* | *c__Verrucomicrobiae* | 0.92 | 3.01 | T | T |
| *Pseudomonadota* | *f__Caulobacteraceae* | 4.37 | -0.54 | T | F |
| *Bacteroidota* | *g__Flavo* | 2.53 | 1.25 | T | T |
| *Actinomycetota* | *f__ HN-HFO4* | 2.57 | 1.18 | T | T |
| *Latescibacteria* | *c__uncultured proteo* | 1.94 | 1.75 | T | T |
| *Bacillota* | *f__uncultured Selenomonadales* | 1.70 | 1.86 | T | T |
| *Pseudomonadota* | *f__Pseudomonadaceae* | 5.61 | -2.07 | T | T |
| *Bacillota* | *f__Clostridiaceae 3* | 1.59 | 1.78 | F | T |
| *Verrucomicrobiota* | *o__Opitutales* | 0.71 | 2.61 | F | T |
| *Cyanobacteriota* | *c__Oxyphotobacteria* | 0.34 | 2.93 | F | T |
| *Armatimonadota* | *c__uncultured* | 2.40 | 0.76 | T | F |
| *Bacillota* | *g__Pelotomaculum* | 2.94 | 0.20 | T | F |
| *Sumerlaeota* | *c__metagenome* | 2.91 | 0.20 | T | F |
| *Bacteroidota* | *f__Bacteroidetes vadinHA17* | 1.70 | 1.38 | T | T |
| *Pseudomonadota* | *g__Thauera* | 0.57 | 2.48 | F | T |
| *Bacteroidota* | *g__Taibaiella* | 2.85 | 0.20 | T | F |
| *Verrucomicrobiota* | *g__Verruc-01* | 1.15 | 1.77 | T | T |
| *Spirochaetota* | *g__Leptospira* | 0.25 | 2.66 | T | T |
| *Actinomycetota* | *g__Solirubrobacter* | 1.70 | 1.12 | T | T |
| *Pseudomonadota* | *f__Methylophilaceae* | 0.61 | 2.18 | T | T |
| *Armatimonadota* | *c__uncultured* | 0.65 | 2.10 | F | T |
| *Pseudomonadota* | *g__Bosea* | 3.24 | -0.49 | T | F |

| | | | | | |
|---|---|---|---|---|---|
| Bacillota | f__Peptostreptococcaceae | 2.61 | 0.12 | T | T |
| Pseudomonadota | g__Auxenochlorella pyrenoidosa | 0.59 | 1.99 | T | T |
| Armatimonadota | c__uncultured | 0.92 | 1.65 | F | T |
| Pseudomonadota | g__Stenotrophomonas | 3.95 | -1.41 | F | T |
| Pseudomonadota | g__Methylomonas | 0.36 | 2.16 | F | T |
| Verrucomicrobiota | o__Verrucomicrobiales | -0.99 | 3.49 | F | T |
| Bacillota | g__Cryptanaerobacter | 0.63 | 1.82 | F | T |
| Pseudomonadota | f__Desulfobulbaceae | -0.08 | 2.49 | F | T |
| Bacillota | g__Ruminococcaceae UCG-014 | 1.98 | 0.37 | T | F |
| Pseudomonadota | f__Diplorickettsiaceae | 1.87 | 0.39 | T | F |
| Armatimonadota | c__uncultured | 0.79 | 1.41 | F | T |
| Spirochaetota | g__RBG-16-49-21 | -0.08 | 2.26 | F | T |
| Pseudomonadota | g__Desulfoprunum | -0.04 | 2.14 | F | T |
| Actinomycetota | f__Micrococcaceae | 2.90 | -0.81 | T | F |
| Armatimonadota | c__uncultured | 1.42 | 0.66 | T | T |
| Pseudomonadota | g__Geobacter | 0.00 | 2.07 | F | T |
| Sumerlaeota | c__uncultured | 0.76 | 1.29 | F | T |
| Cyanobacteriota | f__Nostocaceae | 1.32 | 0.72 | T | T |
| Bacteroidota | g__Terrimonas | 1.65 | 0.37 | T | T |
| Bacteroidota | g__Blvii28 wastewater-sludge group | -0.08 | 2.10 | F | T |
| Bacillota | o__uncultured mle1-9 | -0.08 | 2.08 | F | T |
| Bacteroidota | g__Pedobacter | 1.80 | 0.20 | T | F |
| Pseudomonadota | g__OM27 clade | 1.88 | 0.13 | T | T |
| Pseudomonadota | f__Mitochondria | 1.67 | 0.34 | T | F |
| Pseudomonadota | g__Desulforegula | -0.08 | 2.08 | F | T |
| Pseudomonadota | g__Pseudoxanthomonas | 2.95 | -0.97 | T | F |
| Armatimonadota | c__uncultured | -0.08 | 2.05 | F | T |
| Patescibacteria | f__uncultured gamma proteo | 0.51 | 1.46 | F | T |
| Gemmatimonadetes | o__uncultured actino | 0.70 | 1.25 | F | T |
| Pseudomonadota | g__Ralstonia | 1.74 | 0.20 | T | F |
| Pseudomonadota | g__Aqua | 0.66 | 1.27 | F | T |
| Sumerlaeota | c__uncultured | 0.48 | 1.43 | F | T |
| Cyanobacteriota | c__Sericytochromatia | 2.72 | -0.84 | T | F |
| Pseudomonadota | f__Acetobacteraceae | 1.60 | 0.26 | T | T |
| Pseudomonadota | g__Alcanivorax | 0.28 | 1.55 | F | T |
| Bacteroidota | g__Mongoliitalea | 2.91 | -1.09 | T | T |
| Acidobacteriota | c__Subgroup 17 | 1.53 | 0.28 | T | T |
| Bacillota | g__[Eu] coprostanoligenes group | 3.13 | -1.33 | F | T |
| Pseudomonadota | g__[Desulfo] catecholicum group | -0.08 | 1.87 | F | T |
| Bacillota | f__Paenibacillaceae | -0.94 | 2.72 | F | T |
| Cyanobacteriota | o__uncultured cyano | 2.24 | -0.47 | T | F |
| Deferribacterota | g__Seleniivibrio | 4.01 | -2.23 | T | F |
| Bacillota | g__Acidaminobacter | 0.84 | 0.92 | T | T |

| | | | | | |
|---|---|---|---|---|---|
| *Bacillota* | *g__Soehngenia* | -0.08 | 1.81 | F | T |
| *Sumerlaeota* | *c__uncultured soil* | 0.54 | 1.17 | F | T |
| *Sumerlaeota* | *c__uncultured soil* | 0.54 | 1.17 | T | T |
| *Sumerlaeota* | *c__uncultured soil* | 0.54 | 1.17 | F | T |
| *Sumerlaeota* | *c__uncultured soil* | 0.54 | 1.17 | T | T |
| *Armatimonadota* | *c__uncultured* | -0.08 | 1.76 | F | T |
| *Bacteroidota* | *g__GWE2-42-42* | -0.08 | 1.73 | F | T |
| *Sumerlaeota* | *c__uncultured* | 0.14 | 1.46 | F | T |
| *Pseudomonadota* | *g__Nannocystis* | 1.39 | 0.19 | T | T |
| *Sumerlaeota* | *c__metagenome* | 1.30 | 0.26 | T | T |
| *Acidobacteriota* | *c__Subgroup 18* | 0.47 | 1.08 | F | T |
| *Pseudomonadota* | *f__uncultured Anaeromyxobacter sp,* | 0.65 | 0.88 | F | T |
| *Pseudomonadota* | *g__uncultured Rhodospirillaceae* | -0.08 | 1.60 | F | T |
| *Bacteroidota* | *g__Roseimarinus* | -0.08 | 1.59 | F | T |
| *Bacteroidota* | *f__Chlorobi OLB5* | 1.05 | 0.46 | T | T |
| *Pseudomonadota* | *o__NB1-j* | 1.42 | 0.07 | T | T |
| *Bacillota* | *g__Desulfuribacillus* | -0.22 | 1.69 | F | T |
| *Pseudomonadota* | *g__Wenzhouxiangella* | 0.28 | 1.18 | F | T |
| *Gemmatimonadetes* | *o__uncultured actino* | 1.24 | 0.20 | F | T |
| *Sumerlaeota* | *c__uncultured soil* | -0.08 | 1.51 | F | T |
| *Sumerlaeota* | *c__uncultured soil* | -0.08 | 1.51 | F | T |
| *Sumerlaeota* | *c__uncultured soil* | -0.08 | 1.51 | F | T |
| *Sumerlaeota* | *c__uncultured soil* | -0.08 | 1.51 | F | T |
| *Pseudomonadota* | *g__Pseudorhodoplanes* | 1.38 | 0.03 | T | T |
| *Planctomycetota* | | 1.05 | 0.35 | T | F |
| *Synergistetes* | *g__JGI-0000079-D21* | -0.08 | 1.48 | F | T |
| *Bacillota* | *g__Salipaludibacillus* | 1.66 | -0.28 | T | F |
| *Actinomycetota* | *f__uncultured Acidimicrobiales* | 1.29 | 0.07 | T | T |
| *Armatimonadota* | *c__uncultured* | -0.08 | 1.43 | T | F |
| *Acidobacteriota* | | 0.80 | 0.54 | T | T |
| *Actinomycetota* | *g__uncultured Gaiella sp,* | 1.46 | -0.14 | T | T |
| *Chlamydiota* | *g__Neochlamydia* | 0.76 | 0.55 | T | T |
| *Patescibacteria* | *c__Gracilibacteria* | -0.08 | 1.36 | F | T |
| *Armatimonadota* | *c__uncultured* | 0.92 | 0.35 | T | T |
| *Bacillota* | *g__Alkaliphilus* | -0.86 | 2.08 | F | T |
| *Nitrospirae* | *f__uncultured Sphingobacteriales* | -0.32 | 1.54 | F | T |
| *Armatimonadota* | *c__uncultured* | -0.08 | 1.29 | F | T |
| *Sumerlaeota* | *c__metagenome* | -0.08 | 1.28 | F | T |
| *Rokubacteria* | *g__Sh765B-TzT-35* | 0.23 | 0.97 | F | T |
| *Pseudomonadota* | *f__KF-JG30-B3* | 0.47 | 0.72 | F | T |
| *Pseudomonadota* | *g__uncultured alpha proteo* | 0.83 | 0.35 | T | T |
| *Gemmatimonadetes* | *o__uncultured actino* | 0.92 | 0.26 | T | F |
| *Pseudomonadota* | *g__Nordella* | 1.20 | -0.06 | T | T |

| | | | | | |
|---|---|---|---|---|---|
| *Chloroflexota* | *g__UTCFX1* | 0.68 | 0.46 | T | T |
| *Sumerlaeota* | *c__uncultured* | 0.63 | 0.51 | F | T |
| *Bacillota* | *f__Syntrophomonadaceae* | 2.11 | -0.99 | T | F |
| *Pseudomonadota* | *g__Sva0081 sediment group* | 0.58 | 0.53 | T | T |
| *Sumerlaeota* | *c__uncultured* | 1.35 | -0.24 | F | T |
| *Latescibacteria* | *c__uncultured prokaryote* | 0.56 | 0.50 | F | T |
| *Sumerlaeota* | *c__uncultured* | 0.42 | 0.63 | T | T |
| *Bacteroidota* | *g__Ignavi* | 0.61 | 0.40 | T | T |
| *Gemmatimonadetes* | *o__uncultured actino* | 0.79 | 0.20 | T | F |
| *Sumerlaeota* | *c__metagenome* | -0.08 | 1.05 | F | T |
| *Patescibacteria* | *o__uncultured microorganism* | -0.08 | 1.03 | T | T |
| *Pseudomonadota* | *f__Polyangiaceae* | 0.84 | 0.10 | T | T |
| *Sumerlaeota* | *c__uncultured* | 0.50 | 0.44 | F | T |
| *Sumerlaeota* | *c__uncultured* | 0.50 | 0.44 | T | T |
| *Sumerlaeota* | *c__uncultured* | 0.50 | 0.44 | F | T |
| *Sumerlaeota* | *c__uncultured* | 0.50 | 0.44 | T | T |
| *Sumerlaeota* | *c__metagenome* | -0.08 | 1.01 | F | T |
| *Chloroflexota* | *o__S085* | 1.23 | -0.30 | T | T |
| *WS4* | *c__uncultured organism* | 1.48 | -0.55 | T | F |
| *Chloroflexota* | *o__uncultured Chloroflexi* | 0.98 | -0.06 | T | T |
| *Chloroflexota* | *o__uncultured Chloroflexi* | 0.98 | -0.06 | T | T |
| *Chloroflexota* | *o__uncultured Chloroflexi* | 0.98 | -0.06 | T | T |
| *Chloroflexota* | *o__uncultured Chloroflexi* | 0.98 | -0.06 | T | T |
| *Sumerlaeota* | *c__uncultured soil* | 0.42 | 0.50 | F | T |
| *WS2* | *c__uncultured Firmicutes* | -0.08 | 0.99 | F | T |
| *Bacteroidota* | *g__Flavihumibacter* | 0.64 | 0.26 | T | T |
| *Sumerlaeota* | *c__uncultured* | -0.08 | 0.97 | T | T |
| *Armatimonadota* | *c__uncultured* | 1.03 | -0.14 | T | T |
| *Synergistetes* | *f__Synergistaceae* | 1.36 | -0.48 | T | F |
| *Actinomycetota* | *c__Coriobacteriia* | 1.82 | -0.94 | F | T |
| *Sumerlaeota* | *c__metagenome* | -0.08 | 0.95 | T | T |
| *Bacillota* | *f__Gracilibacteraceae* | 0.21 | 0.65 | F | T |
| *Acidobacteriota* | *c__Subgroup 5* | 0.44 | 0.42 | T | T |
| *Eremiobacterota* | *c__hydrothermal vent metagenome* | -1.62 | 2.47 | T | F |
| *Pseudomonadota* | *f__Magnetospirillaceae* | -0.48 | 1.32 | F | T |
| *Pseudomonadota* | *g__Desulfococcus* | -0.24 | 1.08 | F | T |
| *Armatimonadota* | *c__uncultured* | 0.96 | -0.13 | T | F |
| *Planctomycetota* | *g__Candidatus Brocadia* | 0.34 | 0.49 | F | T |
| *Sumerlaeota* | *c__metagenome* | 0.42 | 0.40 | F | T |
| *Acidobacteriota* | *o__uncultured Holophagae* | -0.02 | 0.83 | F | T |
| *Chloroflexota* | *o__uncultured Chloroflexi* | -0.08 | 0.89 | F | T |
| *Sumerlaeota* | *c__uncultured soil* | -0.09 | 0.89 | F | T |
| *Sumerlaeota* | *c__uncultured soil* | -0.09 | 0.89 | F | T |

| | | | | | |
|---|---|---|---|---|---|
| *Sumerlaeota* | *c__uncultured soil* | -0.09 | 0.89 | F | T |
| *Sumerlaeota* | *c__uncultured soil* | -0.09 | 0.89 | F | T |
| *Acidobacteriota* | *g__Luteitalea* | 0.59 | 0.19 | T | T |
| *Armatimonadota* | *f__uncultured Lutispora sp,* | 0.68 | 0.07 | T | T |
| *Sumerlaeota* | *c__metagenome* | -0.10 | 0.82 | F | T |
| *Sumerlaeota* | *c__uncultured* | -0.12 | 0.84 | F | T |
| *Sumerlaeota* | *c__uncultured* | 0.28 | 0.44 | F | T |
| *Armatimonadota* | *c__uncultured* | 0.61 | 0.10 | T | T |
| *Actinomycetota* | *f__Nocardioidaceae* | 0.54 | 0.17 | T | T |
| *Sumerlaeota* | *c__uncultured* | 0.50 | 0.19 | T | T |
| *Sumerlaeota* | *c__uncultured* | 0.50 | 0.19 | F | T |
| *Sumerlaeota* | *c__uncultured* | 0.50 | 0.19 | T | T |
| *Sumerlaeota* | *c__uncultured* | 0.50 | 0.19 | F | T |
| *Pseudomonadota* | *g__MND1* | 0.59 | 0.10 | T | T |
| *Chloroflexota* | *c__Dehalococcoidia* | 0.51 | 0.18 | T | T |
| *Rokubacteria* | *o__Rokubacteriales* | 0.63 | 0.03 | T | T |
| *Pseudomonadota* | *o__Sva0485* | -0.14 | 0.80 | F | T |
| *Sumerlaeota* | *c__uncultured* | 0.46 | 0.20 | F | T |
| *Latescibacteria* | *c__uncultured prokaryote* | -0.05 | 0.71 | F | T |
| *Chloroflexota* | *o__uncultured Chloroflexi* | 0.56 | 0.09 | T | T |
| *Chloroflexota* | *o__uncultured Chloroflexi* | 0.56 | 0.09 | T | T |
| *Chloroflexota* | *o__uncultured Chloroflexi* | 0.56 | 0.09 | T | T |
| *Chloroflexota* | *o__uncultured Chloroflexi* | 0.56 | 0.09 | T | T |
| *Acidobacteriota* | *o__Acidobacteria WX90* | 0.49 | 0.14 | T | T |
| *Bacteroidota* | *f__AKYH767* | 0.29 | 0.34 | T | T |
| *Pseudomonadota* | *g__uncultured Cystobacterineae* | 0.66 | -0.04 | T | T |
| *Latescibacteria* | *c__uncultured soil PBS-II-5* | -0.06 | 0.67 | F | T |
| *Sumerlaeota* | *c__metagenome* | 0.10 | 0.49 | F | T |
| *Acidobacteriota* | *o__uncultured AD106-G4* | 0.45 | 0.12 | T | T |
| *Gemmatimonadetes* | *o__uncultured Gemmatimonadetes* | 0.65 | -0.08 | T | F |
| *Bacillota* | *g__Brevibacillus* | 0.99 | -0.43 | T | F |
| *Latescibacteria* | *c__uncultured prokaryote* | 0.36 | 0.20 | F | T |
| *Bacillota* | *g__Sporosarcina* | 0.19 | 0.36 | T | T |
| *Sumerlaeota* | *c__uncultured soil* | -0.52 | 1.07 | F | T |
| *Sumerlaeota* | *c__uncultured soil* | -0.52 | 1.07 | F | T |
| *Sumerlaeota* | *c__uncultured soil* | -0.52 | 1.07 | F | T |
| *Sumerlaeota* | *c__uncultured soil* | -0.52 | 1.07 | F | T |
| *Sumerlaeota* | *c__uncultured soil* | -0.47 | 1.01 | F | T |
| *Sumerlaeota* | *c__uncultured soil* | -0.47 | 1.01 | F | T |
| *Sumerlaeota* | *c__uncultured soil* | -0.47 | 1.01 | F | T |
| *Sumerlaeota* | *c__uncultured soil* | -0.47 | 1.01 | F | T |
| *Chloroflexota* | *g__OLB13* | -0.49 | 1.03 | F | T |
| *Armatimonadota* | *c__uncultured* | -0.47 | 1.00 | F | T |

| | | | | | |
|---|---|---|---|---|---|
| *Sumerlaeota* | *c__uncultured soil* | 0.04 | 0.48 | F | T |
| *Armatimonadota* | *c__uncultured* | 0.41 | 0.11 | T | T |
| *Sumerlaeota* | *c__uncultured* | -0.62 | 1.15 | F | T |
| *Armatimonadota* | *c__uncultured* | -0.36 | 0.88 | F | T |
| *Armatimonadota* | *c__uncultured* | -0.36 | 0.88 | T | F |
| *Armatimonadota* | *c__uncultured* | -0.36 | 0.88 | F | T |
| *Armatimonadota* | *c__uncultured* | -0.36 | 0.88 | T | F |
| *Rokubacteria* | *f__uncultured SPAM* | 0.34 | 0.18 | T | T |
| *Pseudomonadota* | *f__A21b* | -0.22 | 0.74 | F | T |
| *Patescibacteria* | *f__Saccharimonadaceae* | 1.15 | -0.64 | T | F |
| *Sumerlaeota* | *c__uncultured* | 1.92 | -1.41 | T | T |
| *Sumerlaeota* | *c__uncultured* | 1.92 | -1.41 | F | T |
| *Sumerlaeota* | *c__uncultured* | 1.92 | -1.41 | T | T |
| *Sumerlaeota* | *c__uncultured* | 1.92 | -1.41 | F | T |
| *Latescibacteria* | *c__uncultured proteo* | 0.46 | 0.05 | T | T |
| *Sumerlaeota* | *c__metagenome* | 0.94 | -0.45 | T | F |
| *Sumerlaeota* | *c__metagenome* | 0.94 | -0.45 | T | F |
| *Sumerlaeota* | *c__metagenome* | 0.94 | -0.45 | T | F |
| *Sumerlaeota* | *c__metagenome* | 0.94 | -0.45 | T | F |
| *Pseudomonadota* | *g__Geminicoccus* | -0.47 | 0.94 | F | T |
| *Sumerlaeota* | *c__uncultured* | 0.53 | -0.07 | T | T |
| *Sumerlaeota* | *c__uncultured soil* | 0.41 | 0.05 | F | T |
| *Sumerlaeota* | *c__uncultured soil* | 0.41 | 0.05 | T | T |
| *Sumerlaeota* | *c__uncultured soil* | 0.41 | 0.05 | F | T |
| *Sumerlaeota* | *c__uncultured soil* | 0.41 | 0.05 | T | T |
| *Sumerlaeota* | *c__uncultured* | 0.00 | 0.44 | F | T |
| *Sumerlaeota* | *c__uncultured* | 0.00 | 0.44 | T | T |
| *Sumerlaeota* | *c__uncultured* | 0.00 | 0.44 | F | T |
| *Sumerlaeota* | *c__uncultured* | 0.00 | 0.44 | T | T |
| *Euryarchaeota* | *g__Methanosarcina* | -0.01 | 0.44 | F | T |
| *Pseudomonadota* | *g__Syntrophus* | -0.58 | 1.00 | F | T |
| *Armatimonadota* | *c__uncultured* | -0.83 | 1.24 | F | T |
| *Latescibacteria* | *g__ uncultured* | 0.05 | 0.36 | F | T |
| *Bacillota* | *g__Streptococcus* | 0.45 | -0.04 | T | F |
| *Sumerlaeota* | *c__metagenome* | 0.40 | -0.03 | T | T |
| *Sumerlaeota* | *c__uncultured soil* | -0.05 | 0.40 | T | T |
| *Sumerlaeota* | *c__uncultured soil* | -0.05 | 0.40 | F | T |
| *Sumerlaeota* | *c__uncultured soil* | -0.05 | 0.40 | T | T |
| *Sumerlaeota* | *c__uncultured soil* | -0.05 | 0.40 | F | T |
| *Sumerlaeota* | *c__uncultured soil* | 0.86 | -0.52 | F | T |
| *Latescibacteria* | *c__uncultured prokaryote* | 1.10 | -0.76 | T | F |
| *Bacillota* | *g__Natronincola* | 0.29 | 0.05 | F | T |
| *Spirochaetota* | *g__Treponema 2* | 0.20 | 0.13 | T | T |

| | | | | | |
|---|---|---|---|---|---|
| *Patescibacteria* | *o__uncultured microorganism* | 0.41 | -0.08 | T | T |
| *Sumerlaeota* | *c__uncultured soil* | -0.03 | 0.37 | F | T |
| *Epsilonbacteraeota* | *g__Arcobacter* | 1.15 | -0.84 | T | F |
| *Bacteroidota* | *f__Flavobacteriaceae* | 0.23 | 0.08 | T | T |
| *Bacillota* | *g__Ruminococcaceae UCG-010* | 0.84 | -0.55 | T | F |
| *Pseudomonadota* | *f__Azospirillaceae* | -0.83 | 1.09 | F | T |
| *Bacteroidota* | *g__Chryseo* | 2.27 | -2.01 | F | T |
| *Pseudomonadota* | *f__27F-1492R* | -0.06 | 0.31 | F | T |
| *Pseudomonadota* | *g__Ottowia* | -0.10 | 0.35 | F | T |
| *Actinomycetota* | *f__Atopobiaceae* | 0.06 | 0.18 | F | T |
| *Spirochaetota* | *g__Treponema* | -0.71 | 0.93 | F | T |
| *Chloroflexota* | *o__uncultured Levilinea sp,* | -0.61 | 0.83 | F | T |
| *Chloroflexota* | *f__uncultured sludge A31* | 0.20 | 0.01 | T | T |
| *Bacteroidota* | *f__SB-5* | -0.46 | 0.65 | F | T |
| *Pseudomonadota* | *g__Bdellovibrio* | -0.27 | 0.44 | F | T |
| *Chloroflexota* | *f__Ktedonobacteraceae* | 0.34 | -0.17 | T | F |
| *Actinomycetota* | *g__CL500-29 marine group* | -0.01 | 0.16 | F | T |
| *Acidobacteriota* | *o__unidentified* | 0.03 | 0.10 | F | T |
| *Sumerlaeota* | *c__metagenome* | -0.08 | 0.21 | F | T |
| *Actinomycetota* | *c__MB-A2-108* | 0.05 | 0.07 | F | T |
| *Acidobacteriota* | *c__Subgroup 25* | -0.58 | 0.68 | F | T |
| *Latescibacteria* | *c__uncultured prokaryote* | 0.39 | -0.29 | T | F |
| *Pseudomonadota* | *o__CCD24* | -0.11 | 0.20 | F | T |
| *Cyanobacteriota* | *f__uncultured mle1-12* | -0.15 | 0.24 | F | T |
| *Actinomycetota* | *g__Agromyces* | 0.02 | 0.07 | F | T |
| *Sumerlaeota* | *c__uncultured soil* | 0.37 | -0.31 | F | T |
| *Sumerlaeota* | *c__uncultured soil* | 0.37 | -0.31 | F | T |
| *Sumerlaeota* | *c__uncultured soil* | 0.37 | -0.31 | F | T |
| *Sumerlaeota* | *c__uncultured soil* | 0.37 | -0.31 | F | T |
| *Acidobacteriota* | *g__Thermoanaerobaculum* | -0.15 | 0.20 | F | T |
| *Armatimonadota* | *c__uncultured* | 0.42 | -0.36 | F | T |
| *Armatimonadota* | *c__uncultured* | 0.42 | -0.36 | T | F |
| *Armatimonadota* | *c__uncultured* | 0.42 | -0.36 | F | T |
| *Armatimonadota* | *c__uncultured* | 0.42 | -0.36 | T | F |
| *Sumerlaeota* | *c__uncultured* | 0.74 | -0.68 | T | T |
| *Actinomycetota* | *g__Actinoallomurus* | -0.14 | 0.19 | F | T |
| *Chloroflexota* | *f__uncultured Chlorobi* | -0.11 | 0.15 | F | T |
| *Sumerlaeota* | *c__metagenome* | 0.38 | -0.35 | T | T |
| *Armatimonadota* | *c__uncultured* | 0.31 | -0.28 | T | F |
| *Pseudomonadota* | *g__Hirschia* | -0.06 | 0.08 | F | T |
| *Latescibacteria* | *c__uncultured prokaryote* | -0.49 | 0.50 | T | F |
| *Deinococcus-Thermus* | *g__Truepera* | -2.04 | 2.05 | T | F |
| *Latescibacteria* | *c__uncultured prokaryote* | -0.16 | 0.15 | F | T |

| | | | | | |
|---|---|---|---|---|---|
| *Chloroflexota* | *o__uncultured Chloroflexi* | -0.09 | 0.07 | F | T |
| *Actinomycetota* | *f__Nitriliruptoraceae* | -0.77 | 0.74 | T | F |
| *Pseudomonadota* | *o__Cellvibrionales* | -0.26 | 0.22 | F | T |
| *Sumerlaeota* | *c__metagenome* | -0.50 | 0.40 | F | T |
| *Sumerlaeota* | *c__metagenome* | -0.16 | 0.06 | F | T |
| *Pseudomonadota* | *g__Amaricoccus* | -0.14 | 0.01 | F | T |
| *Acidobacteriota* | *o__uncultured Holophaga sp,* | -0.16 | 0.03 | F | T |
| *Latescibacteria* | *c__uncultured prokaryote* | -0.30 | 0.17 | F | T |
| *Bacteroidota* | *f__Prolixibacteraceae* | -0.99 | 0.84 | F | T |
| *Latescibacteria* | *c__uncultured prokaryote* | -0.19 | 0.04 | F | T |
| *Actinomycetota* | *f__Streptosporangiaceae* | -0.17 | 0.00 | F | T |
| *Bacteroidota* | *f__Saprospiraceae* | -0.39 | 0.21 | F | T |
| *Pseudomonadota* | *g__Microvirga* | -0.24 | 0.04 | F | T |
| *Chloroflexota* | *f__uncultured Caldilineaceae* | 0.13 | -0.34 | T | F |
| *Pseudomonadota* | *f__uncultured delta proteo* | -0.28 | 0.04 | F | T |
| *Sumerlaeota* | *c__Omnitrophica OLB16* | -0.29 | 0.02 | F | T |
| *Acidobacteriota* | *o__uncultured Acidobacteriales* | 0.32 | -0.60 | F | T |
| *Acidobacteriota* | *o__uncultured Acidobacteriales* | 0.32 | -0.60 | T | F |
| *Acidobacteriota* | *o__uncultured Acidobacteriales* | 0.32 | -0.60 | F | T |
| *Acidobacteriota* | *o__uncultured Acidobacteriales* | 0.32 | -0.60 | T | F |
| *Armatimonadota* | *c__uncultured* | 2.07 | -2.36 | F | T |
| *Sumerlaeota* | *c__uncultured* | -0.12 | -0.20 | F | T |
| *Sumerlaeota* | *c__metagenome* | 1.14 | -1.47 | F | T |
| *Armatimonadota* | *c__uncultured* | -0.31 | -0.03 | F | T |
| *Bacillota* | *g__Alkalibacter* | -0.60 | 0.26 | F | T |
| *Sumerlaeota* | *c__metagenome* | 0.47 | -0.82 | F | T |
| *Pseudomonadota* | *g__Amphiplicatus* | -0.58 | 0.23 | F | T |
| *Sumerlaeota* | *c__uncultured soil* | -0.03 | -0.34 | F | T |
| *Pseudomonadota* | *f__Halieaceae* | -0.64 | 0.27 | F | T |
| *Sumerlaeota* | *c__metagenome* | -0.05 | -0.34 | F | T |
| *Latescibacteria* | *c__uncultured prokaryote* | -0.29 | -0.10 | T | F |
| *Acidobacteriota* | *o__uncultured 259* | 0.50 | -0.90 | F | T |
| *Sumerlaeota* | *c__uncultured* | -0.36 | -0.05 | F | T |
| *Chloroflexota* | *f__uncultured Caldilineaceae* | 0.20 | -0.63 | T | F |
| *WS4* | *c__uncultured organism* | -0.09 | -0.34 | F | T |
| *Pseudomonadota* | *g__Desulfovibrio* | 1.20 | -1.67 | F | T |
| *Pseudomonadota* | *g__mle1-7* | -0.28 | -0.19 | F | T |
| *Bacillota* | *c__Clostridia* | -0.58 | 0.10 | F | T |
| *Pseudomonadota* | *g__Syntrophorhabdus* | -0.37 | -0.12 | T | F |
| *Latescibacteria* | *c__uncultured proteo* | -0.26 | -0.24 | T | T |
| *Planctomycetota* | *g__Urania-1B-19 marine sediment* | -1.67 | 1.18 | T | F |
| *Chloroflexota* | *o__uncultured Chloroflexi* | 0.96 | -1.46 | T | T |
| *Chloroflexota* | *o__uncultured Chloroflexi* | 0.96 | -1.46 | T | T |

| | | | | | |
|---|---|---|---|---|---|
| *Chloroflexota* | *o__uncultured Chloroflexi* | 0.96 | -1.46 | T | T |
| *Chloroflexota* | *o__uncultured Chloroflexi* | 0.96 | -1.46 | T | T |
| *Pseudomonadota* | *g__Sulfurifustis* | -0.78 | 0.28 | F | T |
| *Latescibacteria* | *c__uncultured prokaryote* | -0.22 | -0.29 | T | F |
| *Pseudomonadota* | *g__marine metagenome* | -0.68 | 0.16 | F | T |
| *Latescibacteria* | *c__uncultured prokaryote* | -0.14 | -0.38 | F | T |
| *WS4* | *c__uncultured organism* | -0.77 | 0.25 | F | T |
| *Acidobacteriota* | *o__Acidobacteria SCN 69-37* | -0.83 | 0.24 | F | T |
| *Chloroflexota* | *c__JG30-KF-CM66* | -0.86 | 0.27 | F | T |
| *Pseudomonadota* | *o__CCM19a* | -0.76 | 0.17 | F | T |
| *Acidobacteriota* | *o__uncultured deep-sea* | -0.93 | 0.33 | F | T |
| *Sumerlaeota* | *c__metagenome* | 1.29 | -1.91 | F | T |
| *Sumerlaeota* | *c__metagenome* | 0.33 | -0.94 | T | F |
| *Pseudomonadota* | *g__Lautropia* | -0.75 | 0.13 | F | T |
| *Acidobacteriota* | *o__uncultured Acidobacteriales* | -0.76 | 0.13 | F | T |
| *Acidobacteriota* | *o__uncultured Acidobacteriales* | -0.76 | 0.13 | T | F |
| *Acidobacteriota* | *o__uncultured Acidobacteriales* | -0.76 | 0.13 | F | T |
| *Acidobacteriota* | *o__uncultured Acidobacteriales* | -0.76 | 0.13 | T | F |
| *Pseudomonadota* | *f__uncultured Anaeromyxobacter sp,* | -0.69 | 0.03 | F | T |
| *Latescibacteria* | *c__uncultured proteo* | 1.03 | -1.71 | T | T |
| *Caldiserica* | *g__Caldisericum* | 0.06 | -0.74 | T | F |
| *Armatimonadota* | *c__uncultured* | 1.15 | -1.83 | F | T |
| *Sumerlaeota* | *c__uncultured soil* | -0.58 | -0.12 | F | T |
| *Sumerlaeota* | *c__uncultured soil* | -0.58 | -0.12 | F | T |
| *Sumerlaeota* | *c__uncultured soil* | -0.58 | -0.12 | F | T |
| *Sumerlaeota* | *c__uncultured soil* | -0.58 | -0.12 | F | T |
| *Bacillota* | *g__Tepidibacillus* | 1.14 | -1.87 | T | F |
| *Pseudomonadota* | *f__Paracaedibacteraceae* | 0.77 | -1.51 | T | F |
| *Pseudomonadota* | *g__Hydrogenophaga* | -1.00 | 0.24 | F | T |
| *Sumerlaeota* | *c__uncultured soil* | -0.45 | -0.35 | T | T |
| *Pseudomonadota* | *g__Escherichia-Shigella* | 1.12 | -1.94 | F | T |
| *Pseudomonadota* | *g__Ellin6067* | -0.51 | -0.33 | T | F |
| *Bacteroidota* | *g__Bact-08* | 1.97 | -2.81 | F | T |
| *Sumerlaeota* | *c__uncultured soil* | 0.46 | -1.30 | F | T |
| *Sumerlaeota* | *c__uncultured soil* | 0.46 | -1.30 | F | T |
| *Sumerlaeota* | *c__uncultured soil* | 0.46 | -1.30 | F | T |
| *Sumerlaeota* | *c__uncultured soil* | 0.46 | -1.30 | F | T |
| *Gemmatimonadetes* | *g__Gemmatimonas* | -0.40 | -0.45 | T | F |
| *Sumerlaeota* | *c__uncultured soil PBS-II-34* | -0.58 | -0.28 | T | F |
| *Spirochaetota* | *g__Spirochaeta 2* | -0.58 | -0.29 | T | F |
| *Sumerlaeota* | *c__uncultured* | -0.07 | -0.80 | T | T |
| *Patescibacteria* | *f__uncultured Sphingobium sp,* | 0.90 | -1.78 | T | F |
| *Armatimonadota* | *c__uncultured* | -0.09 | -0.82 | T | F |

| | | | | | |
|---|---|---|---|---|---|
| *Bacillota* | *f__Erysipelotrichaceae* | 0.50 | -1.42 | F | T |
| *Armatimonadota* | *c__uncultured* | 0.47 | -1.41 | T | F |
| *Bacillota* | *g__Lysinibacillus* | 1.67 | -2.61 | F | T |
| *Pseudomonadota* | *g__Chthonobacter* | -1.02 | 0.08 | F | T |
| *Sumerlaeota* | *c__uncultured* | 0.29 | -1.24 | F | T |
| *Armatimonadota* | *c__uncultured* | -0.63 | -0.36 | T | F |
| *Bacillota* | *f__Christensenellaceae* | 0.66 | -1.66 | F | T |
| *Armatimonadota* | *c__uncultured* | -0.58 | -0.42 | T | F |
| *Bacillota* | *g__Dehalobacter* | 0.59 | -1.60 | F | T |
| *Sumerlaeota* | *c__metagenome* | -0.21 | -0.82 | T | T |
| *Bacteroidota* | *f__Cyclobacteriaceae* | 0.71 | -1.78 | T | F |
| *Sumerlaeota* | *c__metagenome* | -0.71 | -0.38 | T | F |
| *Sumerlaeota* | *c__metagenome* | -0.71 | -0.38 | T | F |
| *Sumerlaeota* | *c__metagenome* | -0.71 | -0.38 | T | F |
| *Sumerlaeota* | *c__metagenome* | -0.71 | -0.38 | T | F |
| *Acidobacteriota* | *f__uncultured Acidobacteria* | -0.36 | -0.76 | T | T |
| *Actinomycetota* | | 1.04 | -2.18 | F | T |
| *Bacteroidota* | *g__Wandonia* | 0.98 | -2.16 | F | T |
| *Pseudomonadota* | *g__Archangium* | -0.96 | -0.22 | F | T |
| *Chloroflexota* | *g__Leptolinea* | -1.16 | -0.03 | T | F |
| *Latescibacteria* | *c__uncultured prokaryote* | -0.72 | -0.47 | T | F |
| *WS2* | *c__uncultured Firmicutes* | -0.40 | -0.80 | F | T |
| *Sumerlaeota* | *c__uncultured* | -1.42 | 0.21 | F | T |
| *Chloroflexota* | *o__SBR1031* | -0.82 | -0.40 | T | F |
| *Actinomycetota* | *g__Actinomycetospora* | -0.83 | -0.41 | T | F |
| *Pseudomonadota* | *g__Marinibaculum* | -0.70 | -0.57 | F | T |
| *Pseudomonadota* | *g__Roseomonas* | -0.72 | -0.55 | T | F |
| *Bacteroidota* | *g__Petrimonas* | 0.84 | -2.12 | F | T |
| *Bacteroidota* | *f__Tannerellaceae* | 1.38 | -2.67 | F | T |
| *Bacillota* | *g__Acetanaero* | 0.92 | -2.30 | F | T |
| *Verrucomicrobiota* | *g__Cerasicoccus* | -1.42 | -0.01 | F | T |
| *Actinomycetota* | *f__uncultured Coriobacteriaceae* | 0.42 | -1.89 | F | T |
| *Bacillota* | *g__Sporo* | 0.12 | -1.61 | F | T |
| *Deinococcus-Thermus* | *g__Thermus* | -0.99 | -0.51 | T | T |
| *Bacillota* | *g__Cohnella* | -0.81 | -0.74 | T | F |
| *Pseudomonadota* | *g__Pajaroellobacter* | -0.93 | -0.68 | T | F |
| *Acidobacteriota* | *g__Bryobacter* | -0.95 | -0.71 | T | F |
| *Bacteroidota* | *g__Cloaci* | 1.28 | -2.97 | F | T |
| *Chloroflexota* | *o__Thermomicrobiales* | -0.85 | -0.85 | T | F |
| *Pseudomonadota* | *g__uncultured alpha proteo* | -0.63 | -1.08 | T | T |
| *Armatimonadota* | *c__uncultured* | 0.44 | -2.15 | F | T |
| *Armatimonadota* | *c__uncultured* | -1.01 | -0.73 | T | F |
| *Bacillota* | *f__Veillonellaceae* | 0.39 | -2.23 | F | T |

| | | | | | |
|---|---|---|---|---|---|
| *Bacillota* | *g__Ruminiclostridium 9* | 0.45 | -2.32 | F | T |
| *Sumerlaeota* | *c__uncultured soil* | -0.70 | -1.25 | T | T |
| *Sumerlaeota* | *c__uncultured soil* | -0.70 | -1.25 | F | T |
| *Sumerlaeota* | *c__uncultured soil* | -0.70 | -1.25 | T | T |
| *Sumerlaeota* | *c__uncultured soil* | -0.70 | -1.25 | F | T |
| *Latescibacteria* | *c__uncultured prokaryote* | -0.38 | -1.57 | F | T |
| *Cyanobacteriota* | *f__Brachypodium distachyon* | -1.65 | -0.32 | T | F |
| *Bacteroidota* | *f__Hymenobacteraceae* | 0.06 | -2.03 | F | T |
| *Armatimonadota* | *c__uncultured* | 0.85 | -2.85 | F | T |
| *Tenericutes* | *o__Mollicutes RF39* | 0.74 | -2.74 | F | T |
| *Actinomycetota* | *o__Microtrichales* | -0.08 | -1.95 | F | T |
| *Bacillota* | *g__Erysipelotrichaceae UCG-004* | 0.66 | -2.70 | F | T |
| *Sumerlaeota* | *c__uncultured* | -1.34 | -0.74 | T | T |
| *Actinomycetota* | *o__Corynebacteriales* | -0.76 | -1.33 | T | T |
| *Actinomycetota* | *f__Micromonosporaceae* | -0.98 | -1.12 | T | F |
| *Patescibacteria* | *o__Saccharimonadales* | 0.18 | -2.29 | F | T |
| *Pseudomonadota* | *f__Steroidobacteraceae* | -1.17 | -0.95 | T | T |
| *Gemmatimonadetes* | *c__BD2-11 terrestrial group* | -1.76 | -0.37 | F | T |
| *Bacillota* | *g__Desulfito* | 1.02 | -3.21 | F | T |
| *Acidobacteriota* | *o__uncultured 98* | -1.64 | -0.58 | T | F |
| *Armatimonadota* | *c__uncultured* | 0.81 | -3.06 | F | T |
| *Pseudomonadota* | *g__Myxococcus* | -1.26 | -1.03 | T | T |
| *Chloroflexota* | *g__Paraburkholderia tropica* | -1.39 | -0.92 | T | T |
| *Pseudomonadota* | *f__Archangiaceae* | -0.98 | -1.35 | T | T |
| *Epsilonbacteraeota* | *o__Campylobacterales* | -1.29 | -1.04 | F | T |
| *Pseudomonadota* | *g__Rickettsiella* | -1.14 | -1.24 | T | T |
| *Bacillota* | *g__Fonticella* | 1.00 | -3.40 | F | T |
| *Dependentiae* | *f__Vermiphilaceae* | -1.07 | -1.36 | T | T |
| *Bacillota* | *g__Dehalo* | -1.46 | -0.97 | T | F |
| *Gemmatimonadetes* | *o__uncultured actino* | -1.36 | -1.08 | F | T |
| *Sumerlaeota* | *c__uncultured soil* | -0.78 | -1.70 | F | T |
| *Sumerlaeota* | *c__uncultured soil* | -0.78 | -1.70 | F | T |
| *Sumerlaeota* | *c__uncultured soil* | -0.78 | -1.70 | F | T |
| *Sumerlaeota* | *c__uncultured soil* | -0.78 | -1.70 | F | T |
| *Pseudomonadota* | *g__Desulfatiglans* | -1.24 | -1.26 | T | T |
| *Bacillota* | *g__Desulfurispora* | 0.48 | -3.02 | F | T |
| *Bacteroidota* | *f__ enrichment culture clone M624* | 0.75 | -3.29 | F | T |
| *Armatimonadota* | *c__uncultured* | 0.56 | -3.13 | F | T |
| *Sumerlaeota* | *c__uncultured soil* | -0.90 | -1.73 | T | T |
| *Bacillota* | *g__Ruminococcaceae UCG-013* | -0.49 | -2.16 | F | T |
| *Bacillota* | *g__Dolosigranulum* | -1.46 | -1.19 | T | T |
| *Bacillota* | *g__Anaerosporomusa* | -0.60 | -2.05 | T | T |
| *Bacillota* | *g__Oscillibacter* | -0.22 | -2.52 | F | T |

| | | | | | |
|---|---|---|---|---|---|
| *Actinomycetota* | *g__Raoultibacter* | -1.16 | -1.59 | T | T |
| *Pseudomonadota* | *g__Massilia* | -1.40 | -1.36 | T | T |
| *Pseudomonadota* | *g__Desulfuromonas* | -1.74 | -1.03 | T | T |
| *Chloroflexota* | *g__Candidatus Chloroploca* | -1.56 | -1.28 | T | T |
| *Armatimonadota* | *o__uncultured SJA-121* | 0.57 | -3.42 | F | T |
| *Actinomycetota* | *g__Propionicicella* | 1.23 | -4.12 | F | T |
| *Actinomycetota* | | 0.72 | -3.61 | F | T |
| *Bacillota* | *g__Desulfosporosinus* | 1.37 | -4.28 | F | T |
| *Bacillota* | *f__Ruminococcaceae* | 0.83 | -3.75 | F | T |
| *Nitrospirae* | *f__uncultured Nitrospirae* | -1.41 | -1.51 | T | T |
| *Sumerlaeota* | *c__uncultured* | -1.71 | -1.28 | T | T |
| *Bacillota* | *f__Peptococcaceae* | -0.25 | -2.79 | F | T |
| *Bacillota* | *g__Paenibacillus* | -0.15 | -2.92 | F | T |
| *Pseudomonadota* | *f__unidentified wb1_N15* | -0.96 | -2.15 | T | T |
| *Actinomycetota* | *g__Rhodococcus* | -1.30 | -1.81 | T | T |
| *Cyanobacteriota* | *o__Nostocales* | -1.13 | -2.00 | T | T |
| *Bacillota* | *g__Jeotgalicoccus* | -1.43 | -1.76 | T | T |
| *Bacillota* | *g__Anaerosporobacter* | -1.07 | -2.15 | T | T |
| *Sumerlaeota* | *c__uncultured* | -1.82 | -1.54 | F | T |
| *Bacillota* | *f__Planococcaceae* | -1.80 | -1.58 | T | F |
| *Pseudomonadota* | *g__Allorhizobium* | 1.67 | -5.06 | F | T |
| *Patescibacteria* | *o__JGI 0000069-P22* | -2.57 | -0.88 | T | F |
| *Bacteroidota* | *f__Prevotellaceae* | 0.87 | -4.35 | F | T |
| *Actinomycetota* | *g__Brevi* | -1.66 | -1.95 | T | T |
| *Bacillota* | *g__Anaero* | -1.27 | -2.38 | F | T |
| *Actinomycetota* | *f__Eggerthellaceae* | -1.56 | -2.11 | T | T |
| *Actinomycetota* | *f__Propionibacteriaceae* | -0.64 | -3.19 | F | T |
| *Bacillota* | *g__Alkalicoccus* | -2.10 | -1.82 | T | T |
| *Pseudomonadota* | *g__Azospirillum* | 1.97 | -5.90 | F | T |
| *Cyanobacteriota* | *f__Coelastrella sp, M60* | -2.13 | -1.85 | T | T |
| *Spirochaetota* | *f__Spirochaetaceae* | -0.54 | -3.59 | F | T |
| *Bacillota* | *g__Caproiciproducens* | -1.29 | -2.87 | T | T |
| *Bacteroidota* | *f__Weeksellaceae* | 0.72 | -4.95 | F | T |
| *Actinomycetota* | *g__Dietzia* | -0.99 | -3.25 | F | T |
| *Chloroflexota* | *f__A4b* | -2.12 | -2.22 | T | F |
| *Actinomycetota* | *g__Cuti* | -1.25 | -3.12 | F | T |
| *Pseudomonadota* | *f__Xanthobacteraceae* | -0.08 | -4.30 | F | T |
| *Planctomycetota* | *g__Planctomicrobium* | -2.39 | -2.11 | T | T |
| *Pseudomonadota* | *g__Aureimonas* | -1.63 | -2.93 | T | T |
| *Actinomycetota* | *f__Corynebacteriaceae* | -2.04 | -2.56 | T | T |
| *WS4* | *c__uncultured organism* | -2.51 | -2.23 | T | F |
| *Bacillota* | *g__Oxobacter* | -0.24 | -4.51 | F | T |
| *Bacillota* | *o__Clostridiales* | -0.35 | -4.43 | F | T |

| | | | | | |
|---|---|---|---|---|---|
| *Pseudomonadota* | *g__Nitrincola* | -2.07 | -2.73 | T | T |
| *Pseudomonadota* | *g__GKS98 freshwater group* | -0.25 | -4.56 | F | T |
| *Cyanobacteriota* | *f__gut metagenome* | -2.56 | -2.29 | T | T |
| *Pseudomonadota* | *o__Rhodospirillales* | -2.60 | -2.33 | T | T |
| *Acidobacteriota* | *g__Holophaga* | -2.60 | -2.33 | T | T |
| *Actinomycetota* | *g__Brachy* | -2.16 | -2.81 | T | T |
| *Planctomycetota* | *g__Pir3 lineage* | -2.63 | -2.36 | T | T |
| *Bacillota* | *g__uncultured compost* | -2.63 | -2.36 | T | T |
| *Bacillota* | *g__Sinobaca* | -2.64 | -2.36 | T | T |
| *Planctomycetota* | *g__Blastopirellula* | -2.69 | -2.42 | T | T |
| *Latescibacteria* | *c__uncultured proteo* | -2.52 | -2.60 | T | T |
| *Actinomycetota* | *g__Gordonia* | -2.79 | -2.50 | T | T |
| *Pseudomonadota* | *g__Alishewanella* | -0.99 | -4.34 | F | T |
| *Bacillota* | *o__Bacillales* | -1.67 | -3.66 | F | T |
| *Actinomycetota* | *g__Kocuria* | -2.82 | -2.55 | T | T |
| *Actinomycetota* | *g__Micro* | -1.85 | -3.59 | F | T |
| *Acidobacteriota* | *g__Geothrix* | -0.77 | -4.69 | F | T |
| *Bacillota* | *g__Enterococcus* | -2.21 | -3.28 | T | T |
| *Pseudomonadota* | *g__Methylophaga* | -2.52 | -3.00 | T | T |
| *Pseudomonadota* | *g__Gemmobacter* | -2.63 | -3.02 | T | T |
| *Actinomycetota* | *g__Jonesia* | -2.61 | -3.10 | T | T |
| *Cyanobacteriota* | *o__Chloroplast* | -1.95 | -3.76 | F | T |
| *Bacillota* | *f__Limnochordaceae* | -3.04 | -2.76 | T | T |
| *Cyanobacteriota* | *o__Vampirovibrionales* | -3.12 | -2.85 | T | T |
| *Actinomycetota* | *g__Coryne 1* | -3.28 | -3.01 | T | T |
| *Bacteroidota* | *o__Sphingobacteriales* | -3.89 | -2.50 | T | T |
| *Bacteroidota* | *g__Polaribacter* | -2.83 | -3.75 | T | T |
| *Pseudomonadota* | *g__Comamonas* | -3.61 | -3.00 | T | F |
| *Dependentiae* | *g__candidate division TM6* | -2.87 | -3.82 | T | T |
| *Actinomycetota* | *g__Nesterenkonia* | -3.27 | -3.69 | T | T |
| *WS4* | *c__uncultured organism* | -3.90 | -3.63 | T | T |
| *Bacillota* | *g__Salinicoccus* | -4.09 | -3.99 | T | T |
| *Chlamydiota* | *o__Chlamydiales* | -3.98 | -7.67 | F | T |
| *Pseudomonadota* | *g__Roseibaca* | -6.50 | -5.77 | T | T |

**Table S3**: Unique MAGs detected in suspension samples across the three conditions ('before' B, 'initial' I, and 'later' L). MAGs were reconstructed, dereplicated, taxonomically classified, and quality-checked using established binning and QC pipelines.

| Aeration | MAG | Phylum | Family | Genus | Completeness [%] | Contamination [%] | Genome size [Mbp] | GC content [%] |
|---|---|---|---|---|---|---|---|---|
| B | 1 | *Pseudomonadota* | *Beijerinckiaceae* | *Saliniramus* | 97.2 | 3.0 | 3.8 | 60 |
| | 2 | *Pseudomonadota* | *Rhodobacteraceae* | *Paracoccus* | 64.8 | 0.4 | 2.8 | 60 |
| | 3 | *Pseudomonadota* | *Halomonadaceae* | *Vreelandella* | 99.0 | 2.4 | 4.7 | 50 |
| | 4 | *Actinomycetota* | *Microbacteriaceae* | *Microbacterium* | 100.0 | 0.1 | 3.3 | 70 |
| | 5 | *Pseudomonadota* | *Alteromonadaceae* | *Aliidiomarina* | 83.7 | 0.6 | 2.3 | 90 |
| | 6 | *Pseudomonadota* | *Midichloriaceae* | | 95.8 | 0.0 | 1.0 | 30 |
| | 7 | *Pseudomonadota* | *Rhodobacteraceae* | *EhC02* | 100.0 | 0.5 | 4.1 | 60 |
| | 8 | *Pseudomonadota* | *Sphingomonadaceae* | *Erythrobacter* | 60.9 | 1.2 | 1.8 | 60 |
| | 9 | *Pseudomonadota* | *Burkholderiaceae* | *Verticiella* | 95.4 | 0.5 | 3.4 | 60 |
| | 10 | *Pseudomonadota* | *Devosiaceae* | *Pelagibacterium* | 93.0 | 0.0 | 3.6 | 50 |
| | 11 | *Chlamydiota* | | | 96.3 | 0.5 | 1.8 | 50 |
| | 12 | *Planctomycetota* | *Planctomycetaceae* | *Rubinisphaera* | 61.3 | 0.4 | 3.3 | 50 |
| | 13 | *Bacteroidota* | *Crocinitomicaceae* | *JALQTG01* | 62.6 | 0.5 | 2.3 | 30 |
| | 14 | *Actinomycetota* | *Nitriliruptoraceae* | | 96.5 | 0.3 | 3.0 | 60 |
| | 15 | *Actinomycetota* | *Microbacteriaceae* | *Microcella* | 96.2 | 3.2 | 2.5 | 60 |
| | 16 | *Pseudomonadota* | *Moraxellaceae* | *Acinetobacter* | 72.2 | 2.4 | 2.3 | 40 |
| | 17 | *Pseudomonadota* | *Alteromonadaceae* | *Aliidiomarina* | 92.4 | 1.0 | 3.0 | 50 |
| | 18 | *Pseudomonadota* | *Methylophagaceae* | *Methylophaga* | 68.5 | 0.8 | 1.7 | 50 |
| | 19 | *Pseudomonadota* | *Vibrionaceae* | *Vibrio* | 88.9 | 2.4 | 3.1 | 40 |
| I | 1 | *Bacteroidota* | *Cyclobacteriaceae* | *Lunatimonas* | 100.0 | 2.1 | 7.2 | 40 |
| | 2 | *Pseudomonadota* | *PWYM01* | *PWYM01* | 56.4 | 3.1 | 1.8 | 60 |
| | 3 | *Pseudomonadota* | | | 96.2 | 0.1 | 1.8 | 40 |
| | 4 | *Pseudomonadota* | *Acetobacteraceae* | *Roseococcus* | 68.1 | 0.5 | 3.1 | 70 |
| | 5 | *Verrucomicrobiota* | *Akkermansiaceae* | *Luteolibacter* | 96.1 | 0.6 | 4.0 | 60 |

| | | | | | | | | |
|---|---|---|---|---|---|---|---|---|
| | 6 | *Pseudomonadota* | *Legionellaceae* | *Legionella_D* | 100.0 | 0.2 | 2.7 | 40 |
| | 7 | *Pseudomonadota* | *Natronospirillaceae* | *Saccharospirillum* | 99.2 | 2.3 | 4.7 | 50 |
| | 8 | *Bacteroidota* | *Cyclonatronaceae* | *Cyclonatronum* | 100.0 | 0.0 | 4.1 | 50 |
| | 1 | *Pseudomonadota* | *Geminicoccaceae* | *SLRI01* | 99.7 | 0.1 | 5.7 | 60 |
| | 2 | *Pseudomonadota* | *Rhodobacteraceae* | *JAFIEN01* | 98.4 | 1.7 | 4.0 | 60 |
| | 3 | *Verrucomicrobiota* | *DYNR01* | | 98.6 | 0.1 | 3.3 | 50 |
| | 4 | *Pseudomonadota* | *Rhodocyclaceae* | *Azoarcus_F* | 78.5 | 0.4 | 2.9 | 60 |
| | 5 | *Bacteroidota* | *Crocinitomicaceae* | | 99.9 | 0.4 | 3.4 | 30 |
| L | 6 | *Myxococcota* | *PHBI01* | *SYAK01* | 92.0 | 3.1 | 5.8 | 70 |
| | 7 | *Pseudomonadota* | | | 79.1 | 0.4 | 3.4 | 70 |
| | 8 | *Pseudomonadota* | *Rickettsiaceae* | *Megaera* | 96.5 | 0.1 | 1.4 | 30 |
| | 9 | *Pseudomonadota* | *UBA12402* | *1-14-0-20-46-22* | 77.2 | 0.3 | 1.4 | 40 |
| | 10 | *Planctomycetota* | *Phycisphaeraceae* | | 93.0 | 1.8 | 3.0 | 60 |
| | 11 | *Bacteroidota* | *HLUCCA01* | *HLUCCA01* | 94.4 | 0.1 | 3.5 | 50 |

**Table S4:** List of genes associated with carbon fixation (ko00710, ko00720) and nitrogen metabolism (ko00910) according to KEGG pathways. Gene names were assigned based on their corresponding KEGG KO identifiers.

| Gene name | KEGG_ko | Metabolism |
|---|---|---|
| *mdh* | K00024 | Carbon |
| *MDH1* | K00025 | |
| *MDH2* | K00026 | |
| *E1.1.1.39* | K00028 | |
| *maeB* | K00029 | |
| *IDH1, IDH2, icd* | K00031 | |
| *E1.1.1.82* | K00051 | |
| *GAPDH, gapA* | K00134 | |
| *gap2, gapB* | K00150 | |
| *porA* | K00169 | |
| *porB* | K00170 | |
| *porD* | K00171 | |
| *porC, porG* | K00172 | |
| *korA, oorA, oforA* | K00174 | |
| *korB, oorB, oforB* | K00175 | |
| *korD, oorD* | K00176 | |
| *korC, oorC* | K00177 | |
| *vorG, porG* | K00189 | |
| *cdhD, acsD* | K00194 | |
| *cooF* | K00196 | |
| *cdhE, acsC* | K00197 | |
| *cooS, acsA* | K00198 | |
| *frdA, fdrA* | K00233 | |
| *sdhA, frdA* | K00239 | |
| *sdhB, frdB* | K00240 | |
| *sdhC, frdC* | K00241 | |
| *sdhD, frdD* | K00242 | |
| *frdA* | K00244 | |
| *frdB* | K00245 | |
| *frdC* | K00246 | |
| *frdD* | K00247 | |
| *E2.2.1.1, tktA, tktB* | K00615 | |
| *TALDO1, talB, talA* | K00616 | |
| *GPT, ALT* | K00814 | |
| *PRK* | K00855 | |
| *PGK* | K00927 | |
| *ppdK* | K01006 | |
| *fbp-SEBP* | K01086 | |

| | |
|---|---|
| *E3.1.3.37* | K01100 |
| *rbcL, cbbL* | K01601 |
| *rbcS, cbbS* | K01602 |
| *pckA* | K01610 |
| *ALDO* | K01623 |
| *FBA, fbaA* | K01624 |
| *rpe* | K01783 |
| *TPI, tpiA* | K01803 |
| *rpiA* | K01807 |
| *rpiB* | K01808 |
| *glpX* | K02446 |
| *FBP* | K03841 |
| *fbp3* | K04041 |
| *GAPA* | K05298 |
| *glpX-SEBP* | K11532 |
| *fbaB* | K11645 |
| *tal-pgi* | K13810 |
| *GGAT* | K14272 |
| *GOT1* | K14454 |
| *metF, MTHFR* | K00297 |
| *pta* | K00625 |
| *ACAT, atoB* | K00626 |
| *ackA* | K00925 |
| *pps, ppsA* | K01007 |
| *folD* | K01491 |
| *fchA* | K01500 |
| *ppc* | K01595 |
| *ACLY* | K01648 |
| *fumD* | K01675 |
| *E4.2.1.2A, fumA, fumB* | K01676 |
| *E4.2.1.2AA, fumA* | K01677 |
| *E4.2.1.2AB, fumB* | K01678 |
| *E4.2.1.2B, fumC, FH* | K01679 |
| *ACO* | K01681 |
| *acnB* | K01682 |
| *fumE* | K01774 |
| *MUT* | K01847 |
| *E5.4.99.2A, mcmA1* | K01848 |
| *E5.4.99.2B, mcmA2* | K01849 |
| *ACSS1_2, acs* | K01895 |
| *sucD* | K01902 |

| | |
|---|---|
| *sucC* | K01903 |
| *fhs* | K01938 |
| *PC, pyc* | K01958 |
| *pycA* | K01959 |
| *pycB* | K01960 |
| *accC* | K01961 |
| *accA* | K01962 |
| *accD* | K01963 |
| *K01964* | K01964 |
| *accB, bccP* | K02160 |
| *por, nifJ* | K03737 |
| *fdhA* | K05299 |
| *MCEE, epi* | K05606 |
| *mcl* | K08691 |
| *meh* | K09709 |
| *pta* | K13788 |
| *acsB* | K14138 |
| *mch, mcd* | K14449 |
| *GOT2* | K14455 |
| *K14465* | K14465 |
| *K14466* | K14466 |
| *4hbl* | K14467 |
| *mcr* | K14468 |
| *K14469* | K14469 |
| *mct* | K14470 |
| *smtA1* | K14471 |
| *smtB* | K14472 |
| *abfD* | K14534 |
| *K15016* | K15016 |
| *K15017* | K15017 |
| *K15018* | K15018 |
| *K15019* | K15019 |
| *K15020* | K15020 |
| *fdhB* | K15022 |
| *acsE* | K15023 |
| *K15024* | K15024 |
| *K15036* | K15036 |
| *K15037* | K15037 |
| *K15038* | K15038 |
| *K15039* | K15039 |
| *K15052* | K15052 |

| | | |
|---|---|---|
| *aclA* | K15230 | |
| *aclB* | K15231 | |
| *ccsA* | K15232 | |
| *ccsB* | K15233 | |
| *ccl* | K15234 | |
| *tfrA* | K18209 | |
| *tfrB* | K18210 | |
| *frdA* | K18556 | |
| *frdB* | K18557 | |
| *frdC* | K18558 | |
| *frdD* | K18559 | |
| *frdE* | K18560 | |
| *E6.2.1.56* | K18593 | |
| *K18594* | K18594 | |
| *K18602* | K18602 | |
| *K18603* | K18603 | |
| *K18604* | K18604 | |
| *K18605* | K18605 | |
| *sdhD, frdD* | K18859 | |
| *sdhD, frdD* | K18860 | |
| *K18861* | K18861 | |
| *fdhF* | K22015 | |
| *metV* | K25007 | |
| *rnfC2* | K25008 | |
| *hydA2* | K25123 | |
| *hycB* | K25124 | |
| *mtgA* | K25221 | |
| *K25774* | K25774 | |
| *mtvA* | K25915 | |
| *frdB, fdrB* | K25995 | |
| *frdC, fdrC* | K25996 | |
| *fccA, fcc3* | K26318 | |
| *mtqA* | K26930 | |
| *acnA* | K27802 | |
| *gudB, rocG* | K00260 | Nitrogen |
| *GLUD1_2, gdhA* | K00261 | |
| *E1.4.1.4, gdhA* | K00262 | |
| *GLT1* | K00264 | |
| *gltB* | K00265 | |
| *gltD* | K00266 | |
| *GLU, gltS* | K00284 | |

| | |
|---|---|
| *nasB* | K00360 |
| *nasB* | K00361 |
| *nirB* | K00362 |
| *nirD* | K00363 |
| *nirA* | K00366 |
| *narB* | K00367 |
| *nirK* | K00368 |
| *narG, narZ, nxrA* | K00370 |
| *narH, narY, nxrB* | K00371 |
| *nasC, nasA* | K00372 |
| *narI, narV* | K00374 |
| *nosZ* | K00376 |
| *ncd2, npd* | K00459 |
| *anfG* | K00531 |
| *arcC* | K00926 |
| *E3.5.1.49* | K01455 |
| *E3.5.5.1* | K01501 |
| *CA* | K01672 |
| *cynT, can* | K01673 |
| *cah* | K01674 |
| *cynS* | K01725 |
| *E4.2.1.1* | K01743 |
| *glnA, GLUL* | K01915 |
| *CPS1* | K01948 |
| *norC* | K02305 |
| *napA* | K02567 |
| *napB* | K02568 |
| *NRT2, narK, nrtP, nasA* | K02575 |
| *nifD* | K02586 |
| *nifH* | K02588 |
| *nifK* | K02591 |
| *nrfA* | K03385 |
| *norB* | K04561 |
| *hcp* | K05601 |
| *NR* | K10534 |
| *hao* | K10535 |
| *pmoA-amoA* | K10944 |
| *pmoB-amoB* | K10945 |
| *pmoC-amoC* | K10946 |
| *GDH2* | K15371 |
| *nrtA, nasF, cynA* | K15576 |

| | |
|---|---|
| *nrtB, nasE, cynB* | K15577 |
| *nrtC, nasD* | K15578 |
| *nrtD, cynD* | K15579 |
| *nirS* | K15864 |
| *nrfH* | K15876 |
| *CYP55* | K15877 |
| *NIT-6* | K17877 |
| *CA2* | K18245 |
| *CA4* | K18246 |
| *NAO* | K19823 |
| *K20932* | K20932 |
| *K20933* | K20933 |
| *K20934* | K20934 |
| *hdh* | K20935 |
| *vnfD* | K22896 |
| *vnfK* | K22897 |
| *vnfG* | K22898 |
| *vnfH* | K22899 |
| *nasE* | K26138 |
| *nasD, nasB* | K26139 |
| *nod* | K27148 |

**Table S5:** Sample overview for carrier-associated communities on MESIF and MUTAG™ matrices. Sampling days are given relative to the switch from ambient air to poultry-house exhaust air on day 0. “Before”, “initial”, and “later” denote temporal sampling windows rather than distinct aeration regimes (see Table S1).

| **Cultivation matrix** | Days relative to switch to exhaust air | Sampling window |
|---|---|---|
| MESIF | -7 | Before |
| | -4 | Before |
| | -3 | Before |
| | 0 | Before |
| | 3 | Initial |
| | 5 | Initial |
| | 10 | Initial |
| | 14 | Later |
| | 17 | Later |
| | 19 | Later |
| | 21 | Later |
| | 25 | Later |
| MUTAG™ | -7 | Before |
| | -4 | Before |
| | -3 | Before |
| | 0 | Before |
| | 3 | Initial |
| | 5 | Initial |
| | 10 | Initial |
| | 14 | Later |
| | 17 | Later |
| | 19 | Later |
| | 21 | Later |
| | 25 | Later |

**Table S6**: Unique MAGs from MESIF and MUTAG™ reconstructed and quality-checked using the described binning and QC pipelines.

| Matrix | MAG | Phylum | Family | Genus | Completeness [%] | Contamination [%] | Genome Size [Mbp] | GC Content [%] |
|---|---|---|---|---|---|---|---|---|
| MESIF | 1 | *Verrucomicrobiota* | *Coraliomargaritaceae* | *SKFT01* | 54.2 | 6.0 | 1.3 | 50 |
| | 2 | *Bacteroidota* | *Cyclobacteriaceae* | *Lunatimonas* | 74.9 | 6.5 | 5.3 | 40 |
| | 3 | *Verrucomicrobiota* | *DYNR01* | | 97.4 | 0.2 | 3.1 | 50 |
| | 4 | *Bacteroidota* | *Cyclobacteriaceae* | *Cecembia* | 51.0 | 5.5 | 2.5 | 40 |
| | 5 | *Pseudomonadota* | *Halomonadaceae* | *Billgrantia* | 81.5 | 3.0 | 3.4 | 60 |
| | 6 | *Pseudomonadota* | *UBA12402* | *1-14-0-20-46-22* | 79.8 | 0.9 | 1.2 | 40 |
| | 7 | *Pseudomonadota* | *Maricaulaceae* | *Oceanicaulis* | 99.7 | 5.7 | 2.9 | 60 |
| | 8 | *Pseudomonadota* | *Maricaulaceae* | *Glycocaulis* | 100 | 0.2 | 2.7 | 60 |
| | 9 | *Pseudomonadota* | *Inquilinaceae* | | 80.3 | 0.7 | 2.7 | 60 |
| | 10 | *Bacteroidota* | *Schleiferiaceae* | *CSBr16-58* | 100 | 0.0 | 2.5 | 40 |
| MUTAG™ | 1 | *Hydrogenedentota* | | | 92.8 | 4.0 | 4.1 | 60 |
| | 2 | *Actinomycetota* | *Microtrichaceae* | *SKRR01* | 95.5 | 0.0 | 3.7 | 60 |
| | 3 | *Bdellovibrionota* | | | 96.0 | 3.0 | 2.8 | 40 |
| | 4 | *Planctomycetota* | *Planctomycetaceae* | *Rubinisphaera* | 64.7 | 1.1 | 3.6 | 50 |
| | 5 | *Pseudomonadota* | *Rhizobiaceae* | *ZYF759* | 98.3 | 4.5 | 4.4 | 60 |
| | 6 | *Pseudomonadota* | *Legionellaceae* | *Legionella_A* | 99.4 | 3.1 | 2.0 | 30 |
| | 7 | *Myxococcota* | *Nannocystaceae* | | 93.7 | 0.5 | 9.1 | 60 |

**Table S7:** Shared MAGs recovered from both MESIF and MUTAG™ cultivation matrices. The table lists phylum, family, and genus assignments together with genome quality metrics, including completeness, contamination, genome size, and GC content.

| MAG | Phylum | Family | Genus | Completeness [%] | Contamination [%] | Genome Size [Mbp] | GC Content [%] |
|---|---|---|---|---|---|---|---|
| 1 | *Pseudomonadota* | *Alcanivoracaceae* | *Isoalcanivorax* | 85.8 | 5.0 | 3.2 | 60 |
| 2 | *Pseudomonadota* | *Algiphilaceae* | | 85.7 | 2.1 | 3.0 | 70 |
| 3 | *Pseudomonadota* | *Alteromonadaceae* | *Alkalimonas* | 100.0 | 0.3 | 3.9 | 50 |
| 4 | *Pseudomonadota* | *Alteromonadaceae* | *Aliidiomarina* | 89.0 | 0.6 | 2.5 | 50 |
| 5 | *Pseudomonadota* | *Beijerinckiaceae* | *Saliniramus* | 95.8 | 3.5 | 3.8 | 70 |
| 6 | *Bacteroidota* | *Cyclonatronaceae* | *Cyclonatronum* | 100.0 | 0.0 | 4.1 | 50 |
| 7 | *Bacteroidota* | *Cyclonatronaceae* | *SLDG01* | 67.9 | 5.1 | 2.4 | 40 |
| 8 | *Verrucomicrobiota* | *DYNR01* | | 97.5 | 0.2 | 3.2 | 60 |
| 9 | *Bacteroidota* | *Flavobacteriaceae* | *Flavobacterium* | 99.6 | 0.1 | 3.1 | 40 |
| 10 | *Pseudomonadota* | *Inquilinaceae* | | 80.3 | 0.8 | 2.7 | 70 |
| 11 | *Eremiobacterota* | *JADMJV01* | | 96.3 | 0.5 | 5.8 | 60 |
| 12 | *Hydrogenedentota* | *JAEUWI01* | | 91.2 | 0.9 | 4.3 | 70 |
| 13 | *Spirochaetota* | *Leptospiraceae* | *GCA-2786225* | 94.8 | 3.3 | 3.9 | 40 |
| 14 | *Actinomycetota* | *Microbacteriaceae* | *Pontimonas* | 93.0 | 3.2 | 1.6 | 60 |
| 15 | *Actinomycetota* | *Mycobacteriaceae* | *Dietzia* | 95.9 | 2.0 | 3.7 | 70 |
| 16 | *Pseudomonadota* | *Natronospirillaceae* | *Saccharospirillum* | 100.0 | 3.4 | 4.6 | 60 |
| 17 | *Pseudomonadota* | *Nitrosomonadaceae* | *Nitrosomonas* | 99.0 | 0.2 | 2.9 | 50 |

| | | | | | | | |
|---|---|---|---|---|---|---|---|
| 18 | *Pseudomonadota* | *Oleiphilaceae* | *JAIUMV01* | 99.3 | 0.0 | 4.0 | 60 |
| 19 | *Pseudomonadota* | *Oleiphilaceae* | *Marinobacter* | 95.1 | 0.2 | 3.8 | 60 |
| 20 | *Myxococcota* | *PHBI01* | *SYAK01* | 95.9 | 4.4 | 5.7 | 70 |
| 21 | *Planctomycetota* | *Pirellulaceae* | *Aureliella* | 99.6 | 1.4 | 6.3 | 50 |
| 22 | *Planctomycetota* | *Pirellulaceae* | *UBA6163* | 99.4 | 0.8 | 5.9 | 60 |
| 23 | *Pseudomonadota* | *Rhodobacteraceae* | *EhC02* | 100.0 | 2.2 | 4.9 | 60 |
| 24 | *Pseudomonadota* | *Rhodobacteraceae* | *Pararhodobacter* | 99.8 | 1.0 | 3.3 | 60 |
| 25 | *Pseudomonadota* | *Rhodobacteraceae* | *Rhabdonatronobacter* | 89.7 | 2.9 | 3.7 | 70 |
| 26 | *Spirochaetota* | *Salinispiraceae* | *SKVS01* | 78.3 | 6.9 | 2.7 | 40 |
| 27 | *Verrucomicrobiota* | *SKLO01* | *SKLO01* | 93.4 | 0.7 | 5.7 | 70 |
| 28 | *Planctomycetota* | *UBA1924* | | 90.5 | 0.5 | 3.4 | 70 |
| 29 | *Planctomycetota* | *UBA1924* | *SLFH01* | 61.5 | 0.5 | 2.4 | 70 |
| 30 | *Pseudomonadota* | *UBA2020* | *UBA2020* | 100.0 | 1.3 | 2.6 | 50 |
| 31 | *Verrucomicrobiota* | *UBA6053* | *PXAS01* | 92.1 | 0.8 | 4.2 | 60 |
| 32 | *Gemmatimonadota* | *UBA6960* | *SKNW01* | 90.8 | 2.5 | 3.8 | 70 |
| 33 | *Pseudomonadota* | *Wenzhouxiangellaceae* | *Wenzhouxiangella* | 100.0 | 1.2 | 4.0 | 60 |

**Table S8:** Detailed eggNOG-mapper annotations of denitrification-associated genes detected in carrier-derived MAGs**.** For each MAG, the putative target gene, eggNOG-mapper e-value and score, COG functional category, functional description, Enzyme Commission (EC) number, and KEGG Orthology (KO) identifier are reported. Genes screened included *narG*, *narZ*, *napA*, *nirK*, *nirS*, *norB*, *norC*, and *nosZ*. Multiple entries for the same gene and MAG represent distinct annotated proteins.

| MAG | Gene | evalue | score | COG | Description | EC | KO |
|---|---|---|---|---|---|---|---|
| MESIF_02 | *nosZ* | 0 | 910 | C | Nitrous oxide reductase | 1.7.2.4 | K00376 |
| MESIF_02 | *nosZ* | 4.7E-124 | 370 | C | Nitrous oxide reductase | 1.7.2.4 | K00376 |
| Shared_03 | *nosZ* | 0 | 1098 | C | Cytochrome c oxidase subunit | 1.7.2.4 | K00376 |
| MESIF_05 | *napA* | 0 | 1487 | C | Catalytic subunit of the nitrate reductase (NAP). Only expressed at high levels during aerobic growth. NapAB complex receives electrons from the membrane-anchored tetraheme protein NapC. Essential function for nitrate assimilation and may have a role in anaerobic metabolism | - | K02567 |
| MUTAG_5 | *napA* | 0 | 1388 | C | Belongs to the prokaryotic molybdopterin-containing oxidoreductase family | - | K00372, K02567 |
| MESIF_05 | *narG* | 1.76E-40 | 140 | C | Belongs to the prokaryotic molybdopterin-containing oxidoreductase family | 1.7.5.1 | K00370 |
| MESIF_05 | *narG* | 0 | 1733 | C | Belongs to the prokaryotic molybdopterin-containing oxidoreductase family | 1.7.5.1 | K00370 |
| Shared_05 | *narZ* | 0 | 2301 | C | Belongs to the prokaryotic molybdopterin-containing oxidoreductase family | 1.7.5.1 | K00370 |
| Shared_06 | *napA* | 7.84E-194 | 576 | PT | Sodium/hydrogen exchanger family | - | - |
| Shared_07 | *napA* | 3.07E-68 | 229 | PT | Sodium/hydrogen exchanger family | - | - |
| MESIF_09 | *nosZ* | 0 | 1059 | C | Nitrous-oxide reductase is part of a bacterial respiratory system which is activated under anaerobic conditions in the presence of nitrate or nitrous oxide | 1.7.2.4 | K00376 |
| Shared_09 | *nosZ* | 0 | 1219 | C | Nitrous oxide reductase | 1.7.2.4 | K00376 |
| Shared_10 | *nosZ* | 0 | 1059 | C | Nitrous-oxide reductase is part of a bacterial respiratory system which is activated under anaerobic conditions in the presence of nitrate or nitrous oxide | 1.7.2.4 | K00376 |
| Shared_11 | *napA* | 1.19E-177 | 529 | PT | Sodium/hydrogen exchanger family | - | - |
| Shared_12 | *napA* | 0 | 1155 | C | molybdopterin cofactor binding | - | K02567 |
| Shared_16 | *narG* | 0 | 2194 | C | the beta subunit is an iron sulfur cluster containing electron | 1.7.5.1 | K00370 |

| | | | | | | | |
|---|---|---|---|---|---|---|---|
| | | | | | transfer subunit | | |
| Shared_16 | *norB* | 0 | 986 | P | Belongs to the heme-copper respiratory oxidase family | 1.7.2.5 | K04561 |
| Shared_18 | *napA* | 0 | 1397 | C | Catalytic subunit of the nitrate reductase (NAP). Only expressed at high levels during aerobic growth. NapAB complex receives electrons from the membrane-anchored tetraheme protein NapC. Essential function for nitrate assimilation and may have a role in anaerobic metabolism | - | K02567 |
| Shared_18 | *narG* | 0 | 2541 | C | Belongs to the prokaryotic molybdopterin-containing oxidoreductase family | 1.7.5.1 | K00370 |
| Shared_18 | *nosZ* | 0 | 1094 | C | Cytochrome c oxidase subunit | 1.7.2.4 | K00376 |
| Shared_19 | *napA* | 0 | 1410 | C | nitrate reductase (NAP). Only expressed at high levels during aerobic growth. NapAB complex receives electrons from the membrane-anchored tetraheme protein NapC | - | K02567 |
| Shared_19 | *narG* | 0 | 2520 | C | Belongs to the prokaryotic molybdopterin-containing oxidoreductase family | 1.7.5.1 | K00370 |
| Shared_20 | *napA* | 0 | 1139 | C | molybdopterin cofactor binding | - | K02567 |
| Shared_23 | *narZ* | 0 | 1739 | C | Respiratory nitrate reductase alpha N-terminal | 1.7.5.1 | K00370 |
| Shared_23 | *nosZ* | 0 | 1045 | C | Nitrous-oxide reductase is part of a bacterial respiratory system which is activated under anaerobic conditions in the presence of nitrate or nitrous oxide | 1.7.2.4 | K00376 |
| Shared_23 | *nosZ* | 0 | 917 | C | Nitrous-oxide reductase is part of a bacterial respiratory system which is activated under anaerobic conditions in the presence of nitrate or nitrous oxide | 1.7.2.4 | K00376 |
| Shared_25 | *narZ* | 0 | 2292 | C | Respiratory nitrate reductase alpha N-terminal | 1.7.5.1 | K00370 |
| Shared_26 | *napA-2* | 9.89E-26 | 100 | P | Belongs to the Dps family | - | K04047 |
| Shared_26 | *napA-1* | 1.52E-246 | 703 | GPT | PFAM Sodium hydrogen exchanger family | - | - |
| Shared_26 | *napA-1* | 1.07E-37 | 132 | G | Phosphoenolpyruvate-dependent sugar phosphotransferase system, EIIA 2 | 2.7.1.2 02 | K02768, K02769, K02770, K02806 |
| Shared_26 | *napA-1* | 6.32E-73 | 243 | GPT | PFAM Sodium hydrogen exchanger family | - | - |
| Shared_26 | *napA-1* | 4.41E-148 | 444 | GPT | PFAM Sodium hydrogen exchanger family | - | - |
| Shared_28 | *napA* | 0 | 1147 | C | molybdopterin cofactor binding | - | K02567 |
| Shared_28 | *nosZ* | 3.86E-301 | 842 | C | Nitrous oxide reductase | 1.7.2.4 | K00376 |

| | | | | | | | |
|---|---|---|---|---|---|---|---|
| Shared_29 | *napA* | 5.38E-183 | 535 | C | molybdopterin cofactor binding | - | K02567 |
| Shared_29 | *nosZ* | 5.83E-266 | 753 | C | Nitrous oxide reductase | 1.7.2.4 | K00376 |
| Shared_31 | *napA-1* | 1.09E-164 | 498 | GPT | PFAM Sodium hydrogen exchanger family | - | - |
| Shared_31 | *napA-1* | 5.47E-76 | 239 | G | phosphoenolpyruvate-dependent sugar phosphotransferase system | 2.7.1.2 02 | K02768, K02769, K02770, K02806, K04752 |
| Shared_32 | *napA* | 9.67E-231 | 671 | PT | Sodium/hydrogen exchanger family | - | - |
| Shared_32 | *napA* | 5.81E-199 | 585 | PT | Sodium/hydrogen exchanger family | - | - |
| Shared_32 | *nosZ* | 8.07E-297 | 830 | C | nitrous-oxide reductase activity | 1.7.2.4, 1.9.3.1 | K00376, K02275 |
| Shared_33 | *narZ* | 0 | 1628 | C | Belongs to the prokaryotic molybdopterin-containing oxidoreductase family | 1.7.5.1 | K00370 |
| Shared_33 | *narZ* | 8.01E-134 | 414 | C | Belongs to the prokaryotic molybdopterin-containing oxidoreductase family | 1.7.5.1 | K00370 |
